# Single-nucleus transcriptome atlas of orbitofrontal cortex in amyotrophic lateral sclerosis with a deep learning-based decoding of alternative polyadenylation mechanisms

**DOI:** 10.1101/2023.12.22.573083

**Authors:** Paul M. McKeever, Aiden M. Sababi, Raghav Sharma, Zhiyu Xu, Shangxi Xiao, Philip McGoldrick, Troy Ketela, Christine Sato, Danielle Moreno, Naomi Visanji, Gabor G. Kovacs, Julia Keith, Lorne Zinman, Ekaterina Rogaeva, Hani Goodarzi, Gary D. Bader, Janice Robertson

**Author notes:** ^⋆^These authors contributed equally. Lead contacts and (Paul M. McKeever, Aiden M. Sababi, Gary D. Bader, Janice Robertson).

## Abstract

Amyotrophic lateral sclerosis (ALS) and frontotemporal lobar degeneration (FTLD) are two age-related and fatal neurodegenerative disorders that lie on a shared disease spectrum. While both disorders involve complex interactions between neuronal and glial cells, the specific cell-type alterations and their contributions to disease pathophysiology remain incompletely understood. Here, we applied single-nucleus RNA sequencing of the orbitofrontal cortex, a region affected in ALS-FTLD, to map cell-type specific transcriptional signatures in C9orf72-related ALS (with and without FTLD) and sporadic ALS cases. Our findings reveal disease- and cell-type-specific transcriptional changes, with neurons exhibiting the most pronounced alterations, primarily affecting mitochondrial function, protein homeostasis, and chromatin remodeling. A comparison with independent datasets from different cortical regions of C9orf72 and sporadic ALS cases showed concordance in several pathways, with neuronal STMN2 and NEFL showing consistent up-regulation between brain regions and disease subtypes. We also interrogated alternative polyadenylation (APA) as an additional layer of transcriptional regulation, demonstrating that APA events are not correlated with identified gene expression changes. To interpret these events, we developed APA-Net, a deep learning model that integrates transcript sequences with RNA-binding protein expression profiles, revealing cell type-specific patterns of APA regulation. Our atlas illuminates cell type-specific pathomechanisms of ALS/FTLD, providing a valuable resource for further investigation.

## Introduction

The advent of single-cell sequencing technology facilitates a deeper interrogation of the mechanisms underpinning disease, especially in complex tissues such as the brain. ALS is an adult-onset neurodegenerative disease caused by degeneration of motor neurons in the brain, brainstem, and spinal cord. Over 50% of ALS patients also exhibit progressive impairments in cognition, behaviour, or language caused by FTLD, with 10-15% of patients fulfilling the diagnostic criteria for frontotemporal dementia (FTD) ^1^. Although the vast majority of ALS cases are sporadic (sALS), with no family history of the disease, over 30 genes have been associated with disease causation, including SOD1, TARDBP (TDP-43), and FUS^2^. T he most common genetic cause linking ALS and FTLD is a G4C2 hexanucleotide repeat expansion within intron 1 of C9orf72 (C9) ^3,4^. This mutation leads to complex pathological mechanisms including dipeptide repeat protein toxicity, RNA-binding protein (RBP) sequestration by repeat-containing RNA, and reduced C9orf72 expression^3–5^. C9orf72 modulates the activity of small GTPases and has been linked with several membrane trafficking processes, including the autophagy lysosomal pathway, immune system regulation, synaptic plasticity and nucleocytoplasmic transport ^6–9^.

The molecular mechanisms underlying cortical dysfunction associated with FTLD in ALS remain elusive but have been linked to changes in expression and/or subcellular localizations of various RBPs. Nuclear depletion and cytoplasmic aggregation of TDP-43 is the hallmark pathology of motor neurons in ALS and cortical neurons in FTLD^10,11^. Other RBPs involved in ALS/FTLD include FUS ^12^, SFPQ ^13,14^, TIA1^15^, and heterogeneous nuclear ribonucleoproteins (hnRNP)s ^16,17^. RBPs regulate diverse functions across cell types, including transcription ^18–21^, auto-regulation^22,23^, alternative splicing (AS) ^19^, and APA ^23–27^. Disruption of RBP functions has been documented in ALS ^19,28^, with brain region-specific APA patterns observed in postmortem bulk RNA-seq from ALS patients ^29^ and in iPSC-derived motor neurons from patients with TARDBP and VCP mutations ^30^. However, the cell type-specific mechanisms underlying APA in ALS remain poorly understood.

Here, we generated a single-nucleus RNA sequencing (snRNA-seq) atlas comprising 103,076 nuclei to uncover transcriptomic changes in post-mortem orbitofrontal cortex from ALS caused by mutations in C9orf72 (C9-ALS) and sporadic ALS cases. We also analyzed C9-ALS cases with FTLD, identified by TDP-43 pathology in the frontal cortex (C9-ALS/FTLD). Our snRNA-seq analysis of the orbitofrontal cortex, revealed distinct molecular signatures in C9-ALS/FTLD, C9-ALS, and sALS cases. Each disease subtype showed unique patterns of pathway dysregulation across cell types, while also sharing common alterations in processes such as protein homeostasis and mitochondrial dysfunction. By integrating our findings with published datasets from the dorsolateral prefrontal and primary motor cortex^31–34^, we discovered concordant cell type-specific transcriptional changes across brain regions, suggesting these represent fundamental disease mechanisms rather than region-specific responses. These molecular signatures were particularly striking in neuronal populations, where we observed consistent patterns of gene expression changes across datasets.

We also investigated cell type-specific dysregulation of APA in ALS, identifying thousands of significant APA events in 3^′^ untranslated region (3^′^-UTR)s and upstream internal regions in C9-ALS and sALS compared to controls. To further understand this APA dysregulation, we developed an interpretable deep learning method called Alternative Polyadenylation Network (APA-Net). APA-Net enabled us to identify a range of cis and trans regulators correlated with the observed APA events in the disease. By interpreting APA-Net, we reveal potential RBP interactions in dysregulating APA in ALS, shedding light on the regulatory programs likely to induce APA in ALS. These findings enhance our understanding of cell-type specific transcriptomic changes occurring in the frontal cortex in ALS, and provide mechanistic insights into the potential coordinated interaction of RBPs in regulating APA in disease.

## Results

### Transcriptomic profiling of orbitofrontal cortex cell types in ALS patients with and without FTLD

We generated and analyzed a single cell atlas containing 103,076 nuclei derived from 23 snRNA-seq samples from orbitofrontal cortex (Brodmann area 11) of C9-ALS/FTLD (n=6 individuals), C9-ALS no FTLD (n=3), and sALS no FTLD (n=8) cases compared to controls (n=6) (Figure 1a, Supplementary Fig. 1a, Supplementary Data Table 1). Standard quality control (QC) steps were applied to the snRNA-seq data ^35^. We used an established snRNA-seq data integration method ^36^, ensuring consistent alignment of cell clusters across technologies for sex (Supplementary Fig. 1b), condition (Supplementary Fig. 1c), and samples (Supplementary Fig. 1d).

**Figure 1:**
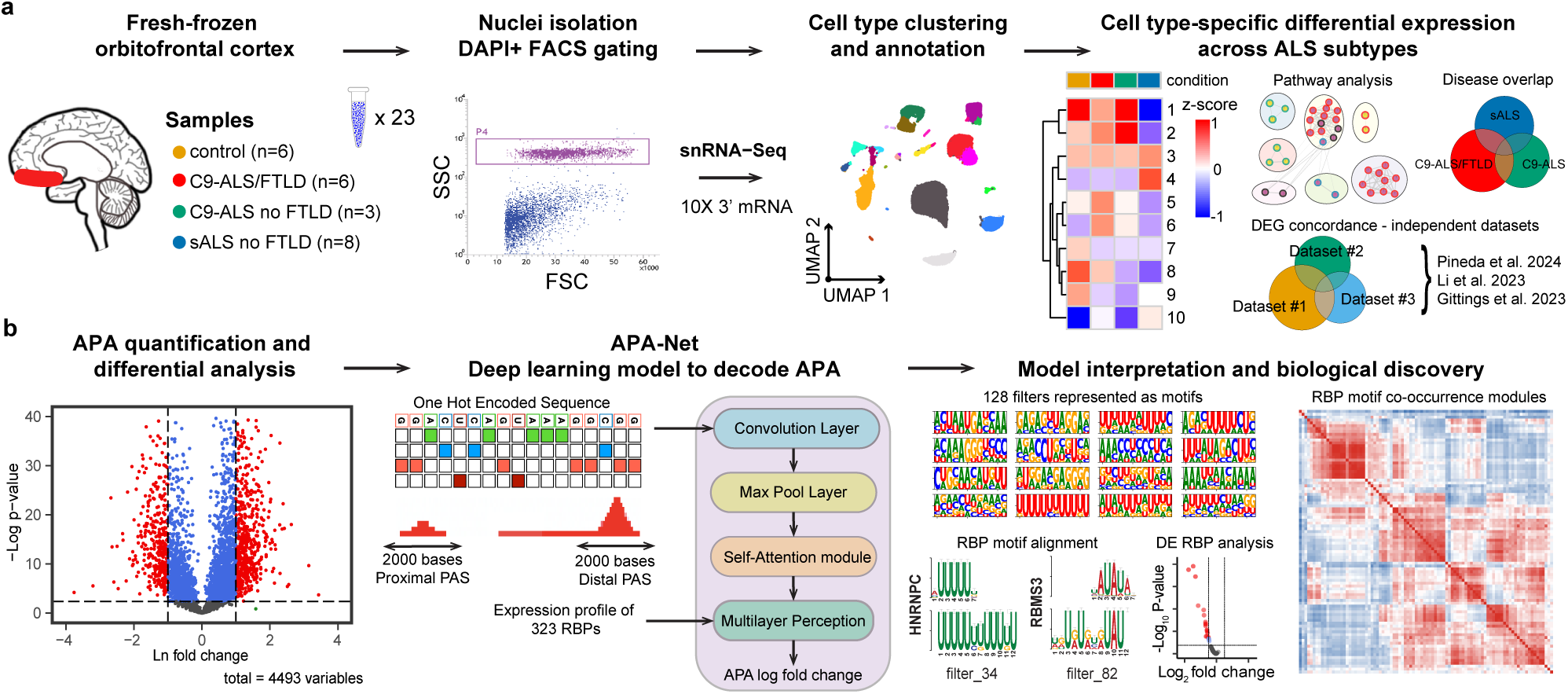
Single nucleus atlas for the study of transcriptome changes in ALS frontal cortex. **(a)** Workflow for the generation of a single nucleus atlas of the frontal cortex from controls (n=6 individuals), C9-ALS/FTLD (n=6), C9-ALS no FTLD (n=3), and sALS no FTLD (n=8). Nuclei were isolated using fluorescent activated cell sorting (FACS) nuclei labeled and gated with DAPI. Analyses include cell type clustering, machine learning and reference-based annotation, differential expression analysis, pathway analysis, overlap across ALS subtypes, and concordance with independent single nucleus datasets. **(b)** APA analysis across ALS subtypes and cell types. The APAlog software facilitated quantification and differential expression analysis of APA events from assigned reads. APA grammar was decoded using a multi input deep learning model called APA-Net which consists of a convolutional neural network with a multi-head attention mechanism. APA-Net uses the sequence information around polyadenylation site (PAS) and RBP expression profiles to predict APA log-fold change (LFC).

To delineate cell types, we applied an established clustering algorithm ^37^ and cell annotation approach ^38,39^ (Supplementary Fig. 1e,f, Methods). We uncovered 23 orbitofrontal cortex cell subtypes, including oligodendrocytes, oligodendrocyte precursor cells (OPC), astrocytes, endothelial cells, vascular leptomeningeal cells (VLMC)s, microglia and perivascular macrophages (PVM), ten excitatory neuron subtypes, and seven inhibitory neuron subtypes (Supplementary Fig. 1g). A catalogue of specific gene expression markers for each cell type is presented (Supplementary Fig. 1h). To facilitate downstream analysis, we grouped cell subtypes into nine major cell types: oligodendrocytes, OPCs, astrocytes, endothelial-VLMC, microglia-PVM, upper-layer excitatory neurons, intermediate-layer excitatory neurons, deep-layer excitatory neurons, and inhibitory neurons, based on canonical markers (Supplementary Fig. 2a-b). However, the nuclei from endothelial-VLMC were excluded from all downstream analyses due to poor yield across samples. All other cell types are detected across samples, showing similar distributions of cell types when categorized by disease subtype (Supplementary Fig. 2c-e).

### Converging and diverging transcriptomic changes across ALS subtypes

To uncover the transcriptomic cell-states in the orbitofrontal cortex of C9-ALS/FTLD, C9-ALS, and sALS cases compared to controls, we conducted a differential expression analysis^40^ (Methods). This analysis identified widespread differentially expressed genes across cell types in C9-ALS/FTLD, C9-ALS, and sALS relative to controls (false discovery rate (FDR) < 0.01; LFC (|log_2_ FC| < 0.5, Figure 2a-g, Supplementary Fig. 3, Supplementary Data Table 2). The majority of gene expression changes are found in excitatory and inhibitory neurons, with the most significant variances observed in C9-ALS/FTLD compared to C9-ALS and sALS (Figure 2a). Venn diagrams show the convergent and divergent changes among the three disease subtypes compared with controls for cortical layer excitatory neurons (Figure 2h-j), inhibitory neurons (Figure 2k), and glial cells: oligodendrocytes, OPCs, astrocytes, and microglia (Supplementary Fig. 6a-c, Supplementary Fig. 7a).

**Figure 2:**
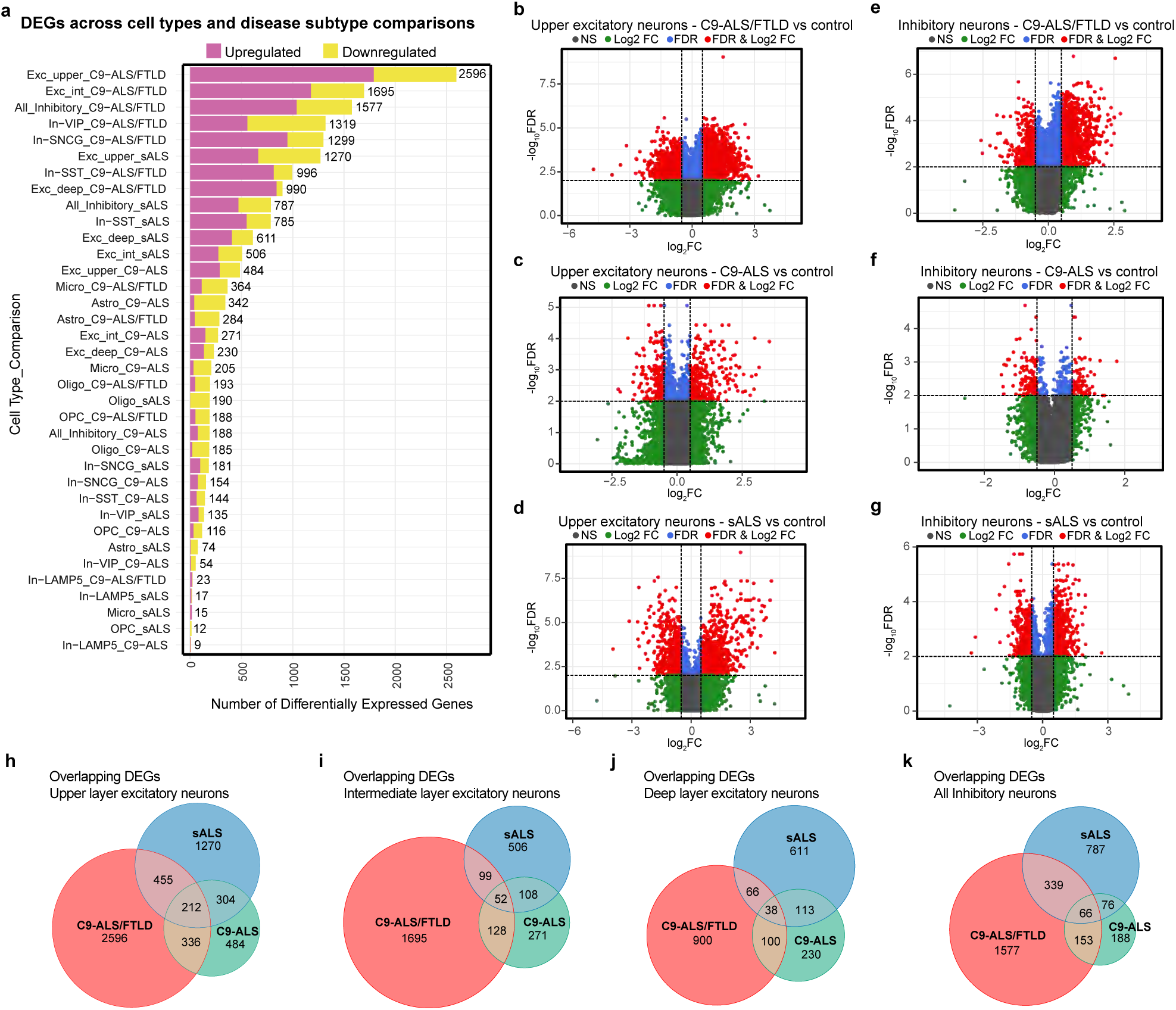
Distribution of differentially expressed genes across cell types in ALS. **(a)** Stacked bar plot indicating the number of differentially expressed genes on the y-axis across ALS cell sub-types. Purple indicates up-regulation and yellow is down-regulation. Volcano plots for differentially expressed genes in upper layer excitatory neurons in **(b)** C9-ALS/FTLD cases vs. controls; **(c)** C9-ALS vs. controls; **(d)** sALS vs. controls; and inhibitory neurons in **(e)** C9-ALS/FTLD cases vs. controls; **(f)** C9-ALS vs. controls; **(g)** sALS vs. controls. Only differentially expressed genes which passed the FDR < 0.01 and |LFC| > 0.5 cutoff were considered significant (shown in red). Venn diagram indicating overlapping differentially expressed genes between C9-ALS/FTLD, C9-ALS, and sALS in **(h)** upper layer excitatory neurons; **(i)** intermediate-layer excitatory neurons; **(j)** deep-layer excitatory neurons; **(k)** inhibitory neurons.

### Excitatory neurons in ALS are perturbed across all layers

To gain insights into the underlying biology of C9-ALS/FTLD, C9-ALS, and sALS, we conducted a pathway analysis ^41^ on differentially expressed genes across cell types (Supplementary Data Table 3). Upper-layer excitatory neurons across disease subtypes show up-regulation of genes associated with ribosomal complex and glutamatergic synapse, and down-regulation of chromatin remodeling, presynaptic signaling, and dendritic spine and postsynapse pathways ^42,43^ (Figure 3a). Major pathways changes in excitatory neurons across layers in C9-ALS/FTLD include up-regulation of the electron transport and respiratory chain, indicative of mitochondrial dysfunction (Figure 3a, Supplementary Fig. 4a,e). Other key pathways include those related to protein homeostasis, such as autophagy, the proteasome, and endoplasmic reticulum stress (Figure 3a, Supplementary Fig. 4a,e)

**Figure 3:**
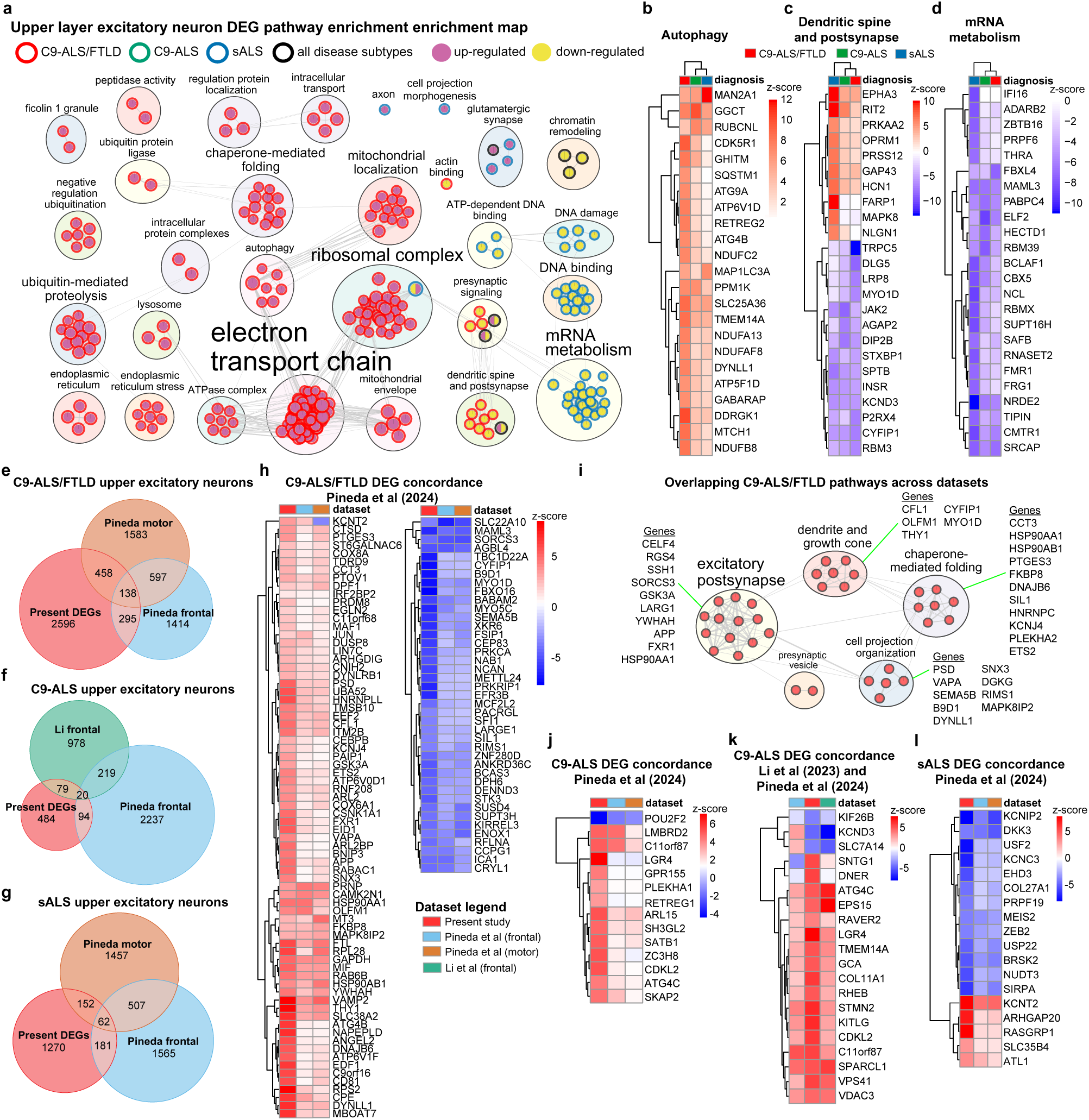
Upper layer excitatory neurons demonstrate converging and diverging changes across ALS disease subtypes. **(a)** Pathway analysis enrichment map for differentially expressed genes in upper layer excitatory neurons across ALS subtypes. Circles represent pathways, grouped by similarity (bubbles) and lines indicate shared genes between pathways. Circle centre and border colours indicate disease subtype and differential expression status. Clustered heatmap for genes identified in annotated enrichment map clusters represented by **(b)** autophagy; **(c)** dendritic spine and postsynapse; and **(d)** mRNA metabolism. **(e)** Venn diagram of C9-ALS/FTLD differentially expressed genes in upper excitatory neurons which overlap between the present study, and the motor and frontal cortices ^32^. **(f)** Venn diagram of C9-ALS differentially expressed genes in upper excitatory neurons which overlap between the present study and the frontal cortex from two independent studies ^31,32^. **(g)** Venn diagram of sALS differentially expressed genes in upper excitatory neurons which overlap between the present study, and the motor and frontal cortices ^32^. **(h)** Clustered heatmap of concordant C9-ALS/FTLD differentially expressed genes (positive correlation between z-scores), separated by up-regulated (left) and down-regulated (right) genes. **(i)** Enrichment map for overlapping biological processes enriched in C9-ALS/FTLD frontal and motor areas (based on 3-way intersection in **(e)**). Select genes highlighted from each cluster. **(j)** Clustered heatmap of concordant C9-ALS differentially expressed genes in frontal regions alone. **(k)** Clustered heatmap of concordant C9-ALS differentially expressed genes in frontal and motor regions. **(l)** Clustered heatmap of concordant sALS differentially expressed genes in frontal and motor regions. DEG = differentially expressed genes. Significance set at FDR < 0.01 and |LFC| > 0.5.

No specific pathways are observed for upper-layer excitatory neurons in C9-ALS, but up-regulation of postsynaptic organization (Supplementary Fig. 4a) and down-regulation of cerebral cortex migration (Supplementary Fig. 4e) are seen in intermediate and deep-layer excitatory neurons, respectively. In sALS, all cortical layer excitatory neurons show down-regulation of messenger RNA (mRNA) metabolism, whereas DNA damage and binding is altered in upper-layer neurons (Figure 3a, Supplementary Fig. 4a,e). Deep-layer excitatory neurons in sALS show specific up-regulation of glutamatergic and GABAergic synapses, axonogenesis, and forebrain development (Supplementary Fig. 4e), identifying many previously implicated axonal and potassium channel receptor genes^44,45^. Many convergent gene expression changes in excitatory neurons across layers showed concordance in the directionality of expression between disease subtypes, including genes associated with autophagy, neuronal function, mRNA metabolism, and the respiratory chain (Figure 3b-d, Supplementary Fig. 4b-d,f,g). This analysis revealed several genes directly relevant to the pathophysiology of ALS, including upregulation of SQSTM1, VCP, CHCHD10, UCHL1, and NEFL^2,46^.

To contextualize our findings, we next analyzed the overlap of differentially expressed genes from upper-layer excitatory neurons in our study with the same cells in two independent single nucleus datasets of dorsolateral prefrontal and primary motor cortex from C9-ALS, C9-FTLD, and sALS cases (Figure 3e-k, Supplementary Fig. 5a-d, Supplementary Data Table 4) ^31,32^. From this, we identify 138 overlapping differentially expressed genes between C9-ALS/FTLD cases from our data with frontal and motor cortex of C9-FTLD cases described in Pineda et al. (Figure 3e,h, Supplementary Fig. 5a). This overlap shows high concordance, with 115 genes showing the same directionality of change in expression (74 up- and 41 down-regulated; Figure 3h). Among concordant genes, PACRGL and ICA1, both known to contain cryptic exons repressed by TDP-43^47^, are down-regulated. Pathway changes related to overlapping differentially expressed genes include excitatory postsynapses, dendrites and axonal growth cones, chaperone-mediated folding, and cell projection organization (Figure 3i). These results suggest that despite regional differences, a core group of overlapping genes and pathways contribute to excitatory neuron perturbations in C9-ALS/FTLD.

In C9-ALS, we identify 20 overlapping differentially expressed genes between our dataset and those from the frontal cortex in Li et al. and Pineda et al. (Figure 3f), of which 16 of these show concordance (Figure 3j, Supplementary Fig. 5b,c). Notably, STMN2 shows consistent up-regulation across datasets in upper-layer excitatory neurons, a gene known to undergo cryptic splicing and premature polyadenylation under TDP-43 deficiency in ALS ^25,47–49^. C9-ALS cases also consistently show up-regulation of ATG4C, C11orf87, and CDKL12 across all regions and datasets (Supplementary Fig. 5c). These genes are not characterized in the context of ALS but are highly expressed in excitatory and inhibitory neurons ^50^.

For sALS, we identify 62 overlapping differentially expressed genes between our study and Pineda et al.’s motor and frontal cortex datasets (Figure 3g). Concordant expression patterns across datasets include dysregulation of 18 genes involving potassium channels (KCNT2, KCNIP2, KCNC3), neuronal differentiation (BRSK2, ATL1, ARHGAP20), and neuronal stress/structural markers (USF2, EHD3, PRPF19, ZEB2, USP22, SIRPA), of which most are down-regulated (Figure 3k). The consistent dysregulation of genes involved in potassium channels, neuronal differentiation, and stress response across multiple independent datasets highlights common molecular mechanisms of excitatory neuron vulnerability in both sporadic and C9-linked ALS.

### Inhibitory neurons in ALS show mitochondrial dysfunction across ALS subtypes

A pathway analysis of gene expression changes in inhibitory neurons revealed unique and overlapping alterations across ALS subtypes (Figure 4a). The most up-regulated pathway across disease subtypes is the electron transport chain, strongly implicating changes in mitochondrial function as a significant contributor to disease pathophysiology. Chromatin remodeling is down-regulated, representing a consistent change between both inhibitory and excitatory neurons across disease subtypes. Specific pathway changes in C9-ALS/FTLD include up-regulation of ribosomal subunits, RNA localization, mitochondrial function, and proteostasis (Figure 4a). C9-ALS shows up-regulation in axonal growth cone and down-regulation in double-strand DNA break repair pathways (Figure 4a). sALS is characterized by down-regulation in presynaptic signaling, dendritic structure, mRNA processing, histone binding, and DNA damage response pathways (Figure 4a). Analysis of top-ranked genes in RNA localization, double-strand DNA break repair, and chromatin remodeling show convergence between disease subtypes, including up-regulation of disease-relevant genes SOD1 and SQSTM1 (Figure 4b-d).

**Figure 4:**
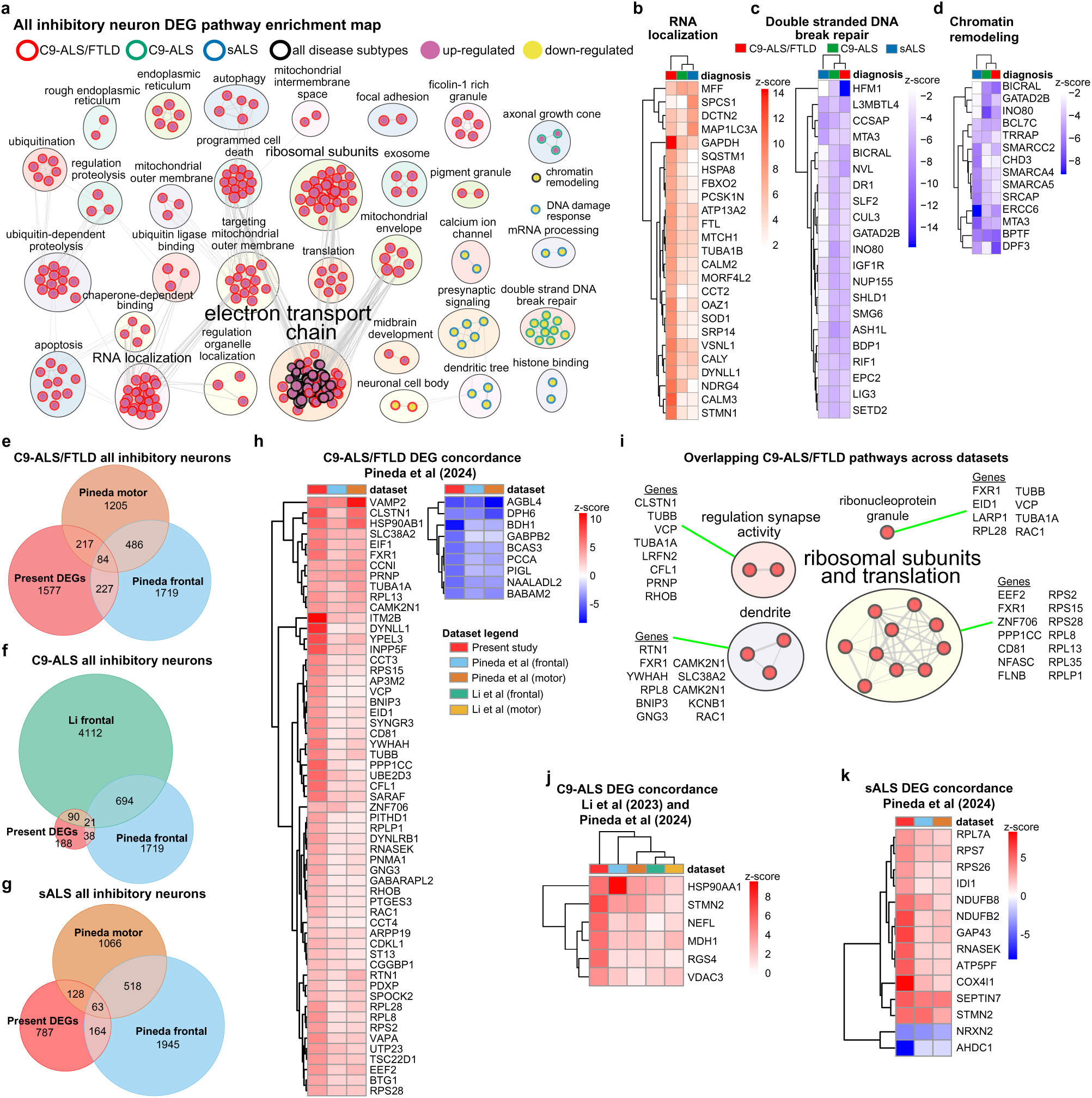
Inhibitory neurons show converging and diverging changes across ALS disease subtypes. **(a)** Pathway analysis enrichment map for differentially expressed genes in inhibitory neurons across ALS subtypes. Clustered heatmap for genes identified in annotated enrichment map clusters represented by **(b)** RNA localization; **(c)** double-stranded DNA break repair; and **(d)** chromatin remodeling. **(e)** Venn diagram of C9-ALS/FTLD differentially expressed genes in inhibitory neurons which overlap between the present study, and the motor and frontal cortices ^32^. **(f)** Venn diagram of C9-ALS differentially expressed genes in inhibitory neurons which overlap between the present study and the frontal cortex from two independent dataset ^31,32^. **(g)** Venn diagram of sALS differentially expressed genes in inhibitory neurons which overlap between the present study, and the motor and frontal cortices from an independent dataset ^32^. **(h)** Clustered heatmap of concordant C9-ALS/FTLD differentially expressed genes (positive correlation between z-scores), separated by up-regulated (left) and down-regulated (right) genes. **(i)** Enrichment map for overlapping biological processes enriched in C9-ALS/FTLD frontal and motor areas (based on 3-way intersection in **(e)**). Select genes highlighted from each cluster. **(j)** Clustered heatmap of concordant C9-ALS differentially expressed genes in frontal regions alone. **(k)** Clustered heatmap of concordant sALS differentially expressed genes in frontal and motor cortex regions ^32^. Significance set at FDR < 0.01 and |LFC| < 0.5. DEG = differentially expressed genes.

We further compared our differentially expressed genes in inhibitory neurons with the independent datasets (Figure 4e-j, Supplementary Data Table 4) ^31–33^. In C9-ALS/FTLD, 84 differentially expressed genes overlap with our study and the frontal and motor cortex from Pineda et al., with 66 concordant genes across datasets (57 up-regulated, 9 down-regulated) (Figure 4e,h, Supplementary Fig. 5e). These genes include up-regulation of ALS-related gene VCP^51^. Pathway changes related to overlapping differentially expressed genes include synaptic regulation, dendrite, ribonucleoprotein granules, and translational machinery (Figure 4i) ^32,34^. In C9-ALS, 21 genes overlap with our data and the frontal cortex from Li et al. and Pineda et al. (Figure 4f, Supplementary Fig. 5f). Among these, HSP90AA1, STMN2, NEFL, MDH1, RGS4, and VDAC3 were concordantly up-regulated across datasets (Figure 4i)^31,32^.

In sALS, 63 differentially expressed genes overlap between our study and the frontal and motor cortex from Pineda et al. (Figure 4g, Supplementary Fig. 5g). Of these, 12 genes are concordantly up-regulated, including RPL7A, RPS7, and RPS26, and STMN2, whereas two genes are down-regulated (Figure 4j). This convergent dysregulation of ribosomal genes and STMN2 across cortical regions in both C9-ALS and sALS points to shared molecular vulnerabilities in inhibitory neurons.

Table 1 summarizes examples of concordant differentially expressed genes across the compared studies for both upper-layer excitatory and inhibitory neurons. These findings highlight both converging and distinct molecular changes in inhibitory neurons across frontal and motor regions in ALS, with significant overlap with dysregulated genes in excitatory neurons (Figure 3, Figure 4, Supplementary Fig. 4).

**Table 1:** Examples of overlapping differentially expressed genes and their functions across independent datasets in ALS and FTLD ^31–34^.

| Disease | Gene | Function | Change | Cell |
| --- | --- | --- | --- | --- |
| C9-ALS/FTLD | CFL1<sup>9</sup> | Actin filament dynamics | up | Exc/Inh |
|  | RAC1<sup>9</sup> | Small GTPase | up | Inh |
|  | YWHAH<sup>54</sup> | 14-3-3 protein | up | Exc |
|  | KCNJ4<sup>55</sup> | Potassium channel | up | Exc |
|  | FXR1<sup>56</sup> | RNA-binding protein | up | Exc/Inh |
|  | APP<sup>57</sup> | neuroplasticity, neuroprotection | up | Exc |
|  | CAMK2N1<sup>58</sup> | Kinase inhibitor | up | Exc |
|  | HSP90AA1<sup>32</sup> | Heat shock protein | up | Exc/Inh |
|  | HSP90AB1<sup>32</sup> | Heat shock protein | up | Exc |
|  | DNAJB6 | Co-chaperone | up | Exc |
|  | FTL<sup>59</sup> | Ferritin light chain | up | Exc |
|  | MYO1D | Motor protein | down | Exc |
|  | ENOX1<sup>60</sup> | Oxidoreductase | down | Exc |
|  | ICA1<sup>47</sup> | ARF1 interactor in vesicle budding | down | Exc |
|  | PACRGL<sup>47</sup> | Microtubule binding | down | Exc |
|  | ATP5PF | ATP synthesis | up | Inh |
|  | NDUFBB | Electron transport | up | Inh |
|  | NDUFB2 | Electron transport | up | Inh |
|  | COX4I1 | Electron transport | up | Inh |
|  | EEF2 | Translation elongation factor | up | Inh |
|  | EIF2 | Translation initiation factor | up | Inh |
|  | PPP1CC | Protein phosphatase | up | Inh |
| C9-ALS | PLEKHA1<sup>61</sup> | Lipid binding | up | Exc |
|  | ATG4C | Autophagy | up | Exc |
|  | GABARAPL2 | Autophagy | up | Inh |
|  | RAVER2 | RNA-binding protein | up | Exc |
|  | SPARCL1 | Extracellular matrix protein | up | Exc |
|  | VDAC3 | Mitochondrial porin | up | Exc |
|  | RAC1<sup>9</sup> | Small GTPase | up | Inh |
|  | VCP<sup>51</sup> | ATPase | up | Inh |
|  | PRNP | Prion protein | up | Inh |
|  | TUBA1A | Tubulin | up | Inh |
|  | TUBB | Tubulin | up | Inh |
|  | GNG3 | G protein subunit | up | Inh |
| sALS | KCNIP2<sup>44,45</sup> | Potassium channel interactor | down | Exc |
|  | KCNC3<sup>44,45</sup> | Voltage-gated potassium channel | down | Exc |
|  | KCNT2<sup>44,45</sup> | Potassium channel | up | Exc |
|  | USP22 | Deubiquitinating enzyme | down | Exc |
|  | ARHGAP20 | Rho GTPase-activating protein | up | Exc |
|  | SLC35B4 | Nucleotide sugar transporter | up | Exc |
|  | ATL1 | Endoplasmic reticulum morphology | up | Exc |
| Across ALS | STMN2<sup>47,49</sup> | Microtubule dynamics | up | Exc/Inh |
|  | RGS4 | Regulator of G protein signaling | up | Inh |
|  | NEFL<sup>59</sup> | Neurofilament protein | up | Inh |
|  | MDH1 | Malate dehydrogenase | up | Inh |
|  | SLC38A2 | Glutamine transporter | up | Inh |

**Abbreviations:** Exc - Excitatory, Inh - Inhibitory

### Protein homeostasis pathways are dysregulated in glial cells across ALS subtypes

Oligodendrocytes in ALS may contribute to neuronal dysfunction through poor myelin maintenance, loss of metabolic support, and release of injurious pro-inflammatory signals^34,52^. Across ALS subtypes, oligodendrocytes show up-regulation of chaperone-mediated folding, while mRNA metabolism, and chromatin acetylation and remodeling are down-regulated (Supplementary Fig. 6a,d-f). This indicates broad disruptions in protein homeostasis and gene expression regulation in oligodendrocytes in ALS. In C9-ALS/FTLD, oligodendrocytes show up-regulation in apoptosis and down-regulation in nervous system development and neuron projection pathways. C9-ALS oligodendrocytes exhibit a depletion in double-stranded DNA break repair, with no specific pathways showing up-regulation. Although sALS oligodendrocytes do not show unique pathway alterations, TDP-43 is down-regulated (Supplementary Fig. 6d,f, Supplementary Data Table 2) ^53^. Several of the top-ranked genes for chaperone-mediated folding and mRNA metabolism show convergence across disease subtypes (Supplementary Fig. 6e,f).

In comparison with oligodendrocytes from frontal and motor cortex from Pineda et al., both C9-ALS/FTLD and sALS show down-regulation of SEPTIN4 (Supplementary Fig. 6g,i), a scaffold protein involved in myelination^62^. In C9-ALS/FTLD, there is also down-regulation of NAALADL2 (Supplementary Fig. 6g), which is decreased in oligodendrocytes from myelin-deficient mice and in the striatum of Huntington’s disease ^63^. Additionally, BAZ2B, which is involved in chromatin remodeling, and SRRM1, an RBP involved in splicing, are decreased (Supplementary Fig. 6g). Concordant with oligodendrocytes from independent frontal cortex datasets, C9-ALS shows up-regulation of ATG4B and HSP90AB1 (Supplementary Fig. 6h), both of which are related to autophagy and are targeted by TDP-43^64,65^. Oligodendrocytes from sALS (Supplementary Fig. 6i) show down-regulation of PLLP, a component of the myelin sheath, and CSNK1E, which phosphorylates TDP-43^66^.

In OPCs, ATP metabolism is up-regulated across disease subtypes (Supplementary Fig. 6j,k), suggesting a stressed cellular state. Chaperone-mediated folding is up-regulated in C9-ALS/FTLD and C9-ALS, whereas C9-ALS and sALS show down-regulation of GPI anchor processes and chromatin remodeling (Supplementary Fig. 6j). Specific C9-ALS/FTLD pathways include up-regulation of ubiquitin protein ligase activity and down-regulation in pathways involving regulation of cell morphogenesis/projection and tubulin binding (Supplementary Fig. 6j). OPCs from C9-ALS cases show up-regulation in endogenous hormone response and down-regulation in ATP-dependent DNA binding. In sALS, OPCs show down-regulation of synapse organization, potentially reflecting compromised neuronal connectivity. Several top-ranked genes in ATP metabolism and chromatin remodeling pathways demonstrate concordance across disease subtypes, highlighting areas of shared dysregulation (Supplementary Fig. 6k,l).

Despite limited overlap with independent datasets from Li et al and Pineda et al (Supplementary Data Table 4), two genes show disease-specific alterations in OPCs: in C9-ALS, ANKRD10 is concordantly up-regulated, aligning with its known up-regulation in skeletal muscle in ALS^67^. In sALS, APOD, a lipid-binding protein critical for remyelination, is consistently down-regulated, aligning with altered proteomic levels of APOD in cerebrospinal fluid from ALS patients^68,69^.

Astrocytes are linked with the pathophysiology of ALS through loss of homeostasis and reactive toxicity ^70,71^. Differential expression analysis reveals that astrocytes in C9-ALS/FTLD and C9-ALS exhibit significantly greater dysregulation compared to sALS astrocytes (Figure 2a, Supplementary Fig. 6c). Across ALS subtypes we see enrichment of pathways associated with the unfolded protein response and actin filament bundle formation (Supplementary Fig. 6m,n), whereas genes associated with chromatin organization are depleted (Supplementary Fig. 6m-o). In C9-ALS/FTLD, astrocytes show up-regulation in cholesterol and miRNA metabolism, with down-regulation of filopodium tip genes (Supplementary Fig. 6m). There is also up-regulation of GFAP and CHI3L1 (Supplementary Data Table 2), common markers of reactive astrocytes ^72,73^. For C9-ALS, astrocytes show up-regulation in protein localization and down-regulation in cilium assembly formation^74^ (Supplementary Fig. 6m). In sALS, astrocytes show down-regulation in nuclear protein complexes and TDP-43 (Supplementary Fig. 6m,o), supporting that a reactive astrocyte phenotype is associated with reduced expression of TDP-43^75^. An analysis of unfolded protein response and chromatin organization reveals convergence of gene expression changes in astrocytes across disease subtypes (Supplementary Fig. 6n,o).

A comparison with independent frontal and motor cortex datasets reveals concordant down-regulation of SLC4A4 and GRIA2 in C9-ALS/FTLD (Supplementary Fig. 6p, Supplementary Data Table 4), suggesting disruption of tripartite synapse signaling. In C9-ALS frontal cortex astrocytes, NTRK2, a receptor essential for astrocyte morphology and neuronal communication through BDNF signaling ^76,77^, is down-regulated (Supplementary Fig. 6q, Supplementary Data Table 4) ^70,71^. In sALS astrocytes, four concordant down-regulated genes are observed, including SNRNP70, an RBP with disrupted splicing in ALS ^78^ and interacts with ALS-related genes such as FUS^79^. Together, these findings highlight unique and overlapping astrocytic alterations across brain regions implicated in ALS and FTLD.

Altered cellular states in microglia, the primary immune cells of the central nervous system, are a feature of ALS pathogenesis, and in neurodegenerative disease more broadly^80–82^. Gene expression changes in microglia are more prominent in C9-ALS (with and without FTLD) versus sALS (Supplementary Fig. 7a-e), suggesting a strong link of C9orf72 to the immune system ^83,84^. Across all ALS subtypes, microglia show up-regulation in glucose metabolism, cytokine signaling, and chaperone-mediated refolding with down-regulation in small GTPase signaling, neuron projection morphogenesis, and the interleukin-6 pathway (Supplementary Fig. 7b). In C9-ALS/FTLD, microglia show up-regulation in pathways involved in innate immune response, antigen presentation and phagocytosis, endocytosis, exosome activity, and iron ion homeostasis (Supplementary Fig. 7b). Conversely, there is down-regulation in positive regulation of chemotaxis, glutamatergic synapse function, and negative regulation of biosynthesis (Supplementary Fig. 7b,c). In microglia in C9-ALS, we uncover a shift of these cells toward pro-inflammatory signaling, programmed cell death, and growth pathways (Supplementary Fig. 7b). Pathways showing gene expression convergence across disease subtypes include neuron projection morphogenesis, cytokine signaling, and interleukin-6 pathway (Supplementary Fig. 7c-d). For instance, JAK1 and JAK2 are down-regulated across all ALS subtypes (Supplementary Fig. 7e)^85^.

Orbitofrontal cortex microglia show concordance with independent published microglial data from ALS frontal and motor cortex^31–33^. For example, there is up-regulation of SYTL3 (Supplementary Fig. 7f) and down-regulation of ATP8B4 (Supplementary Fig. 7f), both of which are enriched in microglia ^50^. Rare variants in ATP8B4 are risk factors for Alzheimer’s disease ^86^. We examined microglial states across ALS subtypes by scoring gene expression ^87^ across four independent datasets ^31–33^ using established neurodegenerative disease markers ^81^. In three frontal cortex datasets, C9-ALS and C9-FTLD subtypes show reduced expression of homeostatic microglial markers, including CSF1R, CX3CR1, and P2RY12 (Supplementary Fig. 8a)^31,32^. Both C9-ALS and C9-FTLD subtypes show elevated disease-associated microglia (DAM) stage 1 markers (TYROBP, APOE, B2M)(Supplementary Fig. 8b), while SPP1 was the primary DAM stage 2 marker with increased expression (Supplementary Fig. 7g, Supplementary Fig. 8c). This shift to disease-associated states mirrors patterns seen in other neurodegenerative disease ^80–82^.

Across all analyzed cell types and ALS subtypes, we identified consistent enrichment of protein homeostasis pathways, including autophagy, chaperone-mediated folding, unfolded protein response, and ubiquitin-mediated proteolysis (Figure 3, Figure 4, Supplementary Fig. 4, Supplementary Fig. 6, and Supplementary Fig. 7). This pattern was corroborated through comparative analysis with independent ALS datasets, which confirmed up-regulation of chaperone-mediated folding genes, suggesting a common cellular response to preotein homeostasis disruption in ALS.

### ALS-related risk gene expression across frontal cortex cell types

To investigate the cell-type-specific expression of ALS-related genes, we analyzed gene expression patterns for 30 genes implicated in familial ALS or linked to heightened risk in sporadic cases^88,89^. Hierarchical clustering of ALS-related genes across cell types (Figure 5a) uncovered modules of microglial-enriched genes (Modules 1-3), oligodendrocytes, OPCs, and astrocyte-enriched (Modules 4-5), and neuronal-enriched (Modules 6-7). We next identified differentially expressed ALS-related genes in a cell type-specific manner (Figure 5b). Up-regulated ALS-related genes across disease subtypes include neuron-enriched CHCHD10, TUBA4A, and SOD1, in both C9-ALS/FTLD and sALS. Additionally, SQSTM1 is up-regulated in both neurons and glia in C9-ALS/FTLD and C9-ALS, while VCP shows specific up-regulation in neurons from C9-ALS/FTLD alone. The up-regulation of mitochondrial-associated genes CHCHD10, TUBA4A, SQSTM1, SOD1, and VCP in both excitatory and inhibitory neurons of C9-ALS/FTLD patients reinforces the mitochondrial dysfunction pathways identified through differential expression analysis (Figure 5b). CCNF up-regulation is specific to upper-layer excitatory neurons from C9-ALS. Down-regulation of ALS-related genes occurred in specific cell types (Figure 5b). TARDBP (TDP-43) is down-regulated in oligodendrocytes, astrocytes, and excitatory neurons in sALS. C9orf72 is down-regulated in oligodendrocytes from C9 cases, while FUS shows down-regulation across all excitatory neuron subtypes. SETX is down-regulated in upper-layer neurons in C9-ALS/FTLD. In sALS, ANXA11 is down-regulated in upper-layer excitatory neurons and VIP+ inhibitory neurons. These observations highlight the differential regulation of ALS-related genes across cell types and disease subtypes, providing molecular insights into the mechanisms underlying ALS pathogenesis.

**Figure 5:**
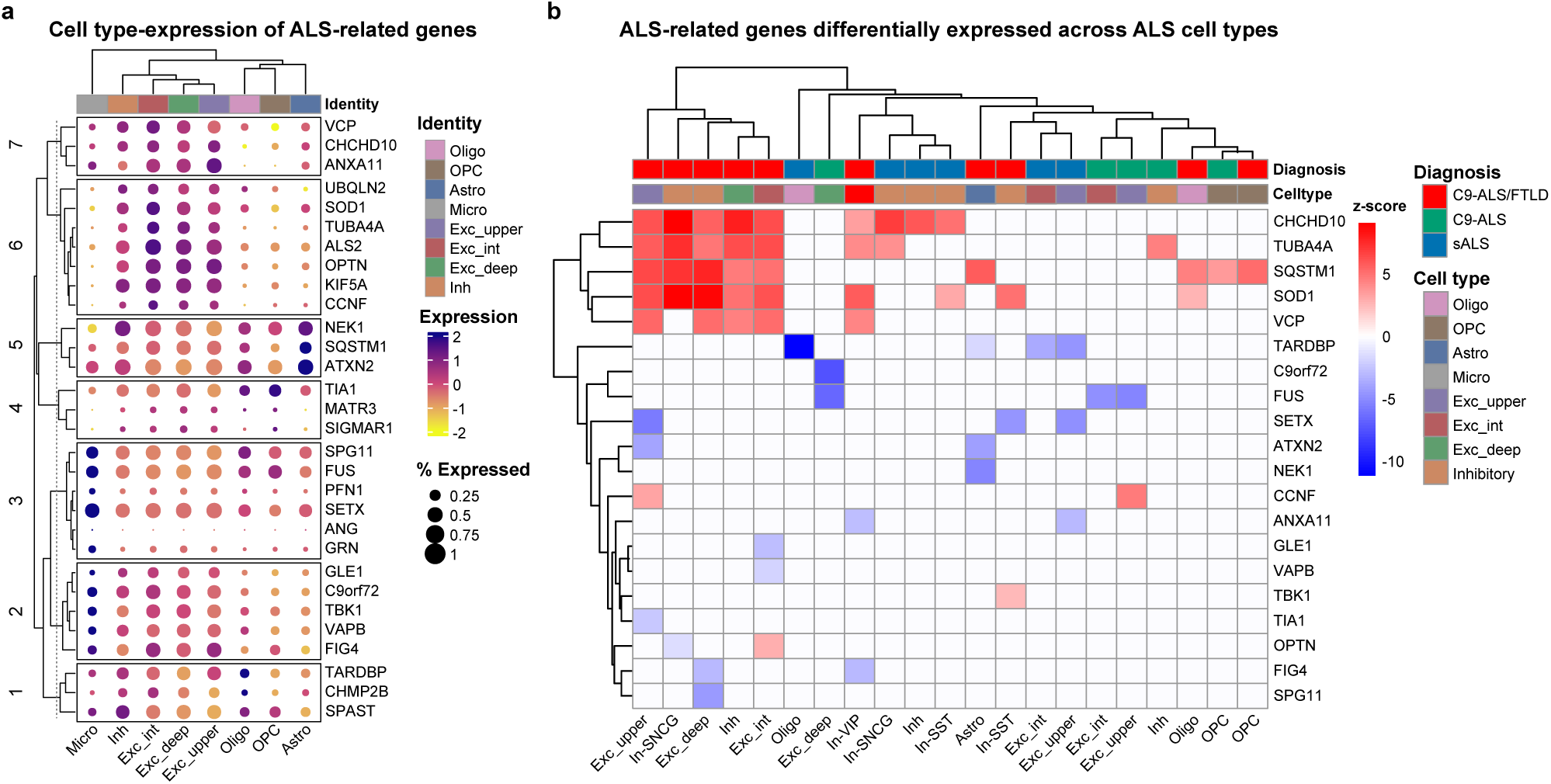
Distribution of differentially expressed ALS-related genes across orbitofrontal cell types in ALS. **(a)** Clustered heatmap dot plot analysis comparing average ALS-related gene expression levels (y-axis) in cell types. The size of dot corresponds to the percent of cells of a given cell type expressing the corresponding gene of interest. Cell type identity indicated in the upper x-axis by colour coding and lower x-axis by label. Columns are clustered hierarchically, whereas rows are clustered and partitioned with k-means clustering (7 clusters). **(b)** Clustered heatmap of ALS-related genes intersecting with differentially expressed genes across cell types and ALS subtypes. Scale is in z-scores, where the red scale indicates up-regulation, the blue scale shows down-regulation, and the white shows the gene was unchanged in a particular cell type for a given disease condition. Significance set at FDR < 0.05 and |LFC| < 0.5. DEG = differentially expressed genes.

### Dysregulation of APA landscape in ALS

While previous studies have demonstrated APA dysregulation in neurodegeneration ^90^ and ALS ^18,29,91^, the underlying mechanisms at the cell type level have remained largely unexplored. We identified known polyadenylation (PA) sites in transcripts across our ALS subtypes and controls and performed differential PA analysis to compare ALS with controls (Supplementary Data Table 5) ^92,93^ (Methods). To mitigate false positives, we conducted an empirical FDR analysis using the control samples to establish cell type-specific FDR thresholds (Supplementary Data Table 6). These stringent criteria enabled us to filter out potential false positives across different cell types, ensuring a FDR of 10% or less for significant APA events. To further validate the APA findings and rule out potential confounding factors, we investigated the relationship between APA events and differentially expressed genes. We examined whether genes exhibiting APA, either lengthening, shortening, or intronic, also show significant changes in overall expression levels across the cell types analyzed. Our analysis reveals no significant correlation between differentially expressed genes and APA events (Supplementary Fig. 9a-e), consistent with a recent study showing the independence of differentially expressed genes and APA ^94^. This also suggests that APA events represent a distinct regulatory layer, potentially influencing disease-specific cellular functions without altering overall gene expression levels.

Our differential analysis revealed numerous significant PA differences between ALS subtypes and controls across all major cell types (Figure 6a,b). In both C9-ALS (with and without FTLD) and sALS, we observe a global trend towards distal PA site usage (Figure 6a,b). However, we also identified a considerable number of intronic and shortened genes, indicating that APA occurs in both directions. This pattern is evident across major cell types in both C9-ALS and sALS, but a higher proportion of intronic APA usage is detected in all neuron subtypes, oligodendrocytes, microglia, and astrocytes in sALS compared to C9-ALS (Figure 6a,b). Intronic APA usage is particularly prevalent in intermediate- and deep-layer excitatory neurons of sALS cases. The abundance of 3^′^-UTR shortened and intronic APAs in ALS compared to controls suggests a potential increase in truncated gene transcripts, which could significantly impact gene function. Conversely, the shift towards distal PA sites and longer 3^′^-UTR could affect function by altering mRNA metabolism, transport, and stability^95,96^.

**Figure 6:**
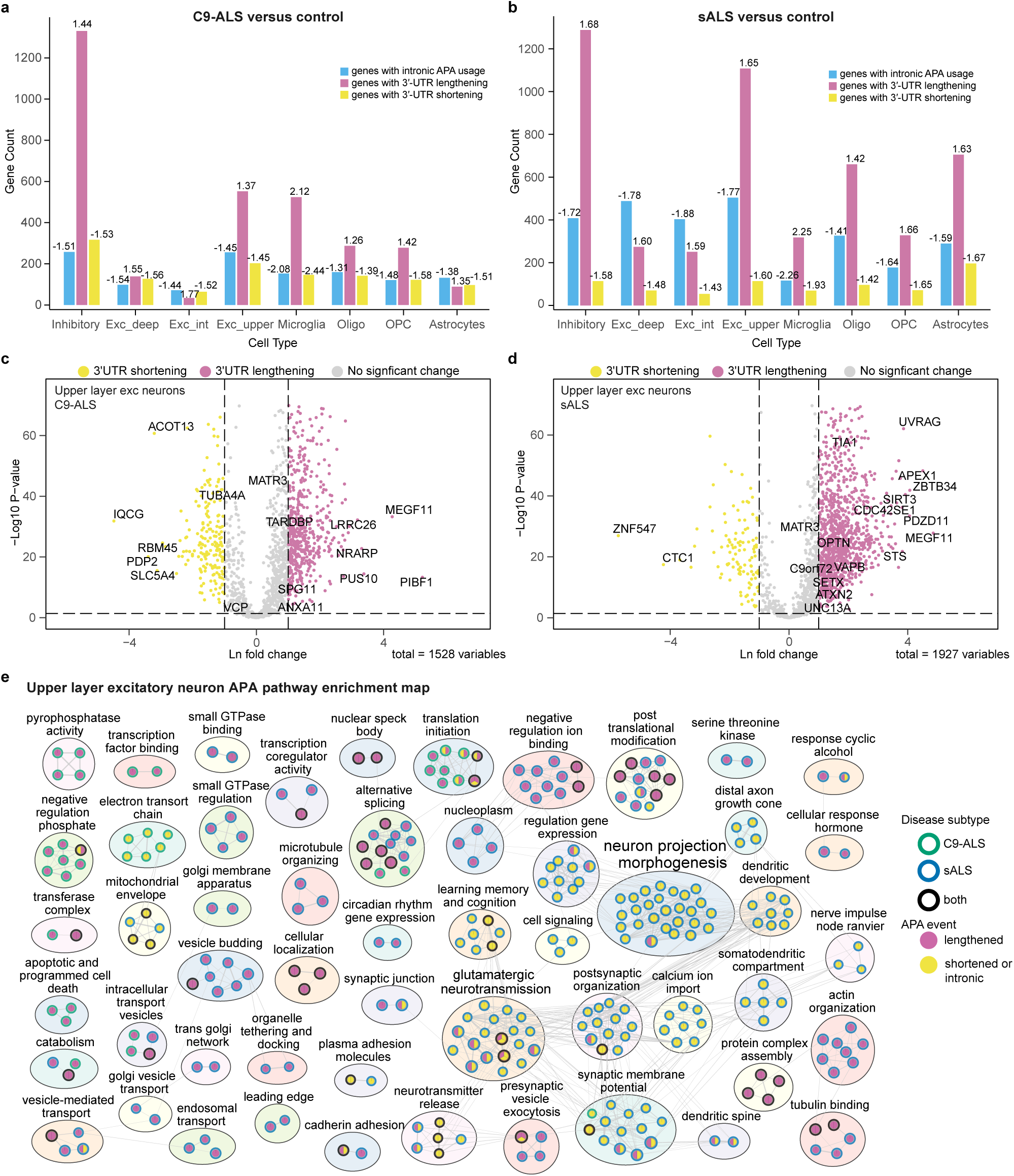
Dysregulation of APA landscape in ALS compared to control. **(a)** Barplot depicting the number of genes undergoing 3*^′^*-UTR lengthening, shortening, and intronic PA site usage in C9-ALS compared to control, across major cell types. The median REDu and REDi scores are shown above each bar. **(b)** Same as (a) for sALS. **(c)** Volcano plot for upper layer excitatory neurons in C9-ALS for genes undergoing 3*^′^*-UTR lengthening and shortening. Only genes that passed the 10% empirical FDR threshold for transcript-wise APA usage are shown (see Methods). **(d)** Same as (c) for sALS. **(e)** Enrichment map showcasing the pathways enriched for genes undergoing lengthening and shortening in C9-ALS and sALS for upper layer excitatory neurons. The node border indicates which ALS subtype is enriched, whereas the fill shows whether the pathway is enriched for transcripts with lengthened 3*^′^*-UTR or shortened 3*^′^*-UTR and/or intronic APA transcripts. The annotation text size for clusters is scaled by the number of nodes within each cluster. See methods for statistical parameters.

**Figure 7:**
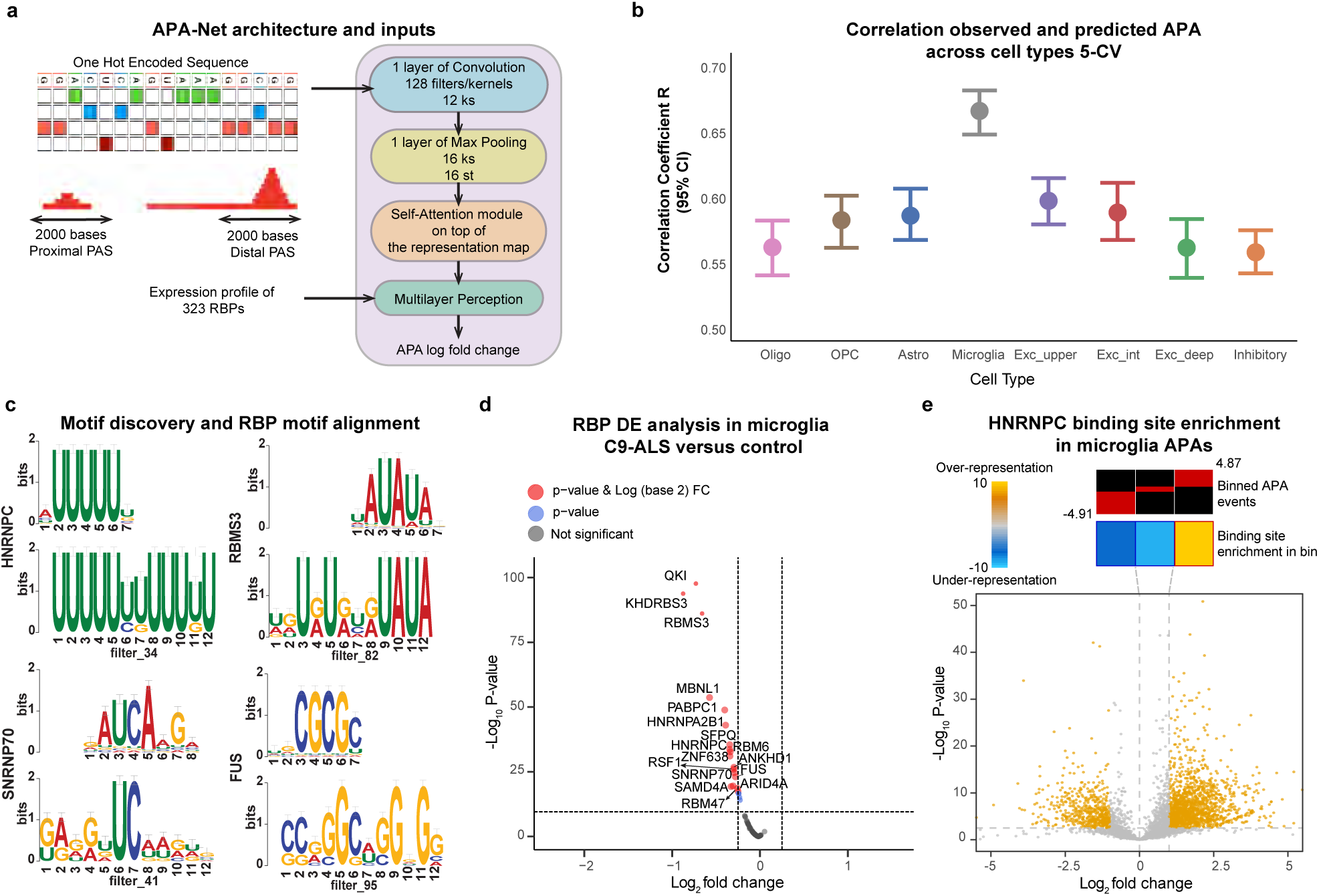
Development and analysis of a deep learning model to unravel the grammar of APA regulation in C9-ALS and sALS cases. **(a)** Schematic representation of the APA-Net architecture and inputs. The model uses two inputs: sequences surrounding the PA sites and the expression profiles of 323 RBPs. **(b)** Performance of APA-Net in predicting APA events across cell types using 5-fold cross-validation. The 95% confidence interval for correlation of observed APA LFC and predicted APA LFCs is shown. **(c)** APA-Net filters interpreted as motifs, which were subsequently aligned to an RBP database to identify corresponding RBPs. **(d)** Differential expression analysis performed on the identified RBPs. The volcano plot shows up-regulated and down-regulated RBPs in microglia from C9-ALS. **(e)** Enrichment analysis of HNRNPC binding sites for APA events in microglia. Top: APA sequences are divided into equally populated bins based on their LFC values. Red shows the proportion of APA events in each bin. Middle: Enrichment score indicating under-representation and over-representation of the binding sites. Bins with significant enrichment (hypergeometric test, corrected *P* < 0.05; red) or depletion (blue) of poly(U) motifs are denoted with a bolded border. Bottom: Volcano plot showing the distribution of changes in APA LFC in microglia from C9-ALS compared to controls. Significant observations are highlighted in orange.

We further observe lengthening and shortening events of several ALS-related genes across major cell types in C9-ALS and sALS, affecting the 3^′^-UTR and introns (Figure 6c,d, Supplementary Fig. 10a-f and Supplementary Fig. 11a-h). We see lengthening of ALS2, TARDBP, TIA1, and C9orf72 across excitatory neuron layers (Figure 6c,d, Supplementary Fig. 10c-f). Most APA events are observed in inhibitory neurons, with shortening of ANXA11 and lengthening of C9orf72 in both C9-ALS and sALS (Supplementary Fig. 10a,b). In microglia, genes such as TARDBP, MATR3, SETX, and TIA1 exhibit lengthening, while ANXA11 show 3^′^-UTR shortening (Supplementary Fig. 11g,h). These results indicate a potential role of APA in the pathogenesis of ALS by affecting ALS-related genes in a cell type-specific manner.

To assess the biological significance of APA dysregulation in C9-ALS and sALS, we analyzed pathway enrichment across neuronal and glial cell types (Figure 6e, Supplementary Fig. 12, Supplementary Fig. 13). Our analysis reveals distinct APA signatures between C9-ALS and sALS subtypes, where APA events across cell types primarily affect a greater number of pathways in sALS than in C9-ALS. Both conditions share disruptions in mRNA metabolism, splicing, and stress response pathways through 3^′^-UTR lengthening in neurons, oligodendrocytes, and microglia.

Excitatory neurons show pronounced layer-specific effects, with upper-layer excitatory neurons showing intronic APAs in glutamatergic signaling pathways (Figure 6e). Strikingly, sALS but not C9-ALS, shows a widespread switch to intronic APA usage in intermediate- and deep-layer neurons, affecting dendritic structure, synaptic organization, and calcium transport (Supplementary Fig. 12a,b). These intronic APA changes encompass known disease-associated genes such as UNC13A ^17,47,61^. In addition, sALS shows lengthening in alternative splicing pathways, while C9-ALS exhibits this pattern in deep-layer neurons.

Specific disruptions in inhibitory neurons from C9-ALS include 3^′^-UTR shortening in mitochondrial pathways, capturing changes also identified from the differential expression analysis (Supplementary Fig. 12c, Supplementary Fig. 13d). In microglia, specific changes in C9-ALS include 3^′^-UTR lengthening in myeloid differentiation and endosomal membrane response, whereas transcriptional machinery and chromatin organization are impacted in sALS (Supplementary Fig. 13d). Moreover, sALS exhibits extensive intronic APA events related to neuron projection and synapses in OPCs and astrocytes (Supplementary Fig. 13b-c), revealing pathway changes not detected by differential gene expression analysis.

Collectively, this analysis reveals distinct APA signatures between two ALS subtypes, most notably the prevalence of intronic APAs in sALS neurons and glia when compared to C9-ALS. While certain pathways are affected across multiple cell types, the cell type- and disease-specific patterns of APA regulation suggest diverse post-transcriptional mechanisms contributing to ALS pathogenesis. These APA-mediated changes often converge with pathways identified through differential gene expression analysis, highlighting the multi-modal nature of gene regulation in ALS.

### APA-Net reveals mechanistic insights into APA dysregulation in ALS

To decode the complex grammar of APA dysregulation in our ALS cohorts, we developed a deep learning model called APA-Net. To comprehensively examine APA changes, we performed differential analysis on all PA pairs of transcripts that showed significant transcript-wise APA compared to controls (Methods, Supplementary Data Table 8). We observe thousands of significant APA changes across cell types (Supplementary Fig. 14). Comparison of APA events between C9-ALS and sALS identified that 12.6-30.8% of events are shared between both disease subtypes (Supplementary Fig. 15a). These shared events demonstrate remarkably high correlation (*p* < 7.2*e* − 176) across all cell types, with robust correlation in neurons (Supplementary Fig. 15b). While most APA events are unique to either C9-ALS or sALS, suggesting distinct regulatory mechanisms in each subtype, these events maintain a positive, though weaker, correlation between the subtypes.

APA-Net is trained to predict the APA LFC (Supplementary Fig. 14) using the RNA sequences surrounding proximal and distal PA sites for each APA event, along with the RBP expression profiles per cell type (Figure 7a) as input. To assess APA-Net performance, we compared the predicted to the observed APA LFC values using Pearson correlation across 5-fold cross-validation. APA-Net achieves a strong Pearson correlation coefficient across the entire dataset (Figure 7b). Thus, APA-Net robustly learns cell type-specific APA profiles across disease subtypes, demonstrating its potential as a powerful tool for studying APA in complex diseases like ALS.

Using a convolutional neural network (CNN) architecture augmented with multi-head attention (MAT), our model is designed to identify cis-regulatory elements that influence PA site selection across different cell types. We optimized the model architecture^97,98^, including kernel size and max pooling steps, to capture relevant genomic motifs (Figure 7a). This hybrid CNN-MAT architecture enhances the model’s ability to capture both local and long-range dependencies in sequence data. This is important for accurately identifying cis-regulatory elements influencing APA, particularly those involved in binding of RBPs other than the core polyadenylation (CPA) machinery.

To identify potential cis-regulatory elements involved in APA, we used the filter weights from the CNN module of APA-Net, which represent learned sequence motifs (Supplementary Fig. 16, Methods). We scanned every sequence within the test dataset using the model’s filters to identify regional subsequences where the filters showed high activation or response. Next, we aligned RBPs from the compendium of RNA-binding motifs ^99^ with the motifs found by APA-Net (Figure 7c, Supplementary Data Table 8). We observe several ALS-related genes, such as FUS^100^, TARDBP^101^, FXR1^56^, G3BP2^102^, as well as numerous APA and AS factors such as HNRNPC, SNRNP70, SFPQ, MBNL1, and SRSF7^103–107^ among the aligned RBPs (Supplementary Data Table 8).

Differential expression analysis^108^ reveals that many of these RBPs, along with others, are significantly dysregulated across major cell types in both C9-ALS and sALS compared to controls (Supplementary Fig. 17, Supplementary Fig. 18). HNRNPC shows significant down-regulation in C9-ALS microglia (Figure 7d), where APA-Net has its strongest predictive performance, though it also remains significantly predictive across other cell types. To investigate the functional implications of this finding, we examined HNRNPC binding patterns using in-house eCLIP-seq data generated from HEK293T cells (Methods). HNRNPC is known to bind regions surrounding proximal PA sites, and its knockdown can promote distal PA site usage ^109–111^. Based on its down-regulation in C9-ALS microglia (Figure 7d), we hypothesized that HNRNPC binding sites would be enriched in proximal sites of the APA events showing lengthening. We analyzed whether HNRNPC binding sites are enriched in microglia APA events. We observe a significant enrichment of binding sites for the lengthened APA events (p-value < 1.0*e* − 04) and a lower enrichment for the shortened APA events (Figure 7e) which supports a regulatory role of HNRNPC in APA in ALS.

Overall, our findings highlight the value of APA-Net’s interpretability, enabling us to generate mechanistic hypotheses about APA dysregulation in ALS.

### RBP interactions reveal cell type-specific mechanisms and dysregulation of APA in ALS

Leveraging APA-Net’s interpretability, we examined RBPs involved in APA dysregulation in ALS. Analysis of the model’s convolutional filter activation patterns revealed co-occurring sequence motifs, which we mapped to specific RBPs. This generated RBP co-occurrence profiles for microglia, neurons, and other glial cells, which we clustered into regulatory modules based on interaction patterns (Figure 8 and Supplementary Fig. 19-Supplementary Fig. 25). These profiles suggest how combinations of RBPs may cooperatively regulate APA in different cell types. Furthermore, these profiles reveal cell type-specific patterns of RBP interactions (Figure 9a,b), with the dissimilarity heatmap (Figure 9a) demonstrating distinct regulatory networks across cell types that are not apparent from RBP expression patterns alone. While many identified modules contain RBPs from the same family or share binding preferences, a significant proportion of co-occurring RBP pairs show zero to negative motif sequence similarity (Fisher’s exact test, *p* < 2.4*e* − 46, Supplementary Fig. 26), suggesting functionally related RBPs may regulate APA through distinct binding motifs.

**Figure 8:**
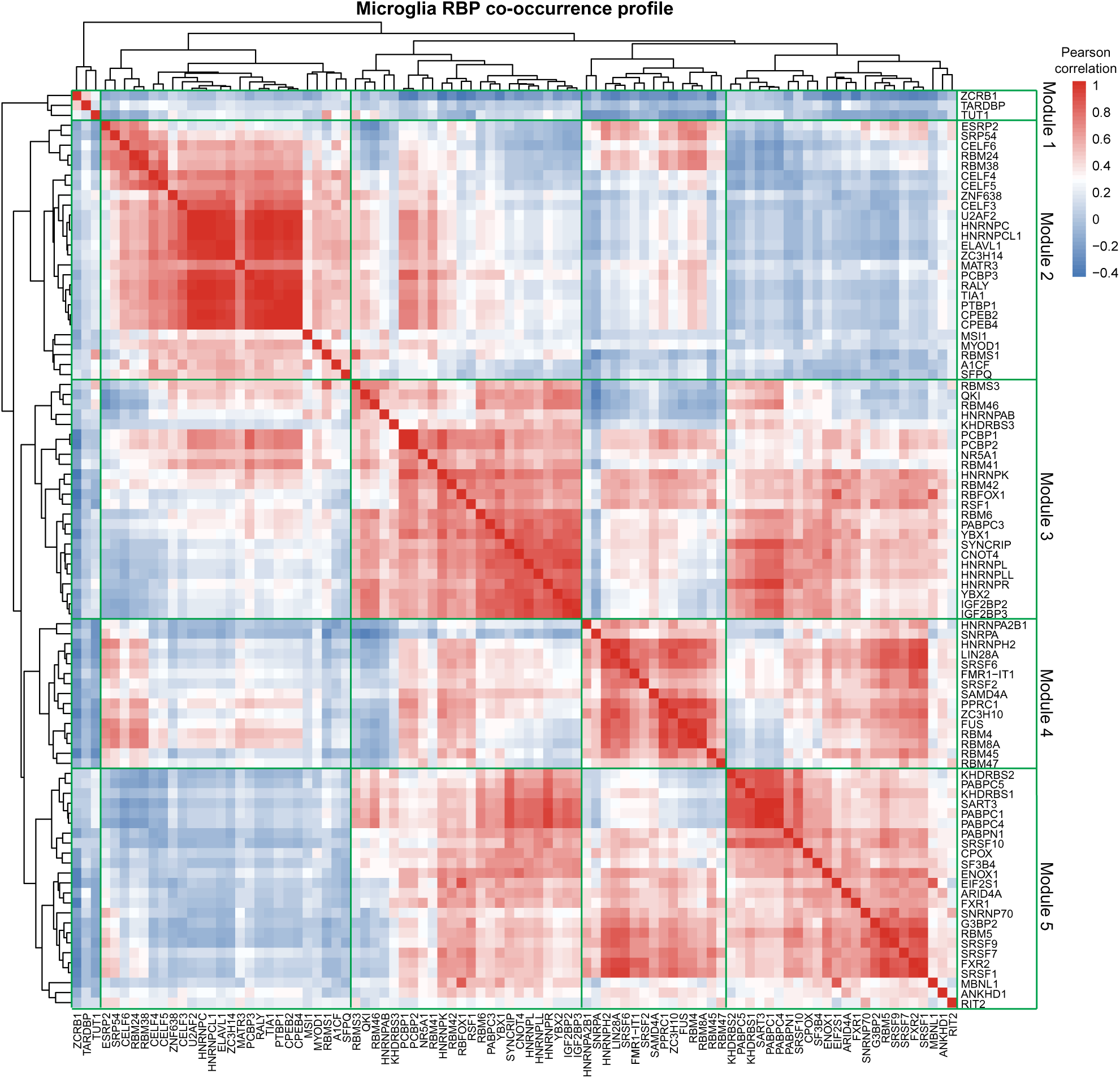
RNA-binding protein interactions in microglia reveal cell type-specific mechanisms and dysregulation of APA in C9-ALS and sALS. Clustered RBP motif co-occurrence profile for microglia. The heatmap is computed using Pearson correlation of co-occurrence of RBP-aligned filters in microglia APA events. Five RBP modules are defined based on this clustering (labeled on y- and x-axes).

**Figure 9:**
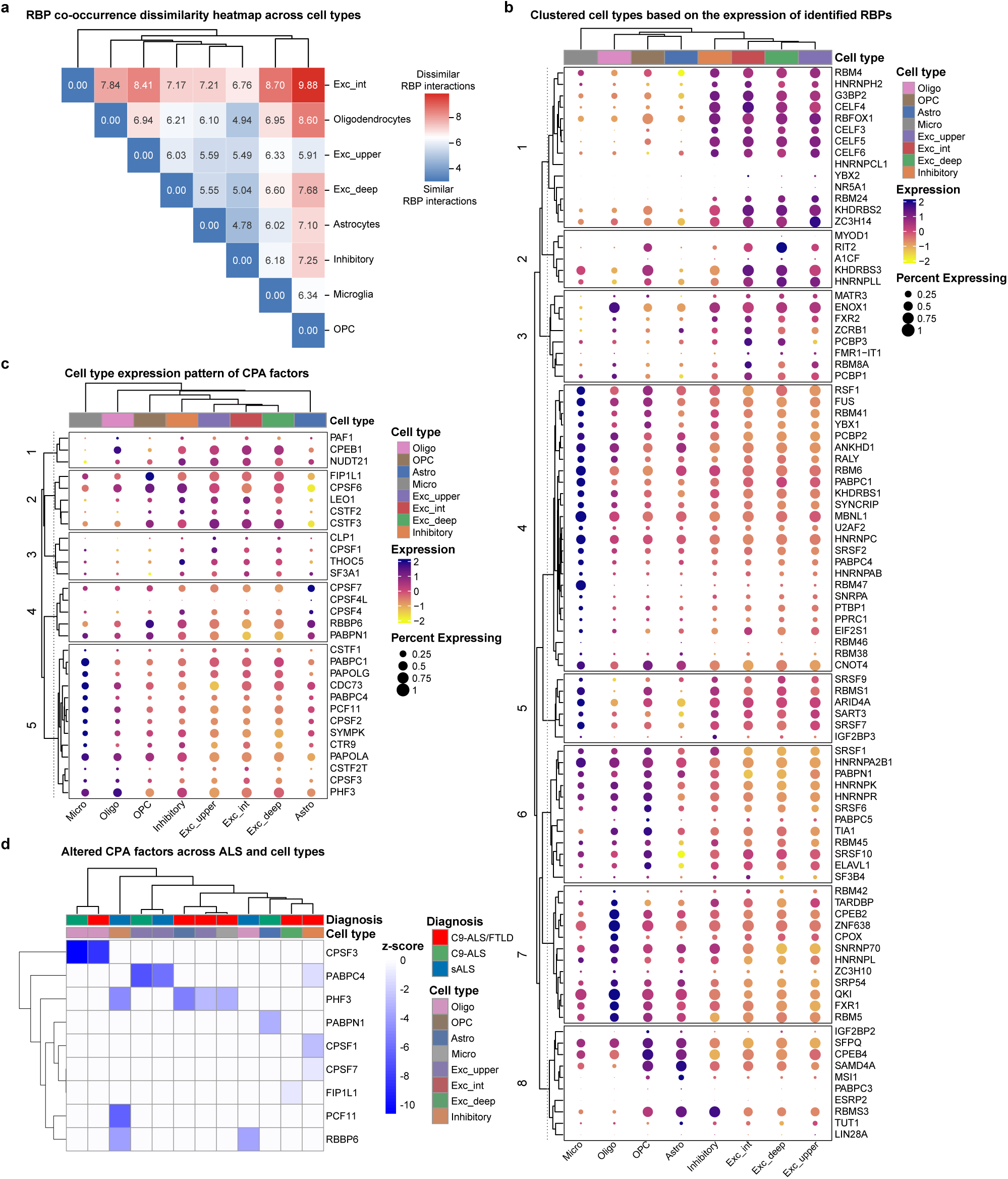
RBP co-occurrences are dissimilar across cell types and show cell type-specific expression profiles in C9-ALS and sALS. **(a)** RBP interaction dissimilarity across cell types in C9-ALS and sALS. Clustered heatmap showing the dissimilarity of RBP correlation profile interactions measured by pairwise Frobenius norm of correlation profile matrices, as depicted in Figure 8 for C9-ALS and sALS (see Methods). **(b)** Clustered heatmap dot plot analysis comparing average RBP gene expression levels (y-axis) in cell types. Size of dot corresponds to percent of cells of a given cell type expressing the corresponding gene of interest. Cell type identity indicated in the upper x-axis by colour coding and lower x-axis by label. Columns are clustered hierarchically, whereas rows are clustered and partitioned with k-means clustering (8 clusters). **(c)** Clustered heatmap dot plot analysis comparing average CPA gene expression levels. k-means clustering uncovered 5 row clusters. **(d)** Clustered heatmap of CPA genes intersecting with differentially expressed genes across cell types and ALS subtypes. Scale is in z-scores, where red scale indicates up-regulation, blue is down-regulation, white shows no significance. Significance for CPA genes is FDR < 0.01 and |LFC| < 0.5.

In microglia, Module 2 (Figure 8) highlights a diverse array of RBPs involved in various aspects of RNA processing and regulation. This module includes RBPs that regulate AS, such as ESRP2^112^, U2AF, and several members of the CUGBP Elav-like family (CELF3, CELF4, CELF5, CELF6)^113^. Additionally, Module 2 features members of the CPEB family, specifically CPEB2 and CPEB4, which bind to specific sites within the 3^′^-UTR region and regulate mRNA transport and metabolism in the cytoplasm ^114,115^. Dysregulation of CPEBs has been documented in various neuronal pathologies^115^ and is correlated with APA changes in Huntington’s disease ^116^. This module also encompasses single-stranded RBPs, including A1CF, RBMS1, and MSI1, as well as HNRNPC, which is involved in maintaining short 3^′^-UTRs, and SFPQ, which plays a role in transcript shortening by activating cryptic last exons in ALS ^117^. This clustering of diverse RBPs with roles in splicing, PA, and 3^′^-UTR processing suggests a coordinated regulation of post-transcriptional events in microglia which are relevant to ALS. Module 3 includes RBPs involved in RNA processing and stress response, such as QKI ^118^, RBOFX1^119^, and YBX1/YBX2^120^, which are important in stress granule formation. It also features RBPs with K-homologous (KH) domains, including PCBP1, PCBP2, KHDRBS3, and HNRNPK, which bind to poly(C) regions and regulate AS, stability, and translation of transcripts^121,122^.

Module 4 is characterized by RBPs involved in mRNA processing and export, such as RBM8A and RBM4^123,124^, which are also involved in AS and stress granule localization. This module includes hnRNPs such as HNRNPA2B1, HNRNPH2, and FUS, which play roles in pre-mRNA splicing and mRNA export ^12,17,125,126^. SAMD4A, which is associated with the coordinated regulation of transcription, AS, and APA ^127^, is also part of this module. The composition of Module 4 suggests an interplay between splicing regulation, mRNA export, and post-transcriptional processing, which may contribute to microglial alterations in ALS.

Module 5 includes a diverse array of RBPs involved in RNA processing, splicing, and stress response. This module features KHDRBS1 and KHDRBS2, which are involved in AS and 3^′^-UTR formation, and members of the PABC family (PABPC1, PABPC4, PABPC5) known for their role in binding to the 3^′^ poly(A) tails of eukaryotic mRNAs. Splicing regulators SART3, RBM5, and SNRNP70, as well as several members of the SR-rich family (SRSF1, SRSF7, SRSF9, SRSF10) are also included, with SRSF7 implicated in transcript shortening ^106^. Module 5 includes FXR1 and FXR2, RNA-processing proteins from the FXP family that regulate stress responses and have shown potential links to ALS pathogenesis ^56^. The co-occurrence of ALS-linked RBP motifs alongside additional splicing and PA factors in this module suggests a complex interplay between these processes in ALS.

We next investigated the expression of RBPs known to be the CPA factors regulating APA ^128^ alongside the RBPs identified by APA-Net. Across cell types, we observe eight distinct expression patterns for RBPs identified by APA-Net (Figure 9b) and five expression patterns for CPA factors (Figure 9c). Additionally, we detect cell type-specific down-regulation of several CPA factors in ALS subtypes compared to control (Figure 9d). This cell type-specificity in RBP interactions and expression profiles hints at the presence of distinct cis- and trans-regulatory elements that may influence APA in ALS. Collectively, our findings begin to map cell type-specific APA regulation in ALS and elucidate the complex molecular landscape underlying disease.

## Discussion

In this study, we present the first snRNA-seq atlas of the orbitofrontal cortex in ALS, a region associated with behavioral impairments along the ALS-FTLD spectrum^129–135^. We compared the orbitofrontal cortex with published dorsolateral prefrontal and primary motor cortex datasets^31–34^. This analysis provides a resource of concordant cell type-specific gene expression changes across ALS subtypes and affected brain regions. We characterized cell type-specific APA dysregulation in ALS subtypes compared to neurologically healthy controls. To further understand this dysregulation, we developed APA-Net, an interpretable deep learning model that integrates transcript sequences and RBP expression profiles from snRNA-seq data to predict APA changes. Our findings underscore that C9-ALS (with and without FTLD) and sALS drive distinct cellular states, marked by unique signatures in gene expression and APA profiles.

Molecular dysregulation in the orbitofrontal cortex reveals consistent patterns across the ALS-FTLD spectrum. Protein homeostasis pathways, particularly chaperone-mediated folding, show enrichment across cell types in C9-ALS/FTLD, while chromatin remodeling pathways show widespread down-regulation across multiple cell populations, including upper-layer excitatory neurons, inhibitory neurons, oligodendrocytes, and astrocytes. Cell type-specific changes in energy metabolism emerge through up-regulation of electron transport chain components in neurons, glucose metabolism in microglia, and ATP metabolism in OPCs, aligning with established patterns of mitochondrial dysfunction in ALS ^136–139^. Our analysis identified widespread dysregulation of ribosomal complexes across upper-layer, intermediate-layer, and inhibitory neuronal populations, most prominently in C9-ALS/FTLD with a subset of these changes also present in sALS.

Independent datasets strongly corroborate our molecular findings, particularly in C9-ALS/FTLD upper-layer excitatory neurons, where 115/138 genes maintain their expression patterns, highlighting disruptions in protein homeostasis, mitochondrial function, and ribosomal processes^31,32,34^. Both excitatory and inhibitory neurons demonstrate consistent up-regulation of ribosomal subunits, VCP, and SQSTM1 across cortical regions, with inhibitory neurons specifically showing increased expression of RGS4, MDH1, SLC38A2, along with similarly corroborated changes in HSP90AA1, STMN2, and NEFL ^32^. Our analysis reveals increased transcription of both STMN2 and NEFL across neuronal populations. The elevated STMN2 expression, in conjunction with its premature polyadenylation, and cryptic splicing, and loss of function with TDP-43 deficiency^25,47–49^, potentially reflects cellular compensation for compromised post-transcriptional processing. Similarly, NEFL upregulation occurs while its protein product, NfL, serves as a validated marker of axonal degeneration in patient biofluids ^140^, suggesting a coordinated neuronal response to maintain axonal integrity. Further corroborating changes include C9-ALS neurons exhibiting changes in actin dynamics (CFL1, RAC1, YWHAH) ^9^ and autophagy (HSP90AA1, HSP90AB1, DNAJB6)^32^, while sALS displays hallmark potassium channel dysregulation ^44,45,141^. The consistency of these alterations across brain regions indicates they represent fundamental disease processes rather than region-specific responses.

In glial populations, astrocytes demonstrate pronounced changes in C9-ALS/FTLD, including up-regulation of cholesterol and miRNA metabolism pathways alongside reactive markers (GFAP, CHI3L1) ^31,32^, while sALS astrocytes show TDP-43 down-regulation ^75^. Common to astrocytes across ALS subtypes is the dysregulation of protein homeostasis and actin filament organization^142^. In oligodendrocytes and OPCs, we observed consistent alterations across ALS subtypes. In sALS across independent datasets ^32^, oligodendrocytes show down-regulation of PLLP, while OPCs specifically demonstrate reduced APOD expression. Given the role of APOD in remyelination^69^, its reduction in OPCs may significantly impair the regenerative capacity of myelin in sALS. This finding aligns with broader evidence of compromised myelin maintenance across ALS subtypes, as demonstrated by the down-regulation of the myelination scaffold protein SEPTIN4 across sALS and C9-ALS/FTLD datasets. In C9-ALS oligodendrocytes specifically, we found concordant up-regulation of autophagy-related genes (ATG4B, HSP90AB1), suggesting that protein homeostasis disruption could exacerbate myelin maintenance deficits in this C9 cases.

The analysis of microglia reveals shared and unique cell type-specific changes across ALS subtypes. These include increased cytokine signaling pathways and elevated glucose metabolism, suggesting a microglial state with heightened energy demands. We find consistent patterns of depleted homeostatic markers and increased DAM stage markers (APOE, TYROBP, and SPP1) across both C9-ALS and sALS^31–34^. C9-ALS microglia specifically show impaired neuronal surveillance, marked by loss of CX3CR1 and P2RY12^81,143^, along with increased inflammatory and stress responses. Both sALS and C9-ALS exhibit altered JAK/STAT signaling, with depleted IL6R, JAK1, and JAK2 but elevated JAK3 and STAT3, supporting the therapeutic potential of JAK/STAT targeting ^85,144,145^. These microglial signatures reveal both unique and shared features with other neurodegenerative conditions, while maintaining strong concordance across existing ALS datasets.

Our APA analysis revealed both shared and subtype-specific patterns across cell types. Differential gene expression and APA changes were largely uncorrelated across all cell types and ALS subtypes. In microglia, C9-ALS showed lengthening of transcripts involved in myeloid differentiation, mitotic, and focal adhesion pathways^81,146^, whereas sALS displayed similar changes in pathways related to chromatin organization and DNA-templated transcription. Both subtypes exhibited lengthening in transcripts associated with nucleocytoplasmic transport ^6^, transcription factor binding, and stress response pathways.

Neurons display subtype-specific dysregulation of both gene expression and APA in key pathways, including mitochondrial function, stress response, synaptic signaling, and chromatin remodeling. sALS excitatory neurons show distinctive APA dysregulation in postsynaptic glutamatergic neurotransmission and calcium signaling. All neuronal subtypes exhibit down-regulation of ALS-relevant RBPs (TARDBP, FUS, ATXN2, SETX, ANXA11) and lengthening of alternative splicing transcripts, highlighting mRNA processing as a central mechanism in neuronal dysfunction ^5,144^.

Microglia highlight the distinct yet intersecting roles of APA and gene expression in regulatory mechanisms. In C9-ALS, both processes converge on pathways such as GTPase signaling and mRNA processing, but through distinct sets of genes. Conversely, sALS microglia exhibit minimal gene expression changes beyond RBP dysregulation, while showing substantial APA alterations, uncovering an unrecognized layer of transcriptional complexity in sporadic disease^94^. This integrated analysis of gene expression and APA reveals regulatory intricacies in ALS that traditional approaches would overlook.

To further investigate cell type-specific APA, we developed APA-Net. While significant progress has been made in deep learning methods for APA predictions ^147–151^, APA-Net is uniquely trained on cell type-specific APA profiles from snRNA-seq data. Using APA-Net, we identified cis-regulatory elements and cell type-specific RBP interactions directly from the model predictions. This analysis revealed RBPs and their potential interactions regulating APA across ALS subtypes. This approach not only enhances our understanding of APA regulation but also reveals the complex interplay of RBPs in disease contexts, providing a valuable tool for exploring molecular mechanisms of APA in various diseases.

Our single-nucleus transcriptomic profiling of the orbitofrontal cortex provides a granular understanding of cellular heterogeneity in ALS and FTLD, complementing existing single-cell ALS mapping efforts ^31–34^. Through evaluation of the orbitofrontal cortex, we reveal concordant and divergent ALS-associated changes across cell types, brain regions, and disease subtypes. Moreover, APA-Net’s ability to predict cell type-specific APA events can be applied to other single-cell transcriptomic atlases. Overall, our findings illuminate the complex, cell type-specific pathomechanisms of ALS, offering a valuable resource for advancing therapeutic research in ALS and other neurodegenerative diseases.

## Limitations of this study

Building upon our findings, several promising avenues for future research emerge. Expanding our cohorts from the orbitofrontal cortex would enable deeper exploration of disease heterogeneity across ALS and FTLD subtypes and better account for age-related variability^152^. Integration of multiple brain and spinal cord regions would further strengthen our findings and enable more comprehensive analyses. For instance, including a sALS cohort with FTLD would illuminate changes specific to C9-ALS compared with sALS, with and without FTLD. Our findings provide a foundation for validation through complementary approaches. Spatial transcriptomics could extend these results by preserving crucial spatial information, while protein-level studies could confirm key molecular changes. For APA analysis, our robust findings from pseudo-bulk snRNA-seq data could be enhanced through long-read sequencing and cell type-specific sorting of cerebral cortex nuclei^153^. These approaches, particularly QuantSeq REV on sorted nuclei, would provide additional validation of APA events. These future directions will build upon our current results, further advancing our understanding of ALS at the molecular level and potentially revealing new therapeutic targets.

## Methods

### Human brain samples

Informed consent was obtained from all participants in accordance with the Ethics Review Boards at Sunnybrook Health Sciences Centre and University of Toronto. ALS clinical diagnosis was determined based on the El Escorial revisited clinical criteria ^154^. Fresh frozen orbitofrontal cortex tissues were collected from ALS cases with pathologically confirmed FTLD or no FTLD, and six non-neurological disease control cases (detailed information in Supplementary Data Table 1). C9-genotypes were determined as described previously^155,156^. Postmortem patient samples comprised C9-ALS/FTLD (n=6), C9-ALS with no FTLD (n=3), and sALS with no FTLD (n=8). Postmortem non-neurological control samples (n=6) were obtained from the Douglas-Bell Canada Brain Bank (DBCBB, Montreal, Canada) (n=4) or University Health Network–Neurodegenerative Brain Collection (UHN–NBC, Toronto, Canada) (n=2). Expert neuroanatomists ensured that each orbitofrontal cortex tissue section contained the entire laminar structure of the cortex (layers I-VI and white matter). A subset of samples were processed using the 10X v2 chemistry for C9-ALS/FTLD (n=3), C9-ALS no FTLD (n=1), and sALS no FTLD (n=4). These v2 samples were used solely for an in-depth cell type annotation process across disease subtypes. For all downstream analyses on snRNA-seq, exclusively v3 chemistry samples were used. Here, the C9-ALS/FTLD (n=4) and C9-ALS without FTLD (n=2) were combined to create the C9-ALS cohort (n=6). The sALS without FTLD (n=4) samples constituted the sALS cohort. These disease subtypes (C9-ALS and sALS) were then compared to the non-neurological disease control group (n=6).

### Single nucleus RNA-seq by FACS

Frozen orbitofrontal cortex (50mg per sample) was dounce homogenized on ice in lysis buffer (0.32mM sucrose, 5mM CaCl2, 3mM Mg(Ac)2, 20mM Tris-HCl [pH 7.5], 0.1% Triton X-100, 0.5M EDTA [pH 8.0], 40U/mL RNase inhibitor in H2O), centrifuged at 800×g for 10 min at 4°C. The supernatant was removed and the pellet was washed twice and resuspended in a resuspension buffer (1x PBS, 1% BSA, 0.2U/µL RNase inhibitor). Resuspended nuclei were sorted by FACS with DAPI (Roche) labeling, removing any debris and nuclei aggregates within DAPI-positive gating, aiming to capture approximately 6000 nuclei per sample. Library preparation was performed using either the 10X Chromium Single Cell 3’ v2 or v3 platform following the manufacturer protocols. The QC of cDNA libraries was conducted on a 2100 Bioanalyzer (Agilent). The cDNA libraries were 100-bp paired end sequenced on either an Illumina NovaSeq 6000 SP XP or NovaSeq6000 S2 standard flow cell at the Princess Margaret Genomic Centre (Toronto, Ontario). Raw Illumina base call files from each sample were demultiplexed to produce FASTQ files with the cellranger mkfastq pipeline (10X Genomics). Reads were aligned to the pre-mRNA GRCh38-2020-A genome and quantified using cellranger count command on Cell Ranger v5.0.0 (10X Genomics).

### snRNA-seq sample processing, QC, and clustering

All processing and QC of snRNA-seq samples was performed using Seurat (v4), and custom R and Python scripts. Seurat objects were created for each sample using the filtered feature-barcode matrices obtained from Cell Ranger (v5.0.0). For each sample, nuclei containing mitochondrial reads with a threshold greater than three mean absolute deviations from the median number of mitochondrial reads with a maximum cut-off of 5% were removed. Next, nuclei with fewer than 200 and greater than 12000 detected genes were removed. Reads pertaining to the cell cycle were scored using the scran R package^157^. Potential doublets were estimated and removed using scDblFinder with default parameters ^158^. The remaining singlet transcriptomes were merged and batch corrected with Harmony ^36^ on log1p normalized counts. Dimensionality reduction was performed using principal component analysis (PCA) on 50 principal components and then visualized using uniform manifold projection and approximation (UMAP) ^159^. Clustering was performed using the Leiden algorithm ^37^ at a resolution of 0.6. Clusters with fewer than 200 cells were filtered out as background ^160^ based on poor representation across samples and disease subtypes. Visualization was performed using the scCustomize ^161^ or dittoSeq^162^ R packages.

### Cluster annotation by machine learning and reference atlas

To uncover orbitofrontal cell types from the identified clusters, we employed marker discovery by machine learning (NSForest v3.9.1) ^38^ and reference-based annotation^39^ using the Allen Brain Atlas http://portal.brain-map.org/atlases-and-data/rnaseq^163^. Default parameters in NSForest were applied to identify binary markers for each cluster. The machine learning-based marker classification approach identified binary markers for each cluster using random forest feature selection and expression scoring. These binary markers for clusters were then used to confirm cortical cell type identity using reference-based annotation. Markers from each dataset were then compared with reference 10X Genomics experiments from two cortical regions in humans, including the primary motor cortex, dorsolateral prefrontal cortex, and medial temporal gyrus from the Seattle Alzheimer Disease Cell Atlas https://portal.brain-map.org/atlases-and-data/rnaseq^163,164^. Final confirmation of cell type identity was confirmed for each cell subtype by manual annotation^39^ using canonical cortical cell markers.

### Differential gene expression analysis

Differentially expressed genes across ALS subtypes relative to control in each cell types was uncovered using the model-based analysis of single-cell transcriptomes (MAST) approach ^40^. As described above, the 10X v3 samples were included in the differential expression analysis to account for differences in the number of detected genes and the unique molecular index (UMI) distribution. A mixed-effect hurdle model was employed with the counts data, analyzing log2-normalized counts as described ^40^. The model was designed to include both fixed effects and the individual sample as a random effect as follows:

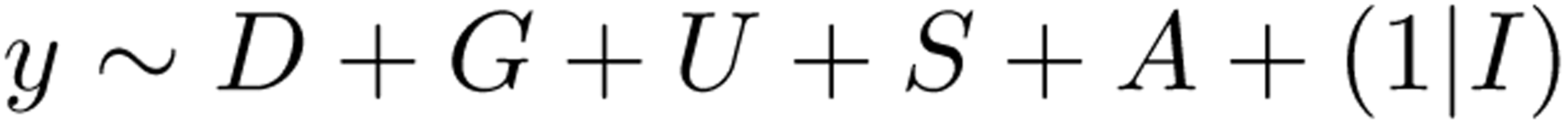

where fixed effects are denoted as *y* for the log2-normalized count of the gene, *D* for the diagnosis (C9-ALS/FTLD vs control, C9-ALS vs control, and sALS vs control), *G* is the number of genes detected, *U* is the UMI distribution, *S* is the sex of the donor, *A* is the age of the donor, and 1|*I* is the random effect for donor ID, accounting for donor-specific variability.

The hurdle component modeled the probability of zero versus non-zero counts using logistic regression, while the count component modeled non-zero counts using a Poisson distribution. This approach enabled us to address the excess zeros in our data and assess the influence of each predictor on the gene expression counts

Next, we performed a likelihood ratio test to identify differentially expressed genes by comparing models with and without the diagnosis term, as described^31^. Briefly, hurdle p-values were reported using the MAST method, and p-values were adjusted for multiple comparisons using the Benjamini and Hochberg FDR method. The MAST model provided LFC due to the disease effect, derived from both the continuous component (nonzero expression) and the discrete component (expressed or not) of the hurdle model. Additionally, we computed average LFC by subtracting the mean log2 counts per million of control nuclei from that of disease nuclei, ensuring consistency between model LFC and average fold change (FC), and excluding genes with significant discrepancies. We considered significance set at FDR < 0.01 and model |LFC| < 0.5. Further analysis of transcriptional signatures within cell types and across disease subtypes was performed using the R package UCell version 2.4^165^. UCell uses the Mann-Whitney U statistic to assign and rank signature scores from a set of pre-determined genes based on their relative expression in cells.

### Comparison with existing ALS datasets

We retrieved pre-processed snRNA-seq data and relevant metadata from three independent ALS/FTLD studies, including Pineda et al. ^32^ from https://www.synapse.org/Synapse:syn51105515/files/ and https://www.ncbi.nlm.nih.gov/bioproject/PRJNA1073234/; Li et al. ^31^ from https://www.ncbi.nlm.nih.gov/geo/query/acc.cgi?acc=GSE219281, and Gittings et al. ^33^ from https://www. synapse.org/Synapse:syn45351388/files/. To allow for the comparative analysis and visualization of gene expression changes, we created Seurat objects for each dataset as described above using the meta-data provided by each study. For differentially expressed genes, we retrieved differential expression data from the published supplementary tables where available ^31,32,34^. For each cell type, we considered significance at either an adjusted p-value or FDR < 0.01 and |LFC| < 0.5. Gene intersections were visualized using Venn diagrams.

### APA quantification and profiling

Pseudo-bulk alignment files were generated for each cell type using the barcodes. We employed the MAAPER software ^92^ to assign sequencing reads to known PA, as defined in the PolyA DB v3 database ^93^. PA sites were considered only if there were at least 25 reads aligning to the sites. To identify genes with significant changes in the length of their 3^′^-most exon, we used the REDu metric provided by MAAPER. REDu measures the relative expression levels between the two most differentially expressed isoforms in the 3^′^-most exon. A positive REDu value indicates transcript lengthening events, while a negative value points to shortening events. For pinpointing genes exhibiting intronic APA usage, we used the REDi metric. REDi compares the relative expression levels of the top differentially expressed isoform in the 3^′^-most exon and the top differentially expressed isoform in an intron or internal exon. The RED score, comparing conditions 1 and 2, is computed using the formula:

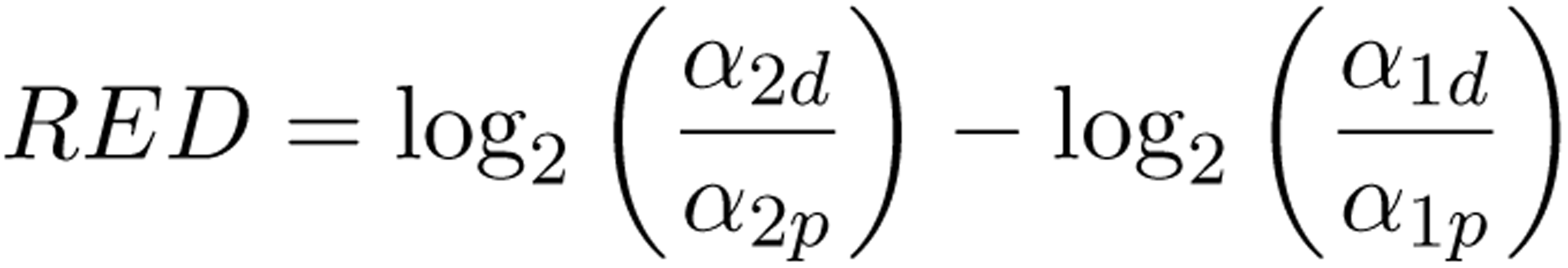

where the proportions of the distal and proximal PAs in conditions 1 and 2 are denoted as *α*_2d_, *α*_2p_, *α*_1d_, and *α*_1p_ respectively ^92,93^.

More than 80% of the genes in PolyA DB v3 database ^93^ have 3 or more annotated PAs. Hence, to investigate differential poly(A) site usage patterns in more detail, we used the APAlog package. APAlog operates on the normalized counts of reads mapped to each PA site to assess the extent and nature of differential usage. For a comprehensive comparison, APAlog was run in Pairwise Test mode, which enables the comparison of all possible pairs of PA sites per transcript^111^. Similarly, the APA LFC values were calculated to identify distal and proximal PA site usage:

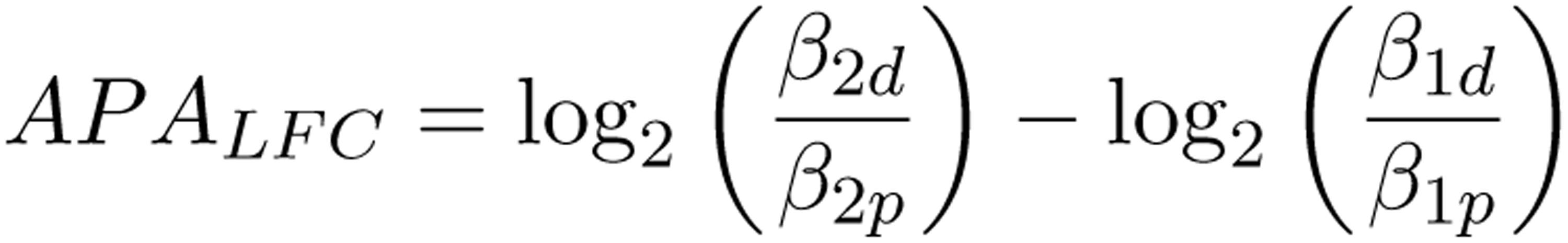

where the proportions of the distal and proximal PAs in conditions 1 and 2 are denoted as *β*_2d_, *β*_2p_, *β*_1d_, and *β*_1p_ respectively.

### Empirical FDR analysis for APA

We conducted an empirical FDR analysis using the controls from individuals without neurological diseases to mitigate false positives. This enabled us to establish cell type-specific FDR thresholds where only 10% of the p-values fell below each threshold (Supplementary Data Table 6). For downstream analysis, we included only transcripts that passed this FDR analysis, ensuring a reliable FDR of 10% or less in our results.

APA analysis was performed on all possible pairs of control vs. control samples, and transcript-level APA p-values were examined. The threshold was selected based on the 10% quantile. For further analysis, we retained only significant APA events after applying the Benjamini-Hochberg FDR correction to the RED values from MAAPER. Similarly, APAlog was used to identify transcripts with significant APA, and the Benjamini-Yekutieli method was applied to correct the p-values of APA LFC values due to the mutual non-independence of p-values from testing pairs of PA sites in transcripts with more than two sites.

### Pathway Enrichment Analysis

Pathway enrichment analyses for snRNA-seq data, including from differentially expressed genes and APA, were performed using gProfiler2^41^ with curated Gene ontology (GO) biological process and cellular compartment gene sets with no inferred electronic annotation were downloaded from http://baderlab.org/GeneSets/ (April 2024 release). We used a minimum gene set size of 15 and the maximum gene set size of 200. Visualization of GO results focused on minimizing GO term redundancy using either individual enrichment or bar plots of select representative findings, or by summarizing results by plotting enrichment maps created with the Enrichment Map plugin for Cytoscape (v3.9.1) in Linux or Windows ^166^. A q-value cutoff < 0.1 and edge cutoff (similarity) of 0.375 was used for plotting either the differentially expressed genes or APA enrichment map results.

### Deep learning model architecture

APA-Net architecture is designed to map the complex regulatory mechanisms underlying APA from 3^′^ single-cell transcriptomics data. Hybrid models that combine convolutional neural networks (CNNs) with architectures traditionally developed for natural language processing have emerged as powerful tools in genomic sequence analysis ^167,168^. APA-Net employs a CNN architecture supplemented with a multi-head attention module, specifically optimized to identify cis-regulatory elements impacting PA site selection across varied cell types. The input region for the CNN and MAT modules encompasses 2 kb surrounding both the proximal and distal PAS. The CNN module contains a single convolutional layer, comprising 128 kernels, each with a size of 12 and a stride of 1. This is followed by a max pooling layer with a kernel size of 16 and stride of 16. The output from this stage feeds into a multi-head attention module, in which each position in the representation map functions as a distinct token. In this context, a ’token’ refers to a discrete unit of information, which is essential for the attention mechanism to effectively process and interpret the complex patterns within the data. A residual connection links the CNN and multi-head attention modules. This residual connection, a key component in deep learning architectures, helps in mitigating the vanishing gradient problem by allowing the flow of information and gradients directly across layers. The attention module’s output, along with the RBP expression profile specific to the cell type, is forwarded to a multi-layer perceptron for final APA effect prediction through a regression task.

The objective of APA-Net therefore is to predict the APA LFC values and the loss for regression can be represented as:

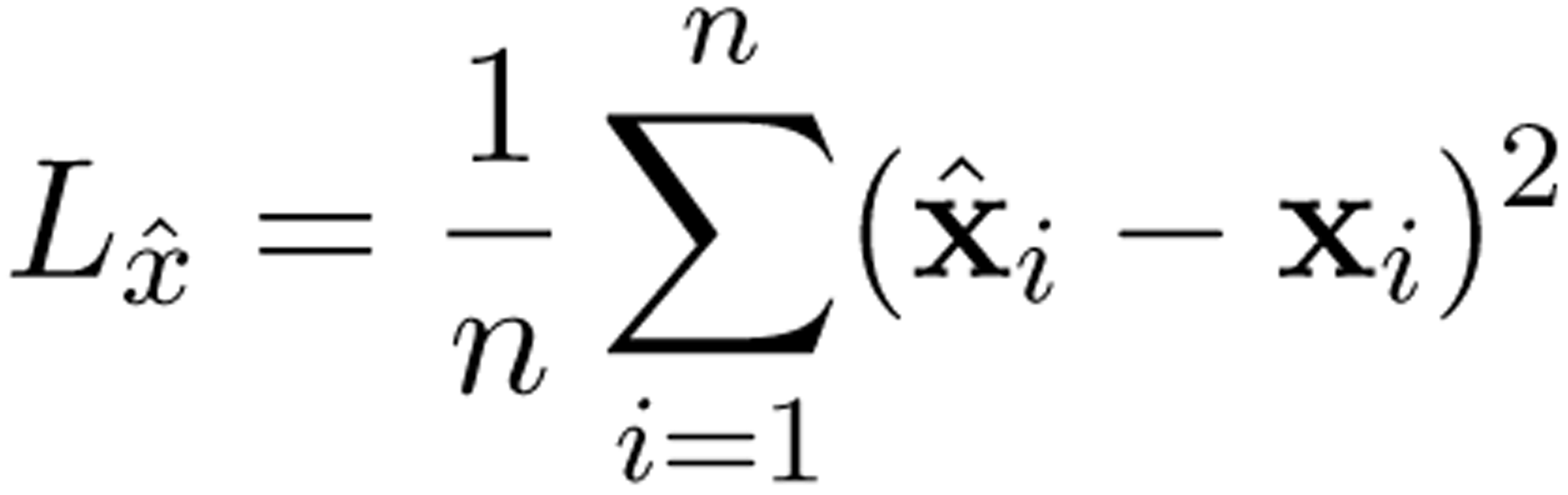

To train APA-Net, we utilized the pairwise APA events identified by APAlog across different cell types. For each APA event, we extracted genomic sequences from the hg38 reference genome surrounding both the proximal and distal PAS, transcribed them to RNA sequences, and oriented them in the sense direction to preserve the proximal/distal usage information. These sequences served as input features, along with the cell-type specific RBP expression profiles. The corresponding APA log fold change (LFC) values were used as regression labels. We employed 5-fold cross-validation, where for each fold, the data was split into 80% training and 20% testing sets while maintaining the representation of APA events across different cell types. This cross-validation approach ensured robust evaluation of the model’s performance across different subsets of the data.

### Deep learning model interpretation

To identify potential cis-regulatory elements, we used the filter weights from the CNN module of APA-Net, which represent learned sequence motifs. Each sequence within the test dataset was scanned using the model’s filters to identify regions with maximal filter activation, hypothesized to correspond to biologically relevant RNA sequence motifs. These motifs were modeled as Position-Weight Matrices (PWMs) to capture the patterns each filter had learned for APA sequences in ALS.

Next, we aligned RBPs from the compendium of RNA-binding motifs ^99^ with the motifs identified by APA-Net. This alignment enabled us to associate the learned motifs with known RBPs. To identify differentially expressed RBPs, we employed DESeq2^108^. This pseudobulk approach enabled the preservation of biological relevance and expected effect sizes of altered RBPs while mitigating cell type variability^169–171^. We used the FindMarkers function from Seurat with the list of identified RBPs to perform the analysis. We then applied stringent statistical cutoffs of adjusted p-value < 0.01 and used a |LFC| < 0.25 to capture a large net of dysregulated RBPs.

### HNRNPC eCLIP-seq processing and enrichment analysis

For eCLIP-seq data processing, reads were initially processed using UMI-tools to extract unique molecular identifiers, followed by quality filtering (Q > 15) and adapter removal using cutadapt. The processed reads were aligned to the human genome (hg38) using BWA (v0.7.17). PCR duplicates were removed and peaks were called using the CLIP Tool Kit (CTK v1.1.13) implementing a valley-seeking algorithm with multiple testing correction. Peak boundaries were determined by combining replicates. We used FIRE ^172^ for binding site enrichment analysis.

### RBP interaction dissimilarity across cell types

We used the Frobenius norm, a measure of the difference between two matrices (similar to Euclidean distance for vectors), defined as the square root of the sum of the absolute squares of their element-wise differences, to measure the dissimilarity of the RBPs interaction profiles across cell types. The Frobenius distance between two matrices *X* and *Y* is given by:

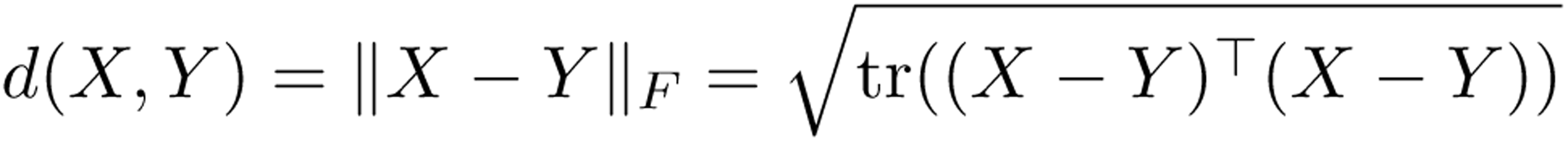

where:

• tr(*A*) denotes the trace of matrix *A*, which is the sum of its diagonal elements.
• *A*^⊤^ denotes the transpose of matrix *A*.
• ∥ · ∥_F_ denotes the Frobenius norm.

## Data and code availability

Raw FASTQ snRNA-seq files are deposited in the National Institute of Health Sequencing Read Archive at https://www.ncbi.nlm.nih.gov/sra?term=PRJNA918304. Source code used in this study is available on GitHub: https://github.com/BaderLab/APA-Net and https://github.com/AidenSb/ALS_APA_2024.

## Acknowledgments

The authors are thankful to the ALS patients and families for their tissue donations in this study as well as the Douglas-Bell Canada Brain Bank (McGill University). This research was enabled in part by access to high performance computing clusters provided through the Digital Research Alliance of Canada https://alliancecan.ca. This work was funded by the James Hunter Family Initiative in ALS Research, a Project Grant from the ALS Society of Canada, and an ERA-LEARN (E-Rare-3/Canadian Institute for Health Research) grant (063-REPETOMICS). P.M. McKeever was supported by the ALS Association Milton Safenowitz Postdoctoral Fellowship (2019-2021) and currently holds the Christopher Chiu Fellowship from ALS Double Play. J. Robertson is the James Hunter Family Chair in ALS Research. E. Rogaeva is supported in part by the Canadian Consortium on Neurodegeneration in Aging. This work was supported by NRNB (U.S. National Institutes of Health, National Center for Research Resources grant number P41 GM103504)

## Notes

### Competing Interest Statement

The authors have declared no competing interest.

### Summary of Updates

In response to the Referees' constructive criticisms, we have substantially strengthened our analysis and presentation of these findings. Specifically, we have: 1. Addressed age as a potential confounder by implementing it as a covariate in our differential expression model design, better ensuring our findings represent disease-associated changes. 2. Integrated our findings with four independent datasets from the dorsolateral prefrontal and primary motor cortex, corroborating consistent molecular signatures across ALS subtypes. 3. Improved our statistical reporting with comprehensive significance thresholds, empirical FDR analysis, and detailed methodological explanation throughout. 4. Expanded our analysis of cell type-specific expression patterns and APA alterations across major cell populations, revealing both shared disease mechanisms and subtype-specific changes. 5. Strengthened APA-Net's technical performance in predicting APA events through cross-validation (correlation coefficient 0.55-0.68 across cell types). These revisions have resulted in a more focused examination of transcriptional dysregulation in ALS that provides novel insights into cell type-specific disease mechanisms.

