## Supplementary Tables 1,6; Supplementary Figures 1-26 for "Single-nucleus transcriptome atlas of orbitofrontal cortex in amyotrophic lateral sclerosis with a deep learning-based decoding of alternative polyadenylation mechanisms"

### 1 Supplementary Data

**Supplementary Data Table 1:** Patient demographic information

| Sample | Pathological diagnosis | Sex | Age onset | Age death | Disease duration | C9 alleles | TDP-43 path.* | Tissue source | snRNA |
|---|---|---|---|---|---|---|---|---|---|
| CTRL1 | Non-neurological disease | F | NA | 50 | 0 | 2/2 | No | DBCBB | 10X 3′ v3 |
| CTRL2 | Non-neurological disease | M | NA | 59 | 0 | 2/2 | No | DBCBB | 10X 3′ v3 |
| CTRL3 | Non-neurological disease | F | NA | 72 | 0 | 2/2 | No | DBCBB | 10X 3′ v3 |
| CTRL4 | Non-neurological disease | M | NA | 72 | 0 | 2/2 | No | DBCBB | 10X 3′ v3 |
| CTRL5 | Non-neurological disease | F | NA | 53 | 0 | 2/2 | No | UHN-NBC | 10X 3′ v3 |
| CTRL6 | Non-neurological disease | F | NA | 48 | 0 | 2/2 | No | UHN-NBC | 10X 3′ v3 |
| C9ALSFTLD1 | ALS with FTLD | M | 59 | 60 | 1 | 2/exp | Yes | Tanz CRND | 10X 3′ v3 |
| C9ALSFTLD2 | ALS with FTLD | M | 57 | 60 | 3 | 2/exp | Yes | Tanz CRND | 10X 3′ v3 |
| C9ALSFTLD3 | ALS with FTLD | M | 57 | 58 | 1 | 2/exp | Yes | Tanz CRND | 10X 3′ v3 |
| C9ALSFTLD4 | ALS with FTLD | F | 45 | 47 | 2 | 2/exp | Yes | Tanz CRND | 10X 3′ v3 |
| C9ALSFTLD5 | ALS with FTLD | F | 58 | 59 | 1 | 2/exp | Yes | Tanz CRND | 10X 3′ v2 |
| C9ALSFTLD6 | ALS with FTLD | M | 70 | 72 | 2 | 2/exp | Yes | Tanz CRND | 10X 3′ v2 |
| C9ALSnoFTLD1 | ALS with no FTLD | F | 67 | 71 | 4 | 2/exp | No | Tanz CRND | 10X 3′ v3 |
| C9ALSnoFTLD2 | ALS with no FTLD | F | 65 | 66 | 1 | 2/exp | No | Tanz CRND | 10X 3′ v2 |
| C9ALSnoFTLD3 | ALS with no FTLD | F | 85 | 88 | 3 | 2/exp | No | Tanz CRND | 10X 3′ v3 |
| sALSnoFTLD1 | ALS with no FTLD | M | 51 | 59 | 8 | 2/2 | No | Tanz CRND | 10X 3′ v2 |
| sALSnoFTLD2 | ALS with no FTLD | F | 54 | 57 | 3 | 2/2 | No | Tanz CRND | 10X 3′ v2 |
| sALSnoFTLD3 | ALS with no FTLD | M | 65 | 75 | 10 | 2/2 | No | Tanz CRND | 10X 3′ v3 |
| sALSnoFTLD4 | ALS with no FTLD | F | 87 | 88 | 1 | 2/2 | No | Tanz CRND | 10X 3′ v3 |
| sALSnoFTLD5 | ALS with no FTLD | M | 39 | 43 | 4 | 2/2 | No | Tanz CRND | 10X 3′ v3 |
| sALSnoFTLD6 | ALS with no FTLD | F | 70 | 72 | 2 | 2/2 | No | Tanz CRND | 10X 3′ v2 |
| sALSnoFTLD7 | ALS with no FTLD | M | 32 | 37 | 5 | 2/2 | No | Tanz CRND | 10X 3′ v2 |
| sALSnoFTLD8 | ALS with no FTLD | M | 47 | 50 | 3 | 2/2 | No | Tanz CRND | 10X 3′ v3 |

ALS = amyotrophic lateral sclerosis; FTLD = frontotemporal lobar degeneration;
exp = hexanucleotide repeat expanded allele in C9orf72;
DBCBB = Douglas Bell Canada Brain Bank;
UHN-NBC = University Health Network – Neurodegenerative Brain Collection;
Tanz CRND = Tanz Centre for Research in Neurodegenerative Diseases;
* presence of TDP-43 pathology indicates FTLD by immunohistochemical labeling with phosphoTDP-43

**Supplementary Data Table 6:** FDR Thresholds for MAAPER and APAlog Analyses Across Cell Types

| Analysis Type | Cell Type | FDR 10% Threshold |
|---|---|---|
| MAAPER | Astrocyte | $1.800 \times 10^{-14}$ |
| | Excitatory (deep) | $4.768 \times 10^{-14}$ |
| | Excitatory (intermediate) | $6.595 \times 10^{-16}$ |
| | Excitatory (upper) | $2.819 \times 10^{-20}$ |
| | Inhibitory | $1.140 \times 10^{-21}$ |
| | Microglia | $9.021 \times 10^{-11}$ |
| | Oligodendrocyte | $1.230 \times 10^{-20}$ |
| | OPC | $2.820 \times 10^{-12}$ |
| APAlog | Astrocyte | $3.493 \times 10^{-4}$ |
| | Excitatory (deep) | $1.062 \times 10^{-6}$ |
| | Excitatory (intermediate) | $5.272 \times 10^{-7}$ |
| | Excitatory (upper) | $1.910 \times 10^{-5}$ |
| | Inhibitory | $1.446 \times 10^{-4}$ |
| | Microglia | $1.826 \times 10^{-5}$ |
| | Oligodendrocyte | $5.378 \times 10^{-3}$ |
| | OPC | $2.278 \times 10^{-4}$ |

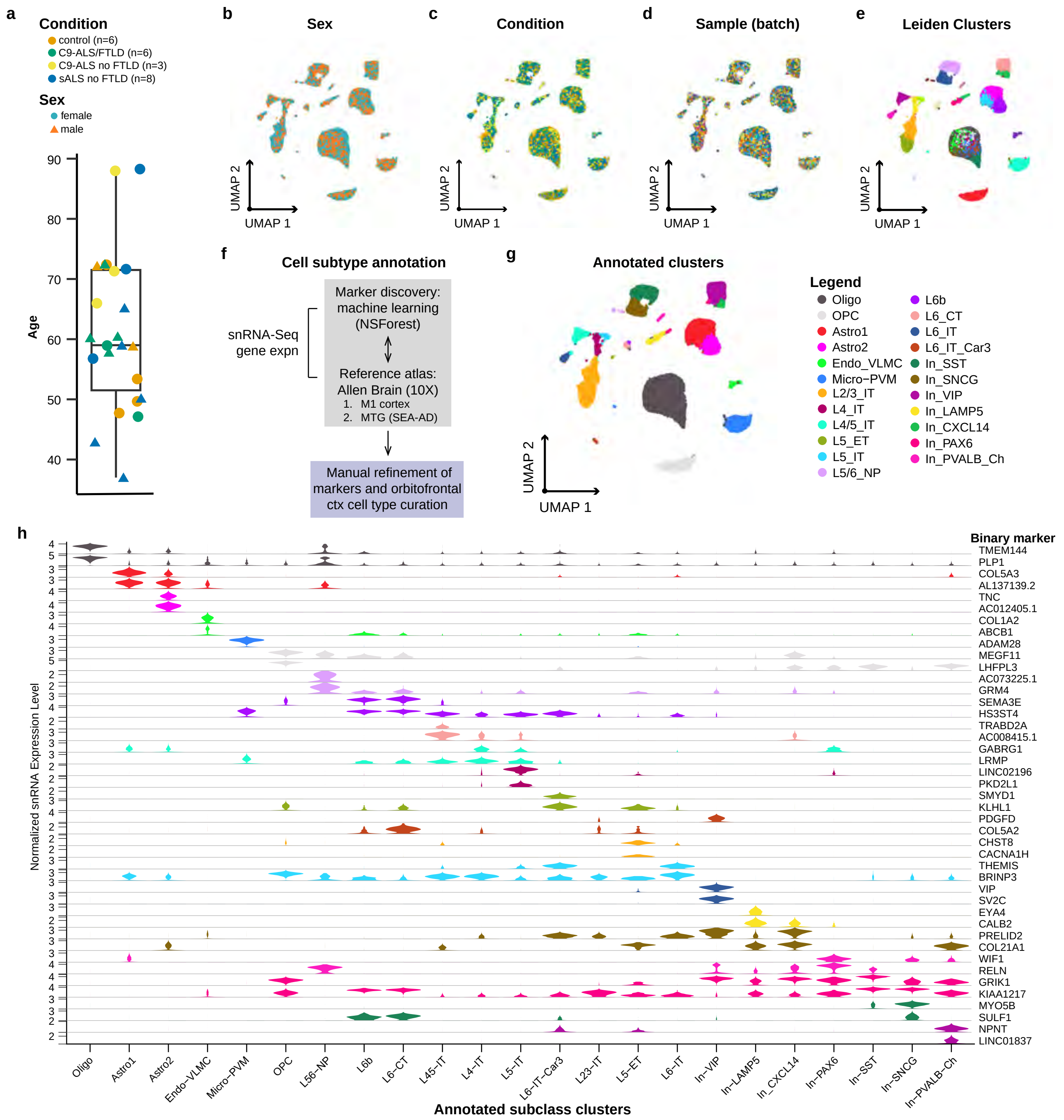


**Supplementary Fig. 1: Annotation of cell subtypes across ALS.** **(a)** Barplot of age (61.6 +/- 13.3 years) across 23 samples and annotated by condition (point colour) and sex (point shape)). **(b)** UMAP plot integrated snRNA-seq data labeled by sex (male and female). **(c)** Same as (b) for condition (control, C9-ALS/FTLD, C9-ALS no FTLD, sALS no FTLD). **(d)** Same as (b) for all 23 samples. **(e)** Same as (b) for Leiden clusters. **(f)** Cell subtype annotation strategy, consisting of a marker discovery phase, reference-based phase, and manual curation/refinement (see Methods). **(g)** Annotated clusters for all identified cell subtypes. **(e)** Binary markers (y-axis) per cell type (x-axis) in the identified using NSForest[38]. Oligo = oligodendrocytes; Astro = astrocytes; Endo-VLMC = endothelial and VLMC; Micro-PVM = microglia and PVM; excitatory neuron subtypes = Layer 2/3 intratelencephalic (L2/3 IT), L4 IT, L4/5 IT, L5 extra telencephalic-projecting (L5 ET), L5 IT, L5/6 near projecting (L5/6 NP) L6B, L6 corticothalamic-projecting (L6 CT), L6 IT, L6 IT Car3; inhibitory neuron subtypes: IN-SST (SST+), In-SNCG (SNCG+), In-VIP (VIP+), In-LAMP5 (LAMP5+), In-CXCL14 (CXCL14+), In-PAX6 (PAX6+), and In-PVALB-Ch (PVALB+ chandelier cells).

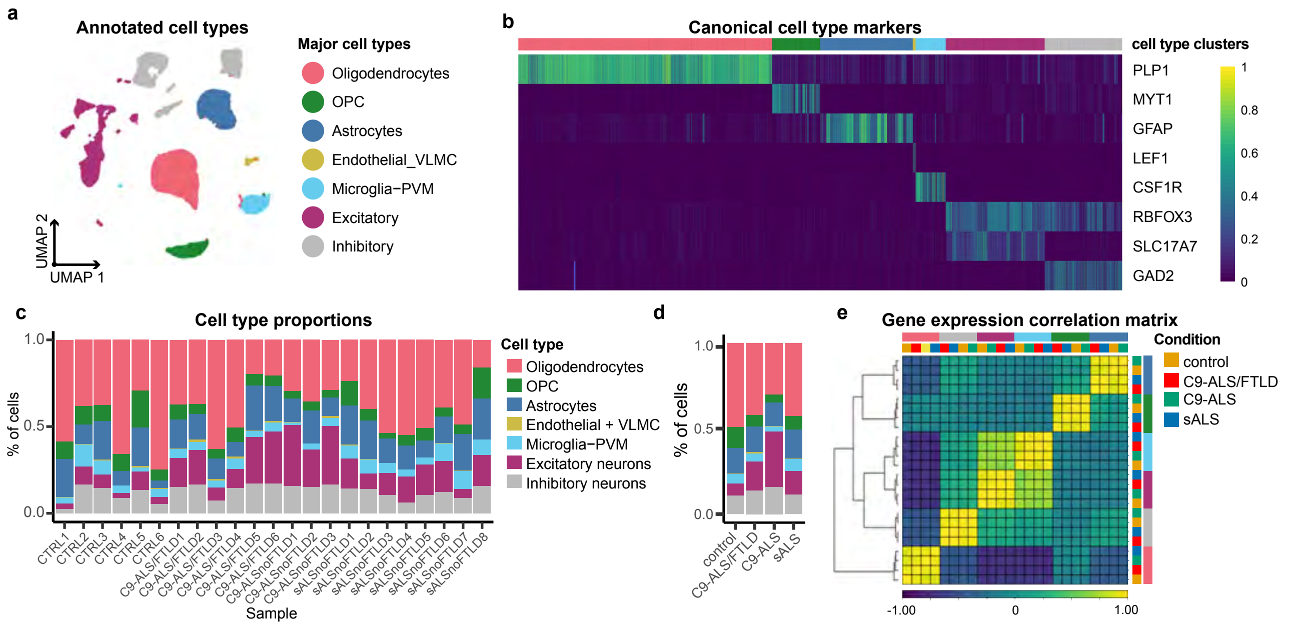


**Supplementary Fig. 2: Proportion of frontal cortex cell types across samples and conditions. (a)** UMAP plot of annotated cell types classified as major cell types. Excitatory neurons can be further subdivided into upper- (layers 2/3), intermediate- (layer 4), and deep- (layer 5-6) neurons. **(b)** Heatmap of major cell types labeled by canonical cell type markers: PLP1 (Oligodendrocytes), MYT1 (OPC), GFAP (Astrocytes), LEF1 (Endothelial-VLMC), CSF1 (Microglia-PVM), neurons (RBFOX3), excitatory neurons (SLC17A7), and inhibitory neurons (GAD2). **(c)** Barplot of cell type proportions across samples. **(d)** Barplot of cell type proportions across conditions. **(e)** Gene expression correlation matrix across celltypes and conditions. Colour scale is represented as a z-score.

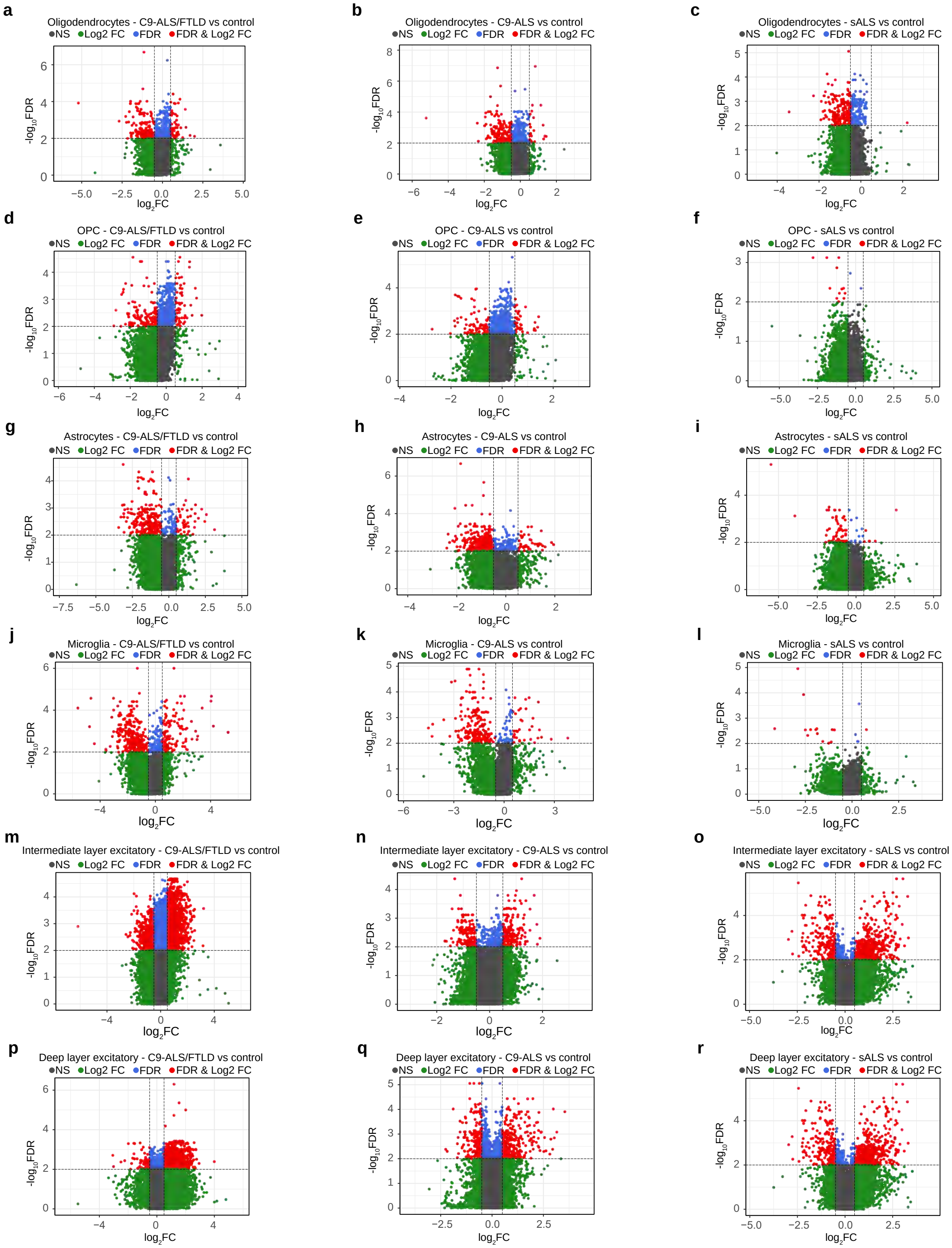


**Supplementary Fig. 3: Differentially expressed genes across glial and neuronal cell types in ALS.** Volcano plots of differentially expressed genes for either C9-ALS/FTLD, C9-ALS, or sALS (respectively) in **(a-c)** oligodendrocytes; **(d-f)** OPC; **(g-i)** astrocytes; **(j-l)** microglia; **(m-o)** intermediate-layer excitatory neurons; and **(p-r)** deep-layer excitatory neurons. Only differentially expressed genes which passed the FDR $< 0.01$ and $|\mathrm{LFC}| < 0.5$ cutoff were considered significant (shown in red).

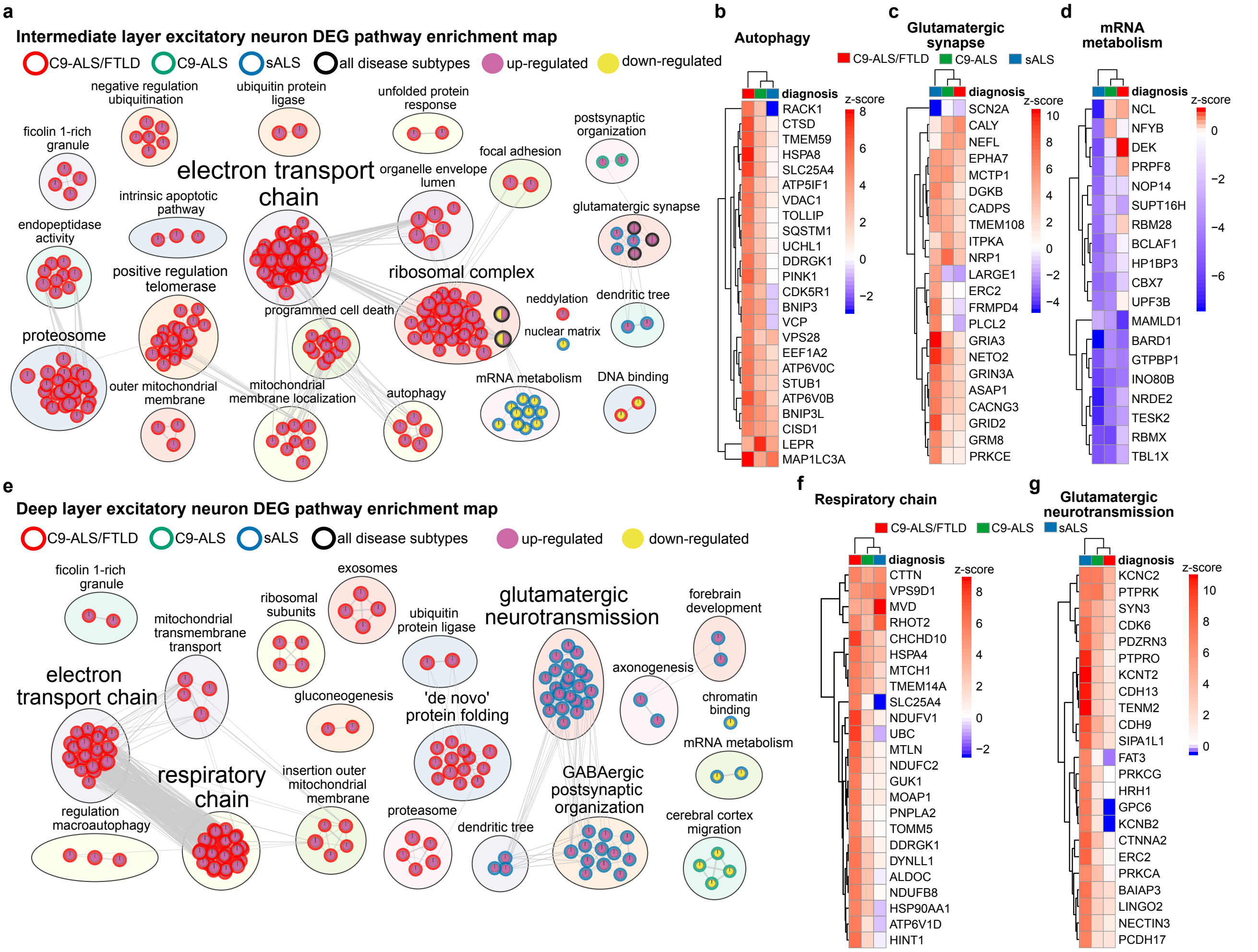


**Supplementary Fig. 4: Intermediate- and deep-layer excitatory neurons pathways alterations in ALS. (a)** Enrichment map of altered pathways in intermediate-layer excitatory neurons across ALS subtypes. Clustered heatmaps for genes identified in the **(b)** autophagy; **(c)** glutamatergic synapses; and **(d)** mRNA metabolism clusters from **(a)**. **(e)** Enrichment map of altered pathways in deep-layer excitatory neurons across ALS subtypes. Clustered heatmaps for genes identified in the **(f)** respiratory chain and **(g)** glutamatergic synapses. DEG = differentially expressed genes. Significance set at FDR $< 0.01$ and $|LFC| < 0.5$.

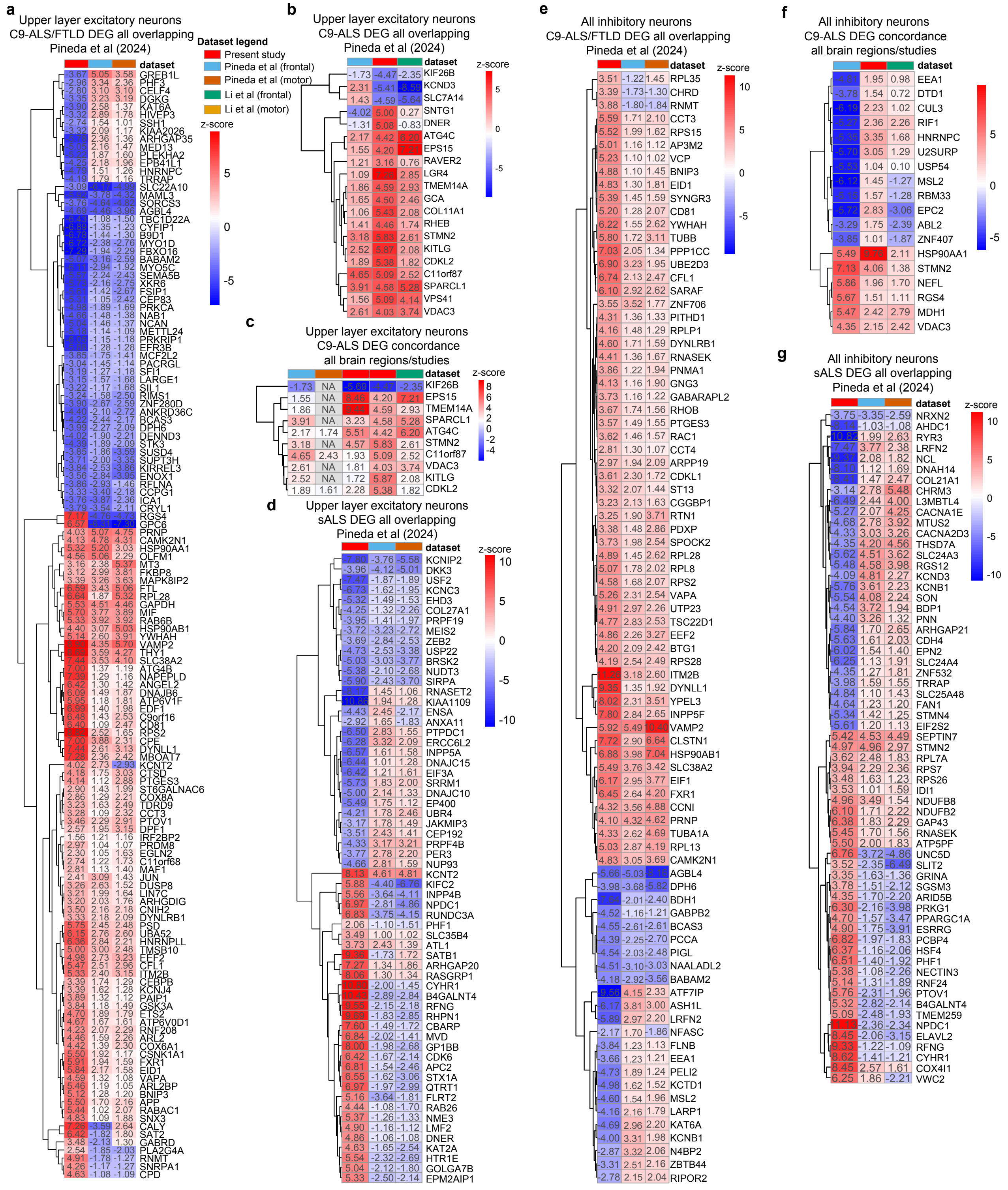


**Supplementary Fig. 5: Distribution of all concordant and discordant differentially expressed genes in cortical neurons in ALS across three independent datasets.** Clustered heatmaps of z-scores for intersecting genes shared between upper layer excitatory neurons in **(a)** frontal and motor areas in C9-ALS/FTLD[32]; **(b)** frontal areas alone in C9-ALS[31,32]; **(c)** frontal and motor regions in C9-ALS[31,32]; and frontal and motor cortex in sALS[32]. (e-g) Same as **(a,b,d)** in inhibitory neurons. Significance set at FDR < 0.01 and |LFC| < 0.5.

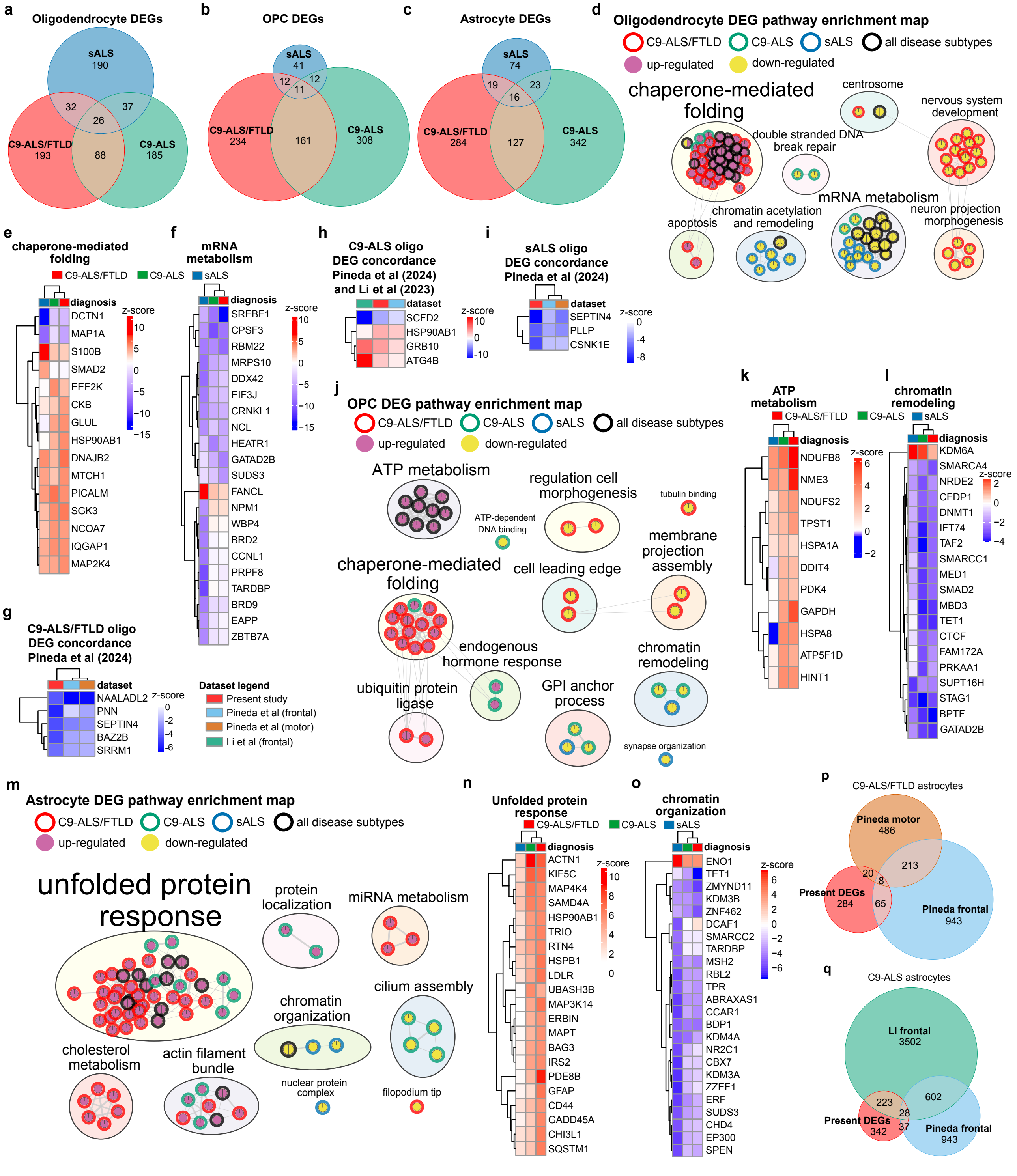


**Supplementary Fig. 6: Glial differentially expressed genes highlight protein folding, mRNA metabolism, and chromatin remodeling in ALS.** Venn diagrams indicating overlapping differentially expressed genes between C9-ALS/FTLD, C9-ALS, and sALS in **(a)** oligodendrocytes; **(b)** OPC; and **(c)** astrocytes. **(d)** Enrichment map of pathway terms enriched across ALS in oligodendrocytes. The node border indicates which ALS subtype is enriched, whereas the fill shows whether the pathway is enriched or depleted. The annotation text size for clusters is scaled by the number of nodes within each cluster. See methods for statistical parameters. Clustered heatmaps for genes identified in the **(e)** chaperone-mediated folding and **(f)** mRNA metabolism clusters in **(d)**. **(g)** Clustered heatmap of concordant C9-ALS/FTLD differentially expressed genes (positive correlation between z-scores), separated by up-regulated (left) and down-regulated (right) genes. **(h)** Clustered heatmap of concordant oligodendrocyte differentially expressed genes in C9-ALS frontal regions alone. **(i)** Clustered heatmap of concordant differentially expressed genes in sALS frontal regions alone. **(j)** Enrichment map of pathway terms enriched across ALS in OPC. Clustered heatmaps for genes identified in the **(k)** ATP metabolism and **(l)** chromatin remodeling clusters in **(j)**. **(m)** Enrichment map of pathway terms enriched across ALS in astrocytes. Clustered heatmaps for genes identified in the **(n)** unfolded protein response and **(l)** chromatin remodeling clusters in **(m)**. **(p)** Venn diagram of concordant C9-ALS/FTLD differentially expressed genes in frontal and motor region astrocytes[32] in ALS. **(q)** Venn diagram of concordant C9-ALS differentially expressed genes in frontal region astrocytes alone[31,32] in ALS. Significance set at FDR $< 0.01$ and $|LFC| < 0.5$.

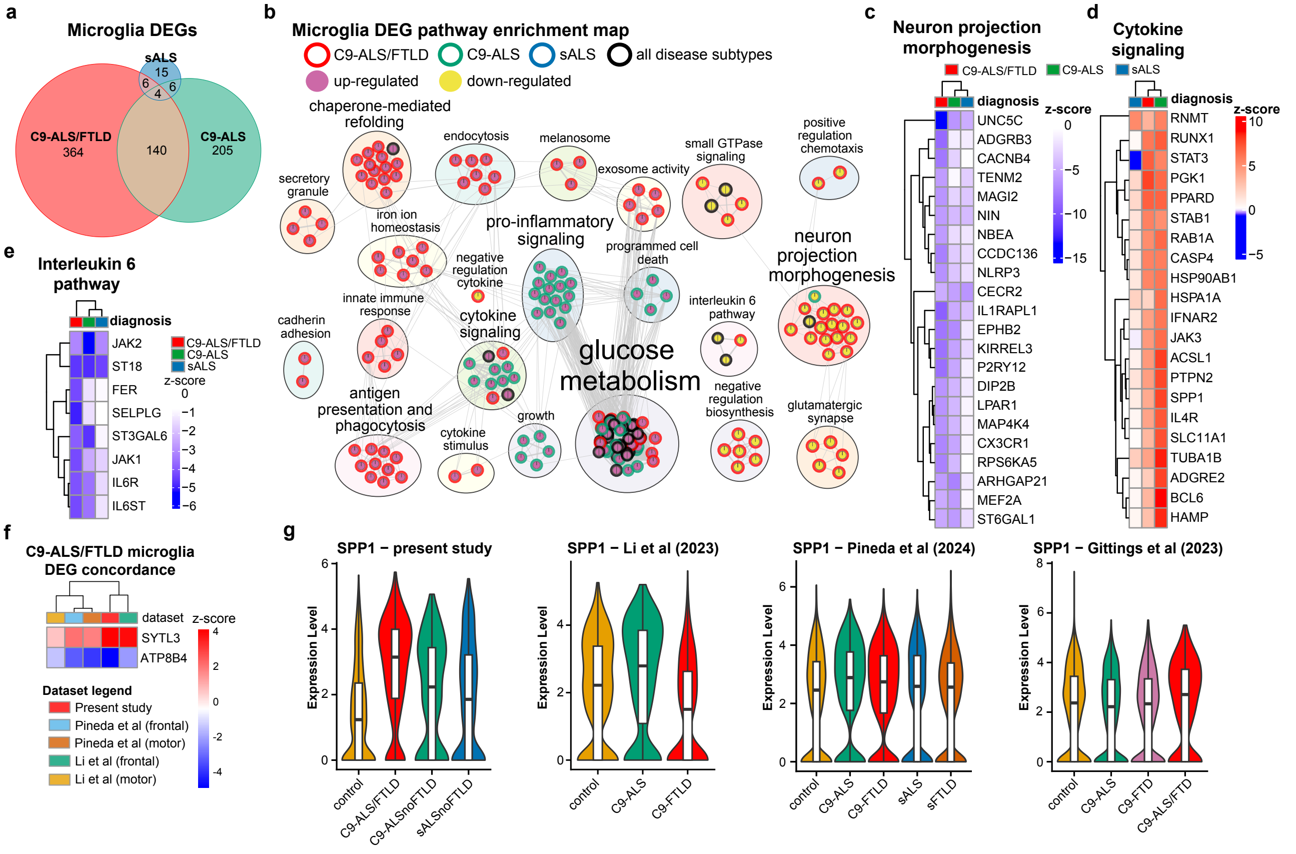


**Supplementary Fig. 7: Microglia differentially expressed genes show more pronounced pathways alterations in C9-ALS than in sALS.** **(a)** Venn diagram showing overlapping microglial differentially expressed genes between C9-ALS/FTLD, C9-ALS, and sALS. **(b)** Enrichment map of altered pathways in microglia across ALS subtypes. The node border indicates which ALS subtype is enriched, whereas the fill shows whether the pathway is enriched or depleted. The annotation text size for clusters is scaled by the number of nodes within each cluster. See methods for statistical parameters. Clustered heatmaps for genes identified in the **(c)** neuron projection morphogenesis; **(d)** cytokine signaling; and **(e)** interleukin 6 clusters from **(b)**. **(f)** Clustered heatmap of concordant differentially expressed genes (SYTL3 and ATP8B4) in C9-ALS/FTLD and C9-ALS in frontal and motor areas[31,32]. **(g)** Violin plots of SYTL3 expression across conditions in four independent frontal cortex datasets[31–33] Significance set at FDR $< 0.01$ and $|LFC| < 0.5$.

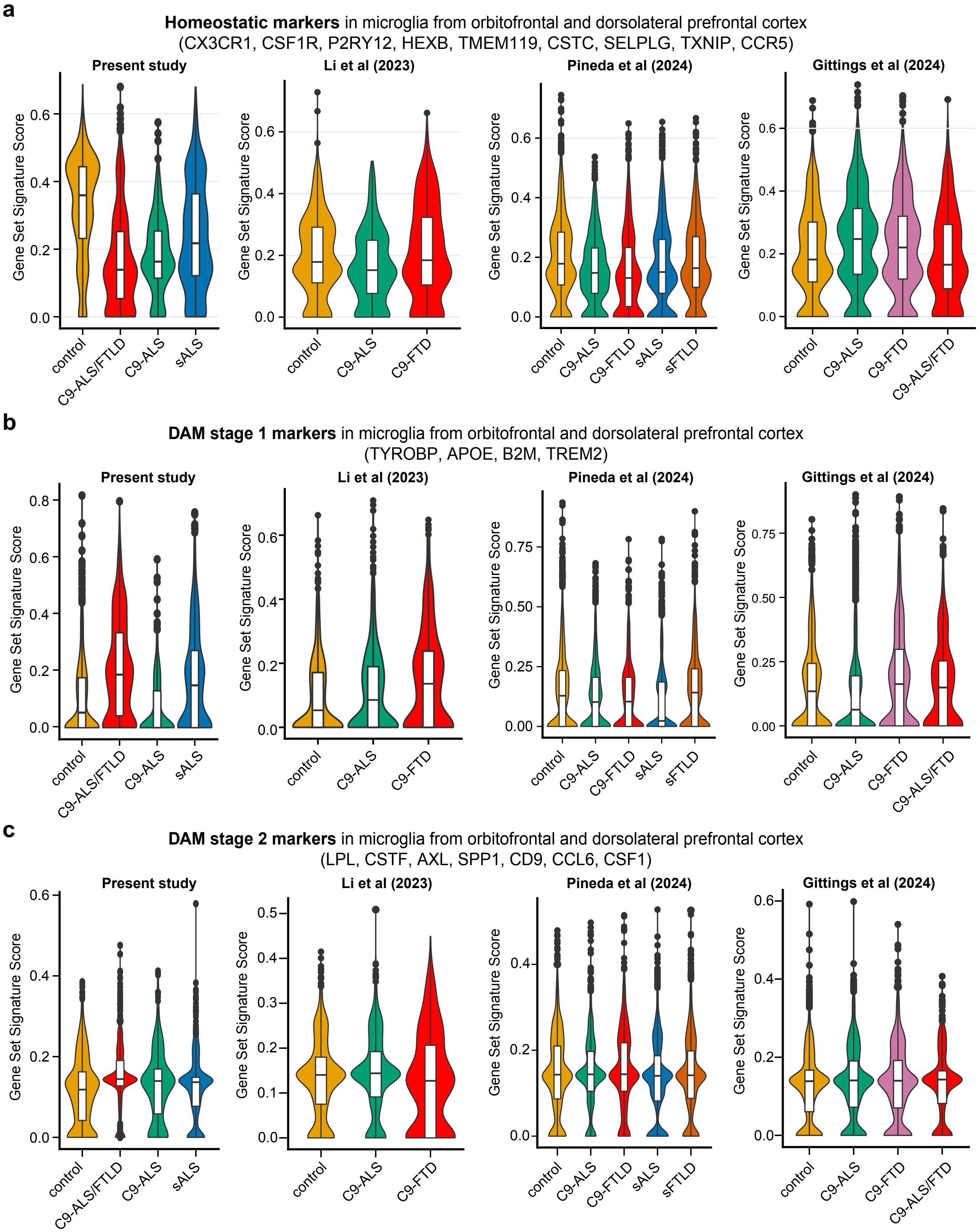


**Supplementary Fig. 8: Microglia differentially expressed genes show loss of homeostatic markers and increases in neurodegenerative DAM markers.** **(a)** Violin and barplots of gene signature scores across four independent datasets[31–33] for **(h)** homeostatic microglial markers (CX3CR1, CSF1R, P2RY12, HEXB, TMEM119, CSTC, SELPLG, TXNIP, CCR5); **(b)** disease associated microglia (DAM) stage 1 (TYROBP, APOE, B2M, TREM2); and **(c)** DAM stage 2 (LPL, CSTF, AXL, SPP1, CD9, CCL6, CSF1). Significance set at FDR $< 0.01$ and $|LFC| < 0.5$.

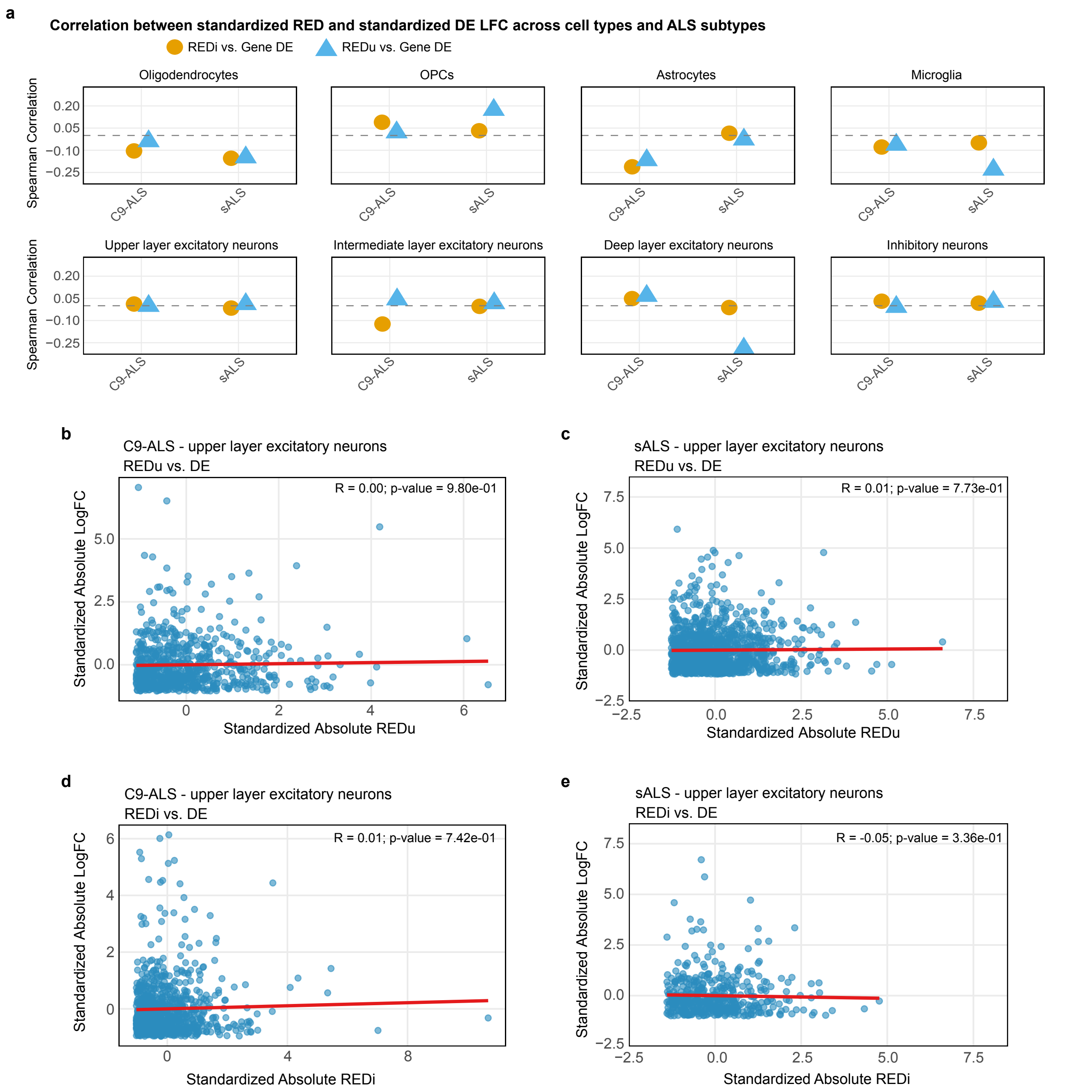


**Supplementary Fig. 9: APA changes are not correlated with differential expression in C9-ALS or sALS.** **(a)** Spearman correlation between standardized differential expression (DE) LFC and standardized relative expression differences (RED) across cell types. Orange dots show correlation between REDi (intronic vs. 3′UTR APA) and DE, while blue triangles show correlation between REDu (3′UTR APA) and DE. **(b-e)** Representative scatter plots from upper layer excitatory neurons showing relationship between standardized DE LFC and standardized absolute REDu **(b,c)** or REDi **(d,e)** in C9-ALS **(b,d)** and sALS **(c,e)**. Spearman correlation coefficients (R) and p-values are shown. The lack of strong correlation suggests that APA regulation occurs independently of changes in gene expression.

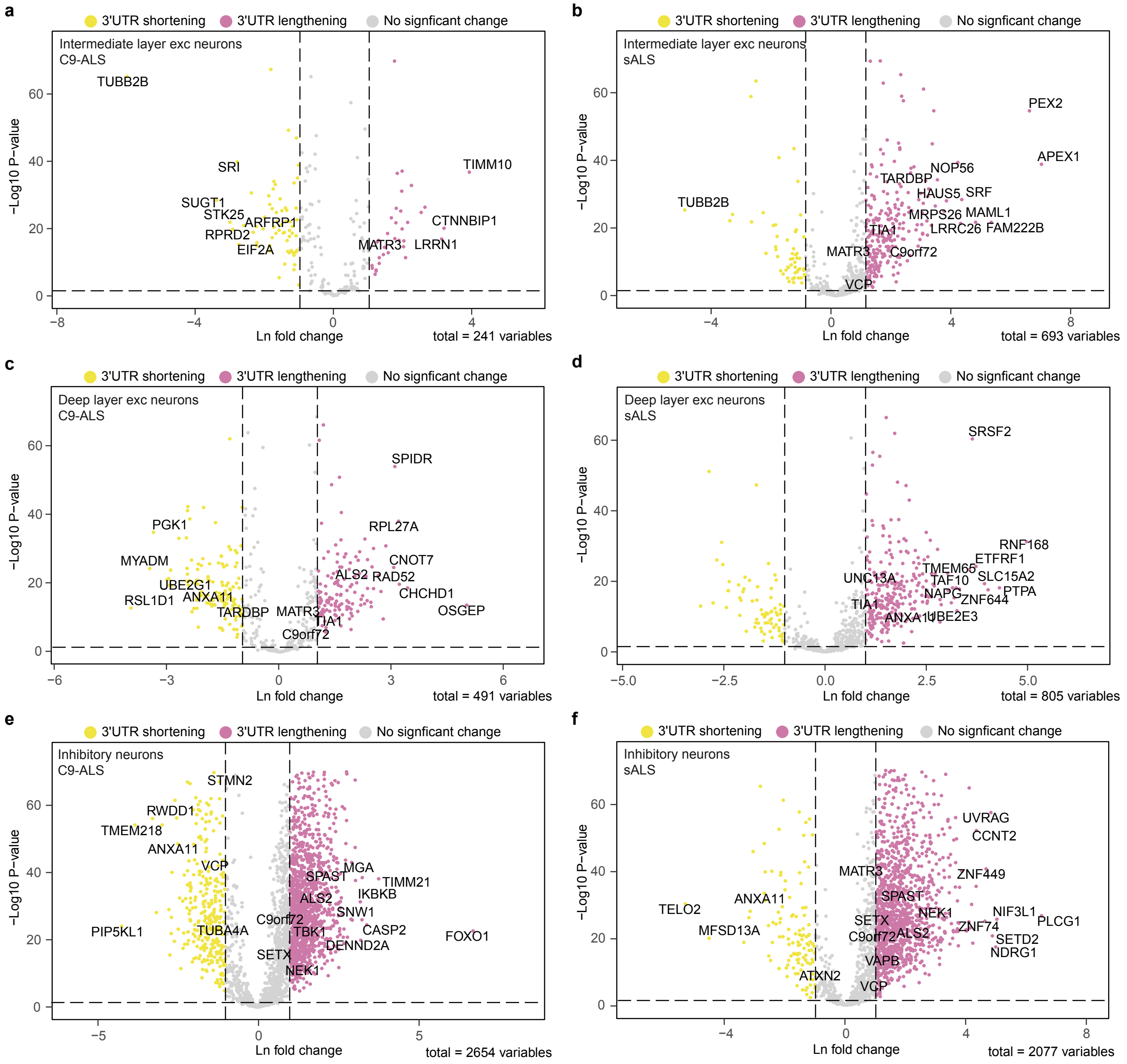


**Supplementary Fig. 10: Dysregulation of APA in ALS across neuronal cell types.** Volcano plots depicting genes undergoing 3′-UTR lengthening and shortening in ALS subtypes across inhibitory neurons in **(a)** C9-ALS and **(b)** sALS; upper layer excitatory neurons in **(c)** C9-ALS and **(d)** sALS; intermediate-layer excitatory neurons in **(e)** C9-ALS and **(f)** sALS; and deep-layer excitatory neurons in **(g)** C9-ALS and **(h)** sALS;.

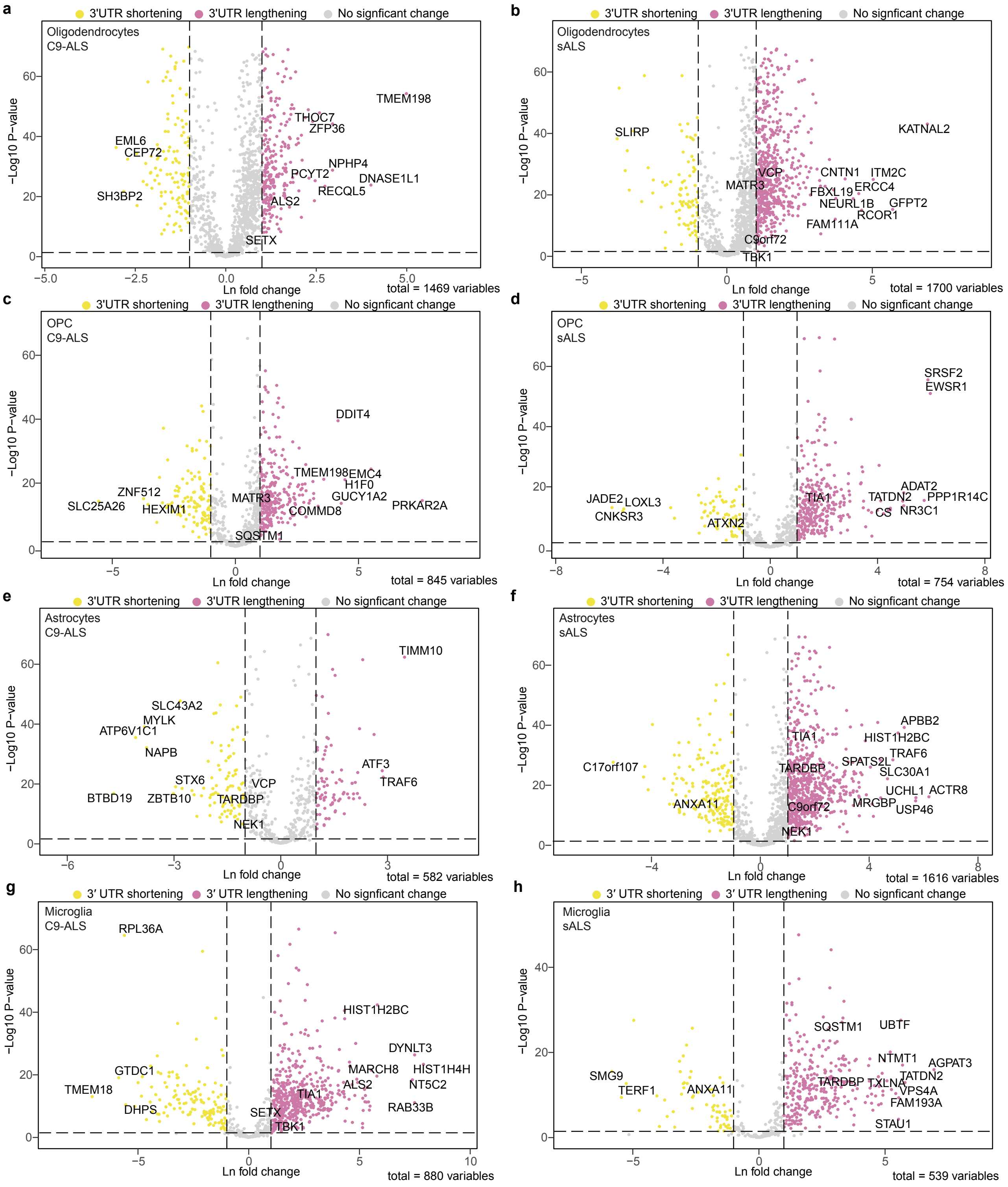


**Supplementary Fig. 11: Dysregulation of APA in ALS across glial cell types.** Volcano plots depicting genes undergoing 3′-UTR lengthening and shortening in ALS subtypes across oligodendrocytes in **(a)** C9-ALS and **(b)** sALS; OPCs in **(c)** C9-ALS and **(d)** sALS; astrocytes in **(e)** C9-ALS and **(f)** sALS, and microglia in **(g)** C9-ALS and **(h)** sALS.

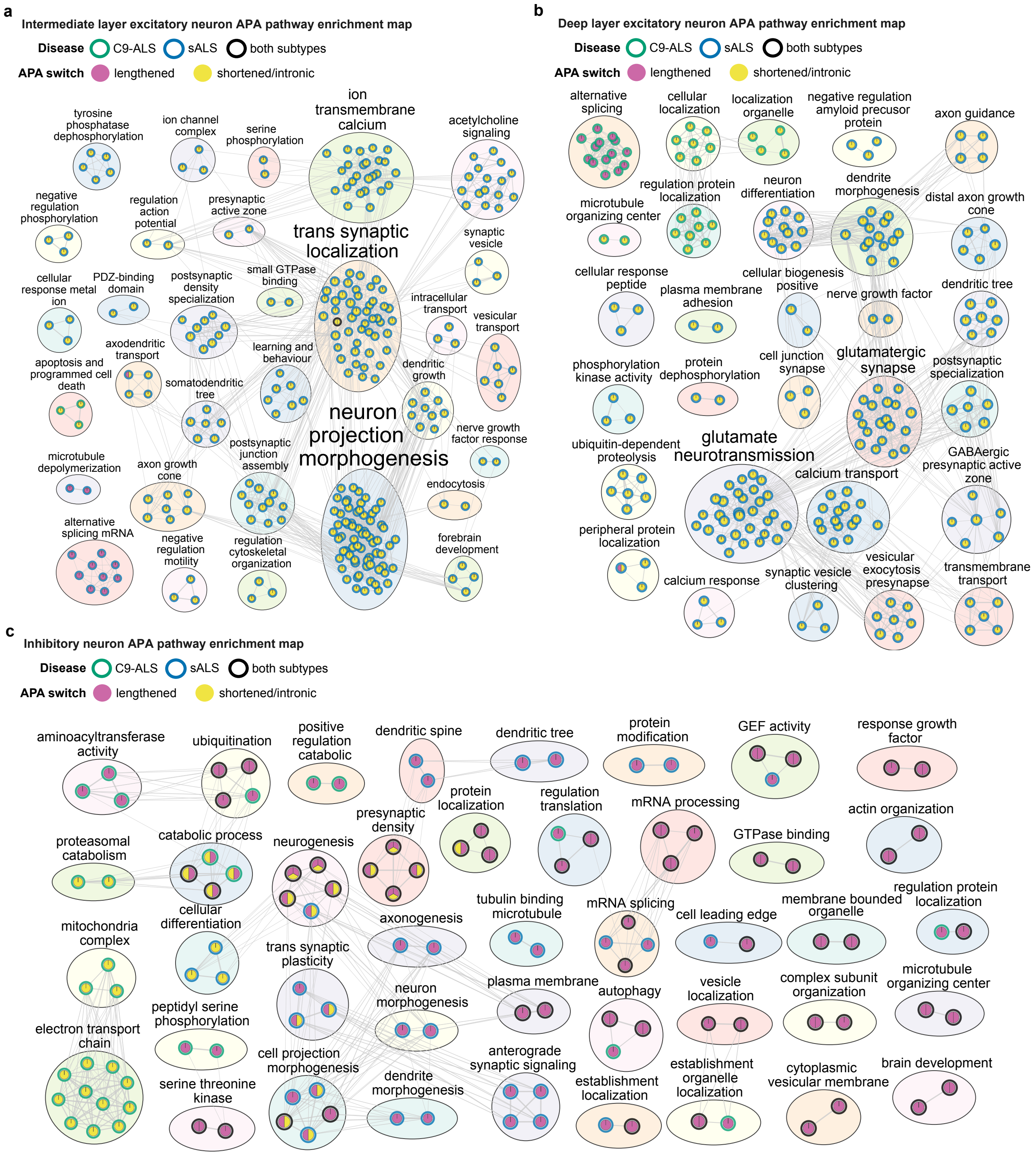


**Supplementary Fig. 12: Pathways enriched in APA events in neurons.** Enrichment map of significantly altered pathways in **(a)** upper layer excitatory neurons; **(b)** intermediate-layer excitatory neurons; and **(c)** deep-layer excitatory neurons across ALS subtypes. The node border indicates which ALS subtype is enriched, whereas the fill shows whether the pathway is enriched for transcripts with lengthened 3′-UTR or shortened 3′-UTR and/or intronic APA transcripts. The annotation text size for clusters is scaled by the number of nodes within each cluster. See methods for statistical parameters.

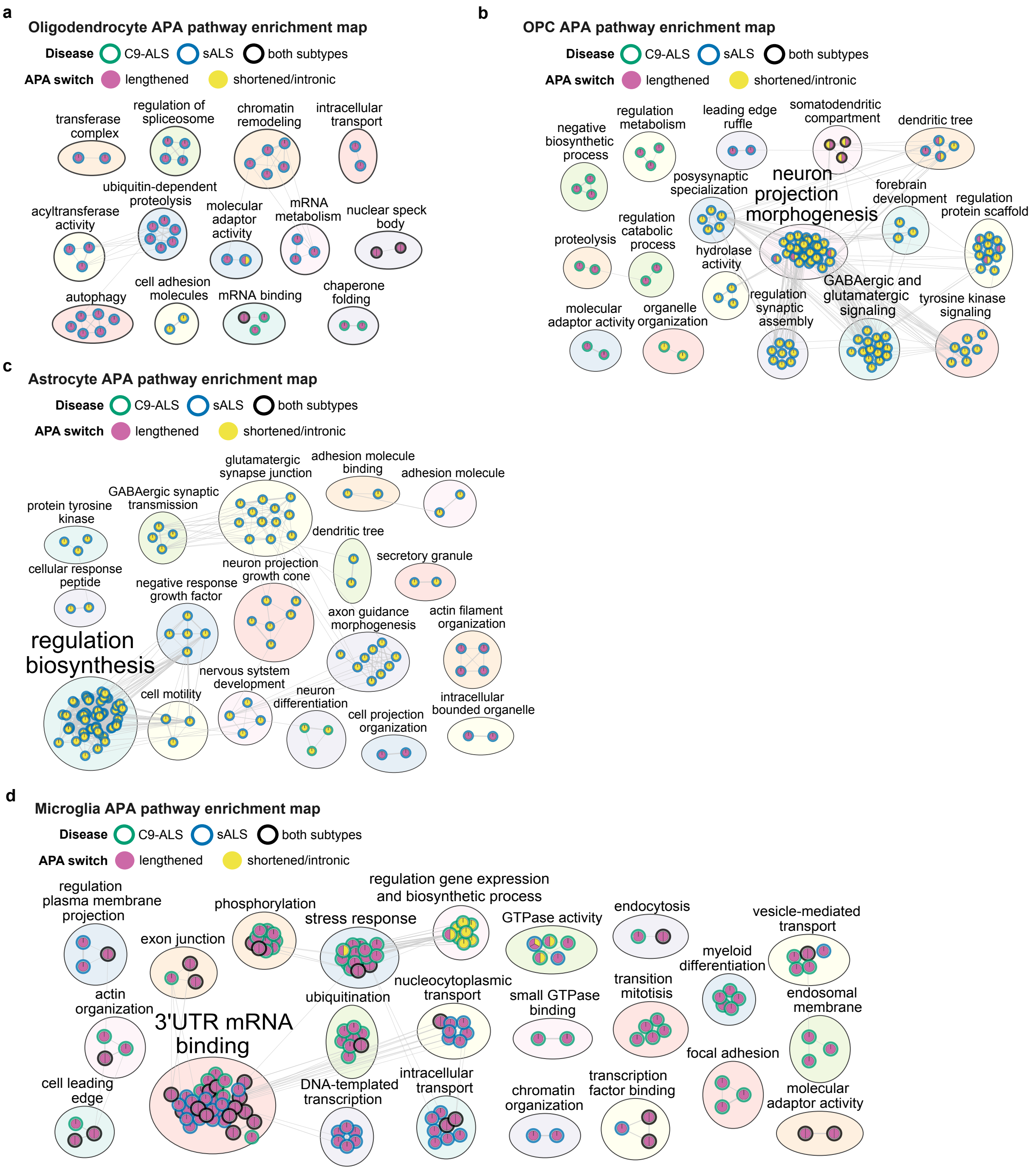


**Supplementary Fig. 13: Pathways enriched in APA events in glial cells and inhibitory neurons.** Enrichment map of significantly altered pathways in **(a)** oligodendrocytes; **(b)** OPCs; and **(c)** deep-layer excitatory neurons across ALS subtypes. The node border indicates which ALS subtype is enriched, whereas the fill shows whether the pathway is enriched for transcripts with lengthened 3′-UTR or shortened 3′-UTR and/or intronic APA transcripts. The annotation text size for clusters is scaled by the number of nodes within each cluster. See methods for statistical parameters.

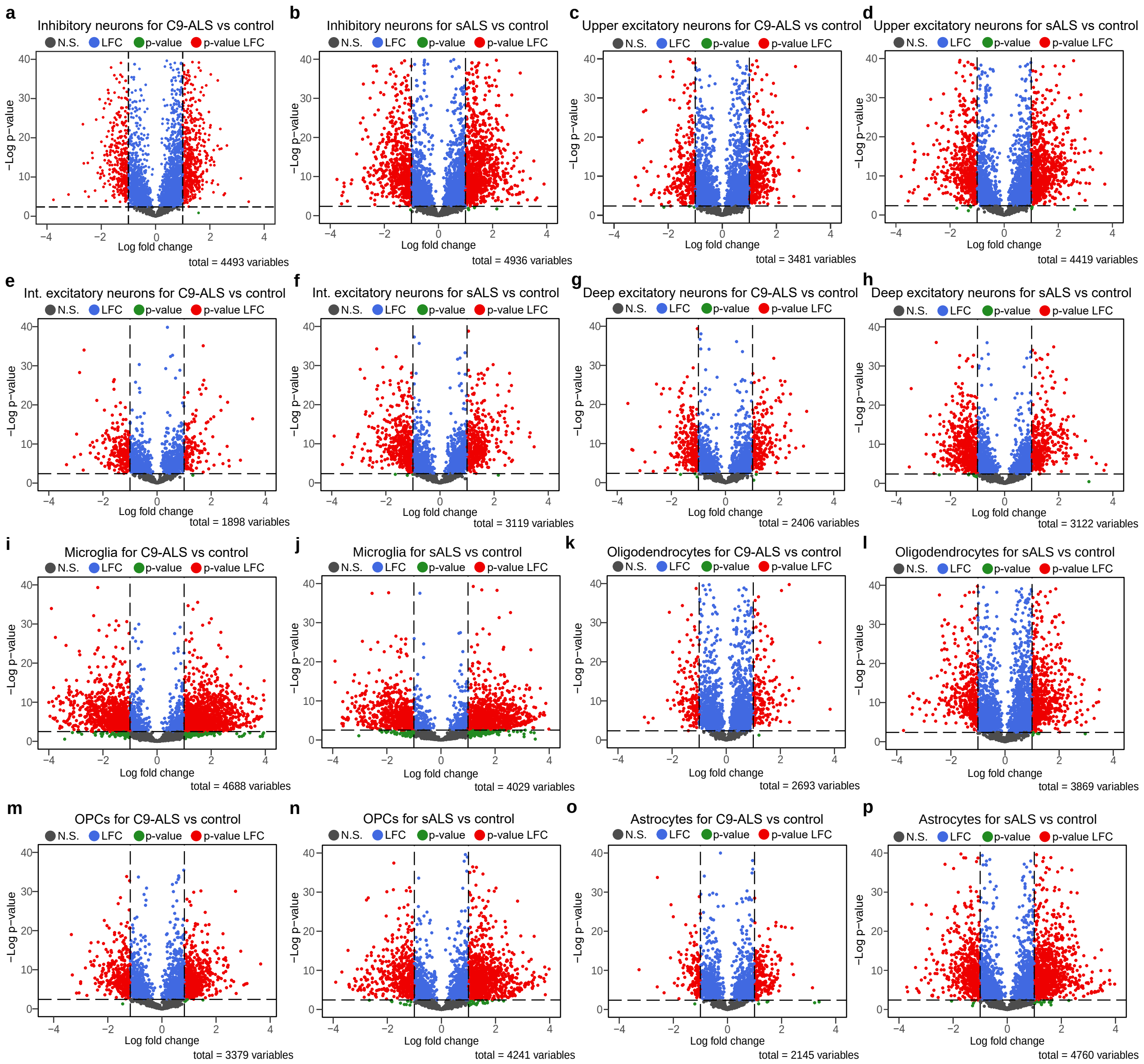


**Supplementary Fig. 14: APA events charting all feasible APA pairs between proximal and distal PA sites in all transcripts with significant APA. (a-o)** Volcano plots of APA events derived from APAlog for either C9-ALS or sALS across all cell types. The x-axis denotes the natural logarithm (Ln) fold change of distal to proximal PAs usage (see Methods). APA events are shown only for transcripts with at least one significant APA based on empirical FDR thresholds (see Methods).

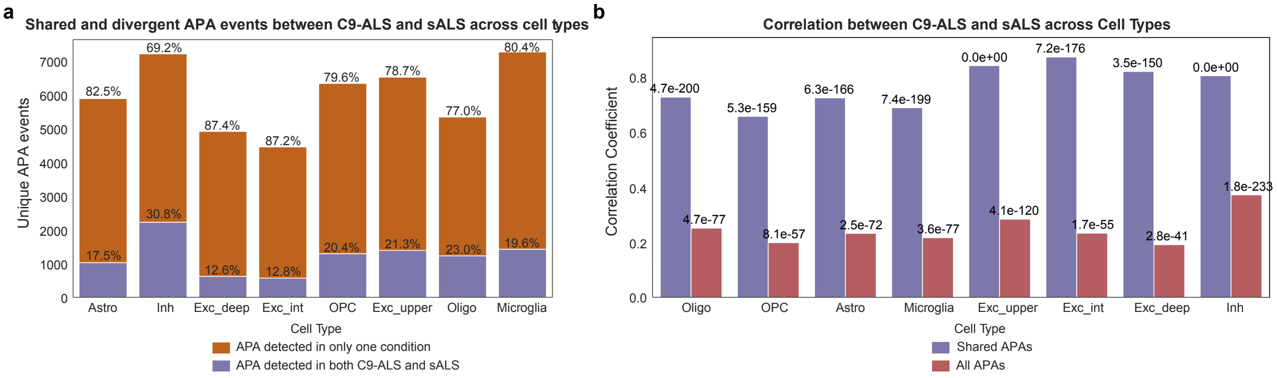


**Supplementary Fig. 15: Shared and divergent APA across the ALS subtypes (a)** Stacked bar plot showing the distribution of APA events across cell types. Purple bars represent events detected in both C9-ALS and sALS (shared events), while brown bars indicate events unique to either subtype. Percentages above each bar show the proportion of unique events. **(b)** Correlation analysis between C9-ALS and sALS across cell types. Purple bars show correlation coefficients for shared events (events present in both subtypes), while red bars represent correlation when including all events (both shared and unique events, where unique events are considered as zero in the other subtype). P-values above bars indicate statistical significance of correlations for shared events.

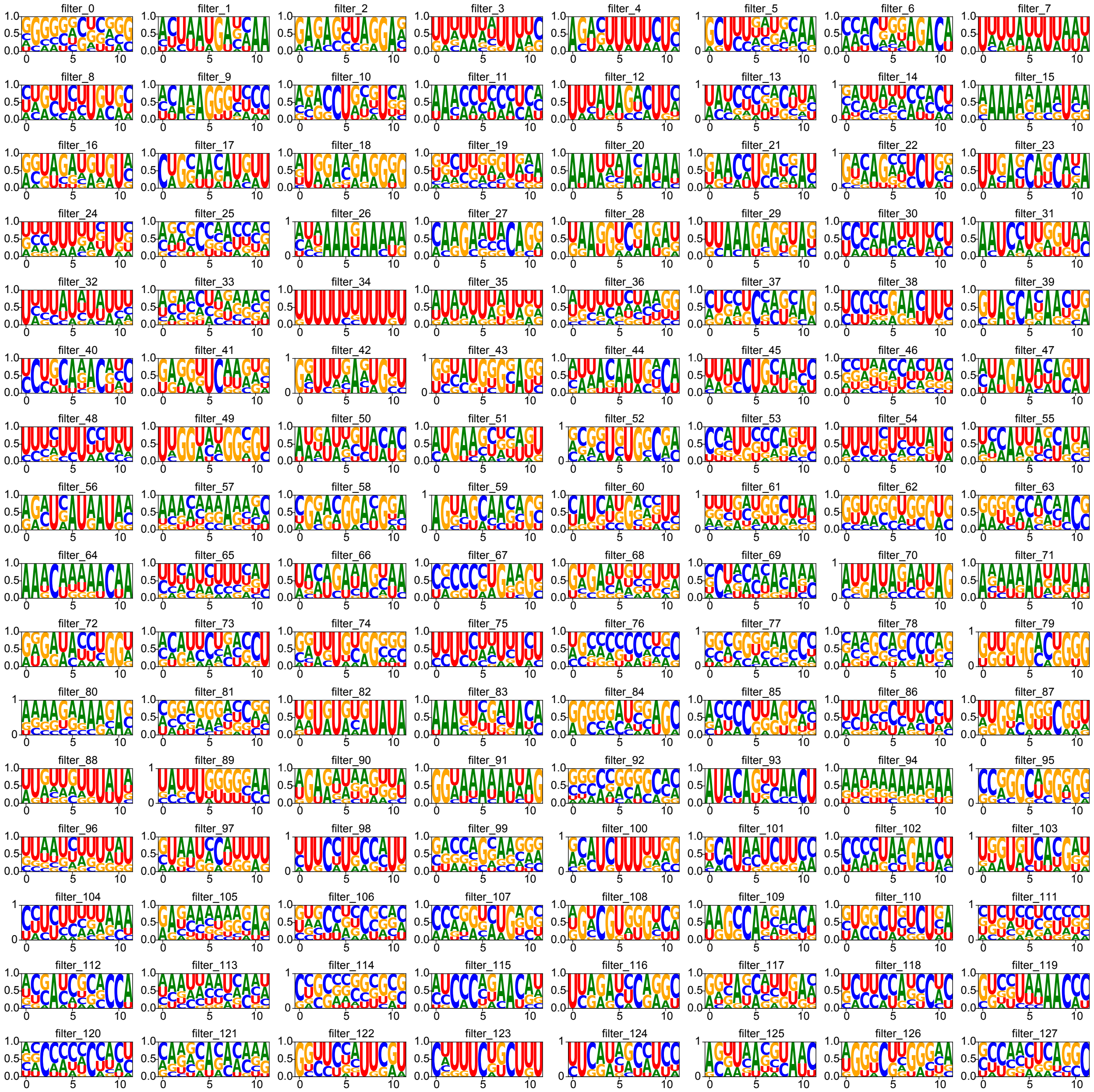


**Supplementary Fig. 16: position weight matrix (PWM) motif representation of 128 filters from the CNN module of APA-Net.** Visualization of 128 motif PWMs for both C9-ALS and sALS. PWMs are generated by scanning the test sequences and aligning the subsequences with high filter activation levels.

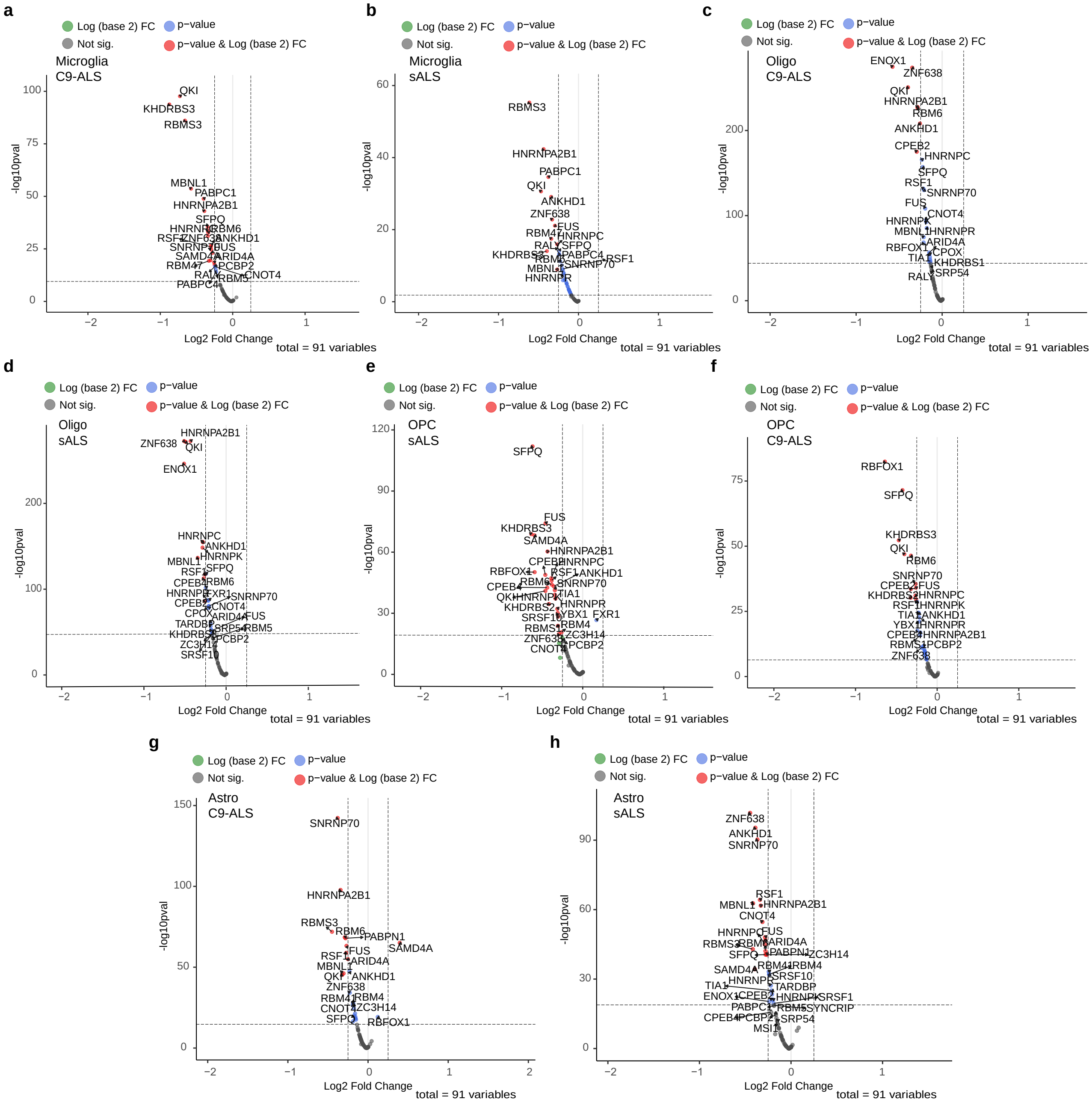


**Supplementary Fig. 17: RBPs are differentially expressed across glial cells from C9-ALS or sALS cases.** Volcano plots of a differential expression analysis on the 91 identified RBPs from APA-Net for microglia in **(a)** C9-ALS and **(b)** sALS versus controls; oligodendrocytes in **(c)** C9-ALS and **(d)** sALS; OPC in **(e)** C9-ALS and **(f)** sALS; and astrocytes in **(g)** C9-ALS and **(h)** sALS. Differential expression was performed using DESeq2 on a subsetted counts matrix consisting of only the 91 unique RBP identifiers (see Methods). Significantly altered RBPs show an adjusted p-value $< 0.01$ and a $|\log_2 \text{fold change}| < 0.25$.

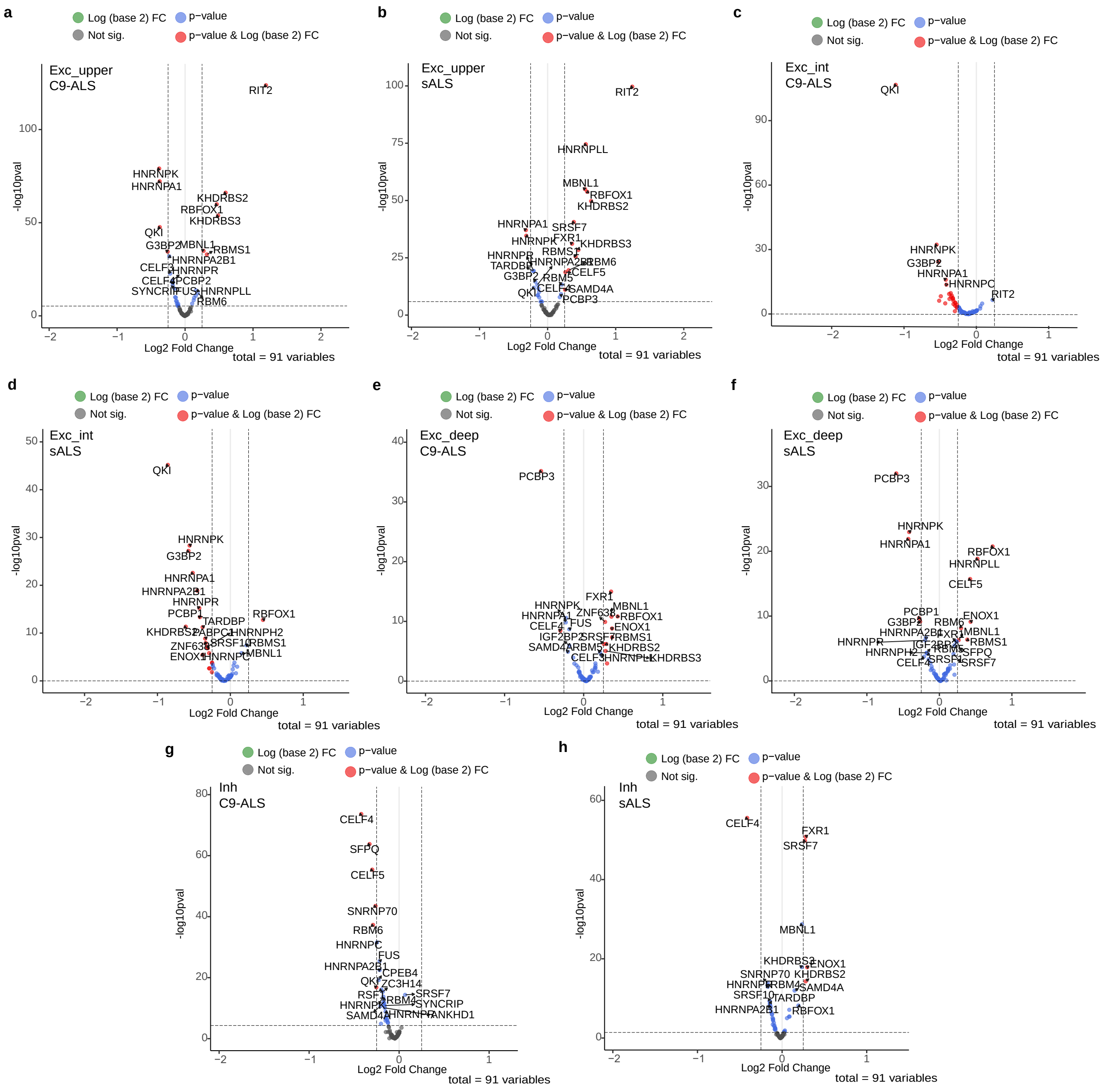


**Supplementary Fig. 18: RBP expression levels are dysregulated across neuron cell types from C9-ALS or sALS cases.** Volcano plots of a differential expression analysis on the 91 identified RBPs from APA-Net for upper layer excitatory neurons in **(a)** C9-ALS and **(b)** sALS versus controls; intermediate-layer excitatory neurons in **(c)** C9-ALS and **(d)** sALS; deep-layer excitatory neurons in **(e)** C9-ALS and **(f)** sALS; and inhibitory neurons in **(g)** C9-ALS and **(h)** sALS. Differential expression was performed using DESeq2 on a subsetted counts matrix consisting of only the 91 unique RBP identifiers (see Methods). Significantly altered RBPs showed an adjusted p-value $< 0.01$ and a $|\log_2 \text{fold change}| < 0.25$.

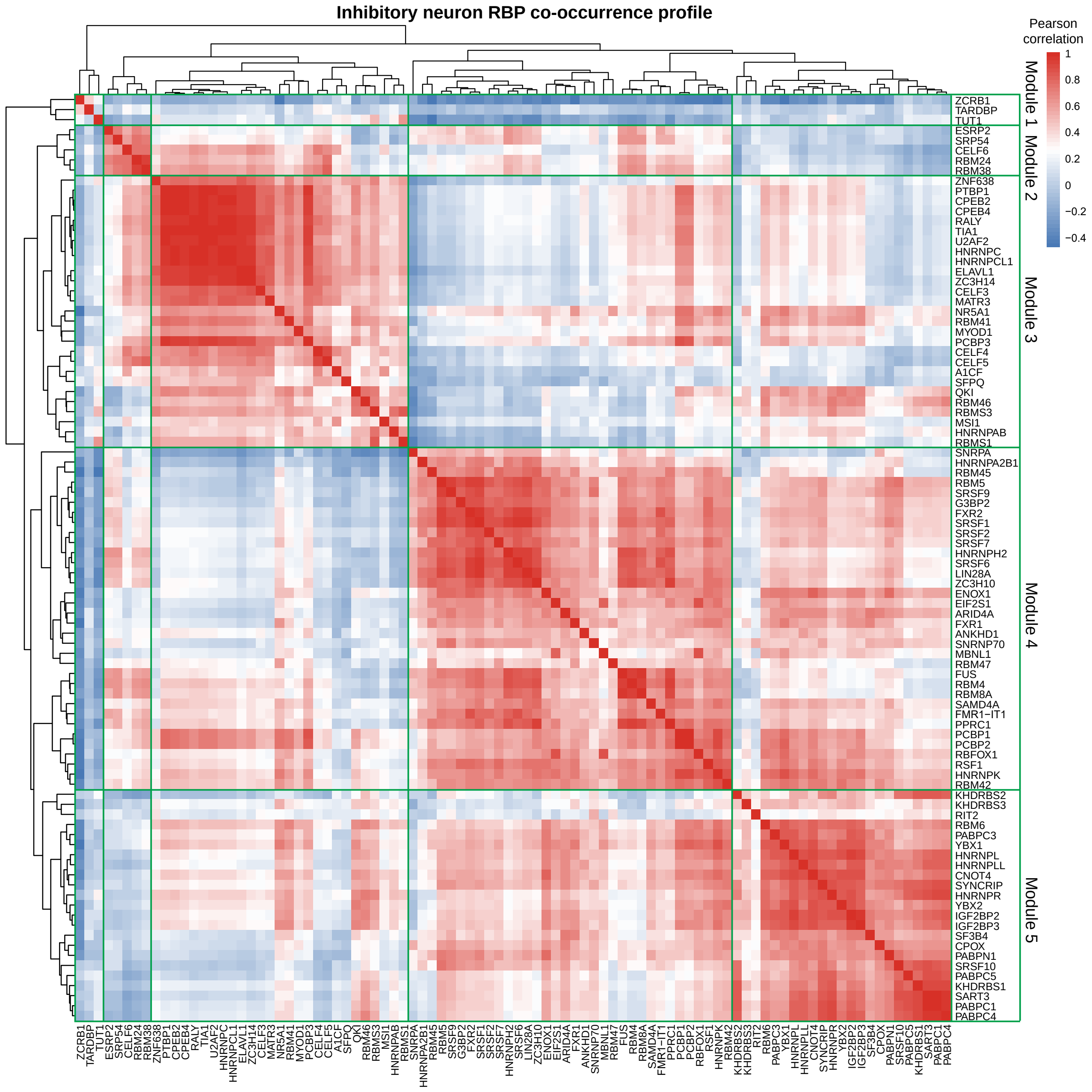


**Supplementary Fig. 19: RBP motif co-occurrence heatmap for inhibitory neurons in C9-ALS and sALS cell types.** Clustered RBP motif co-occurrence profile. The heatmap is computed using Pearson correlation of co-occurrence of RBP-aligned filters in inhibitory neuron APA events. Five RBP modules are defined based on this clustering (labeled on y- and x-axes).

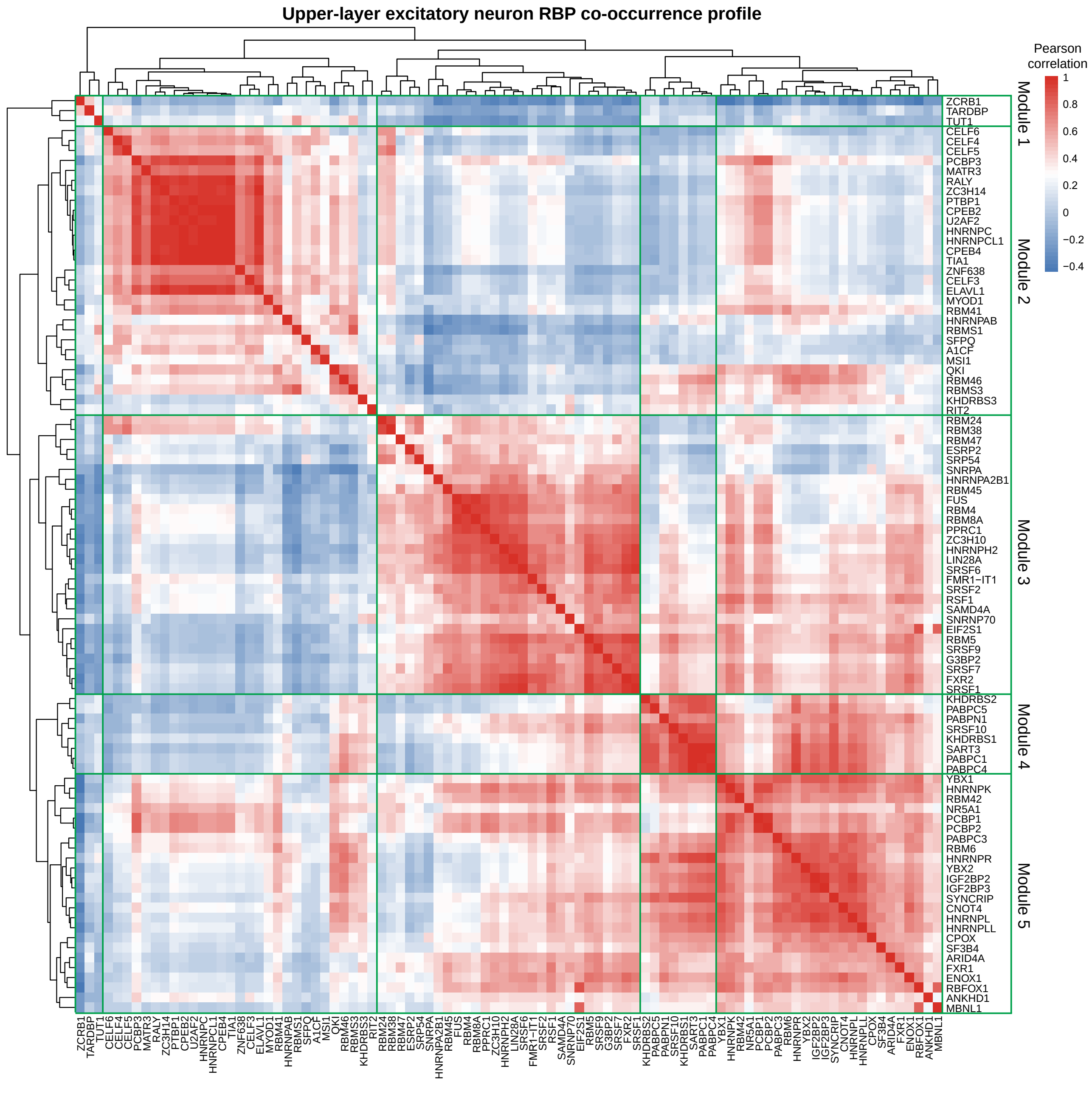


**Supplementary Fig. 20: RBP motif co-occurrence heatmap for upper layer excitatory neurons in C9-ALS and sALS cell types.** Clustered RBP motif co-occurrence profile. The heatmap is computed using Pearson correlation of co-occurrence of RBP-aligned filters in upper layer excitatory neuron APA events. Five RBP modules are defined based on this clustering (labeled on y- and x-axes).

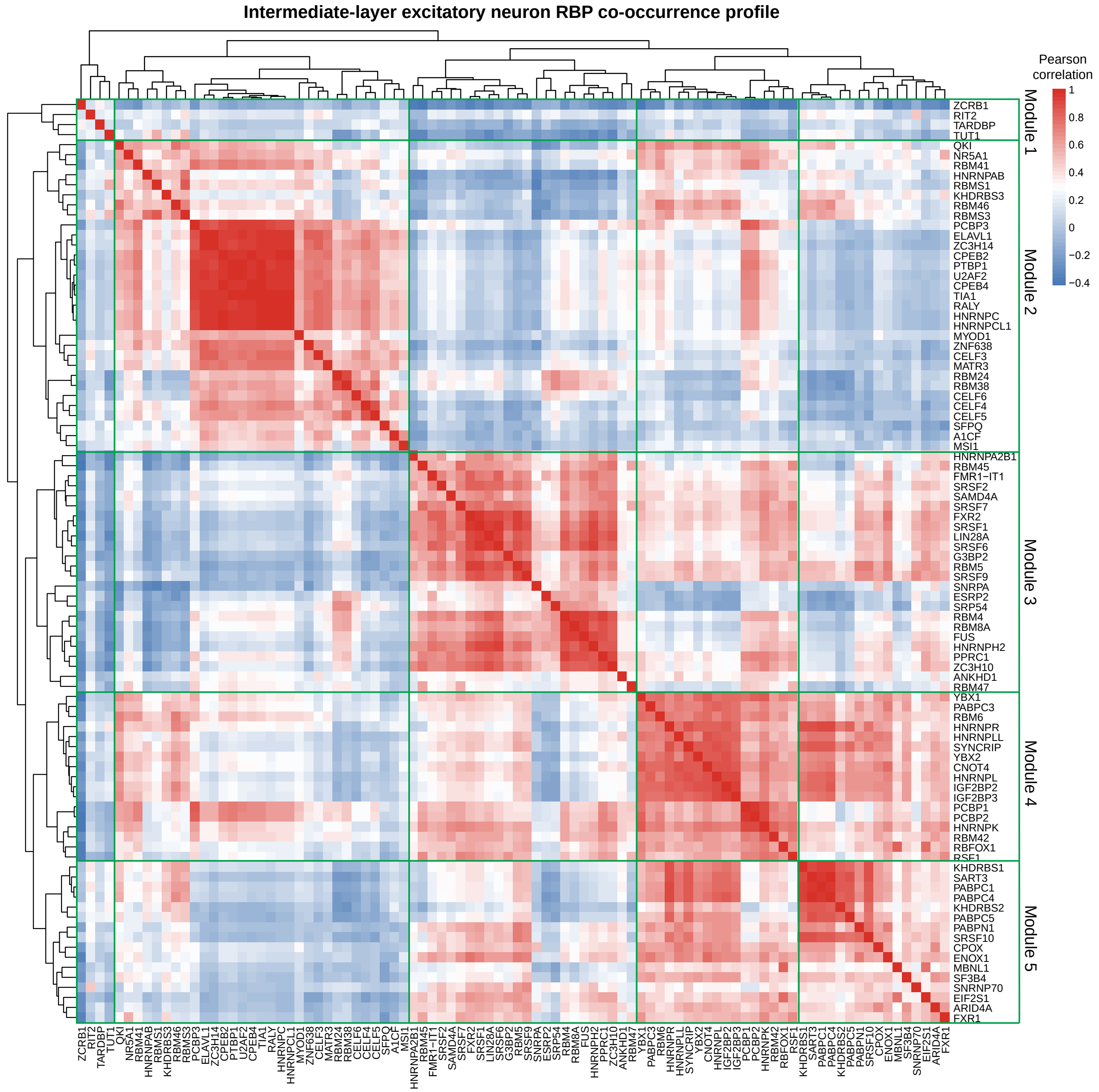


**Supplementary Fig. 21: RBP motif co-occurrence heatmap for intermediate-layer excitatory neurons in C9-ALS and sALS cell types.** Clustered RBP motif co-occurrence profile. The heatmap is computed using Pearson correlation of co-occurrence of RBP-aligned filters in intermediate-layer excitatory neuron APA events. Five RBP modules are defined based on this clustering (labeled on y- and x-axes).

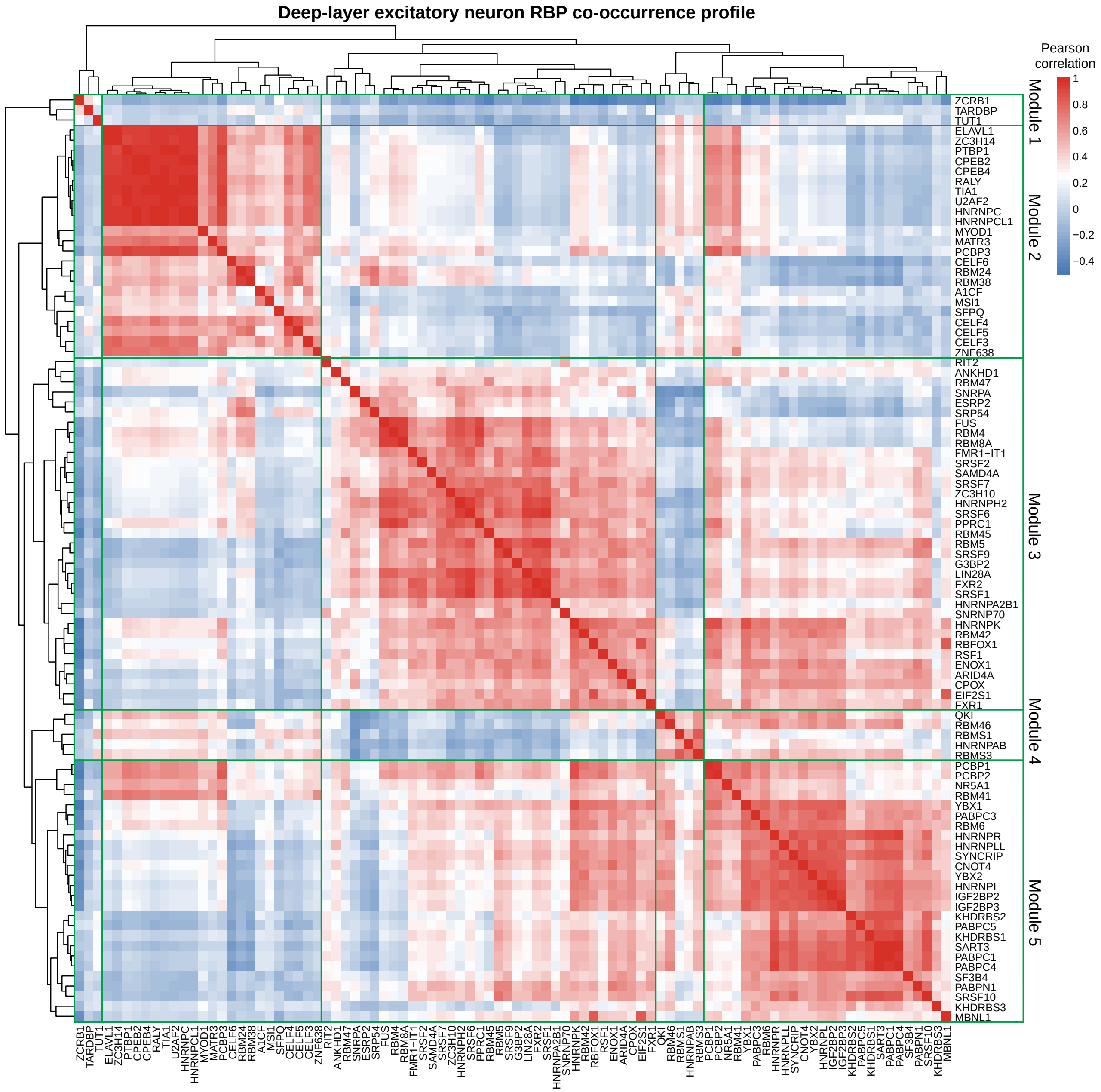


**Supplementary Fig. 22: RBP motif co-occurrence heatmap for deep-layer excitatory neurons in C9-ALS and sALS cell types.** Clustered RBP motif co-occurrence profile. The heatmap is computed using Pearson correlation of co-occurrence of RBP-aligned filters in deep-layer excitatory neuron APA events. Five RBP modules are defined based on this clustering (labeled on y- and x-axes).

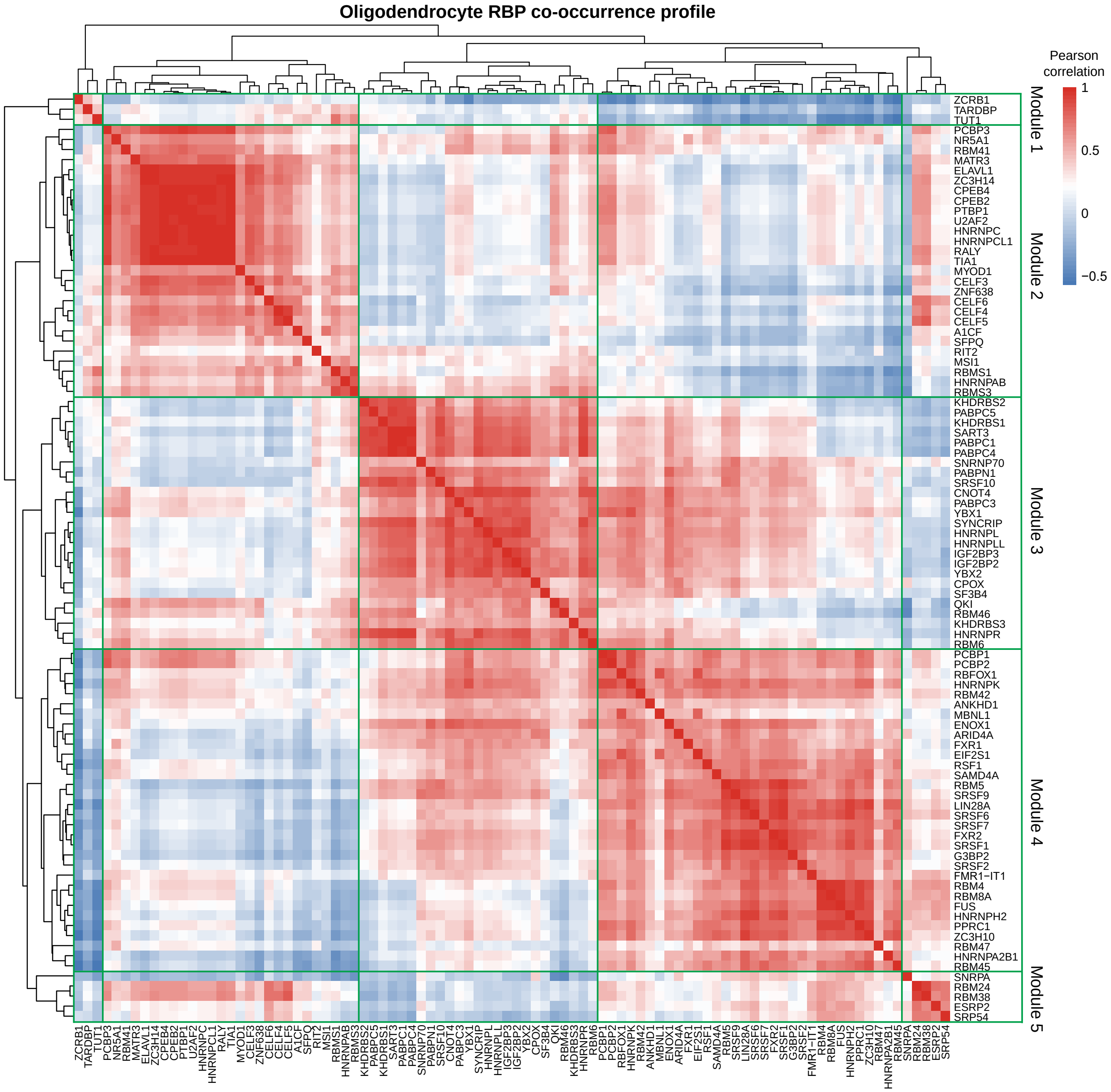


**Supplementary Fig. 23: RBP motif co-occurrence heatmap for oligodendrocytes in C9-ALS and sALS cell types.** Clustered RBP motif co-occurrence profile. The heatmap is computed using Pearson correlation of co-occurrence of RBP-aligned filters in oligodendrocyte APA events. Five RBP modules are defined based on this clustering (labeled on y- and x-axes).

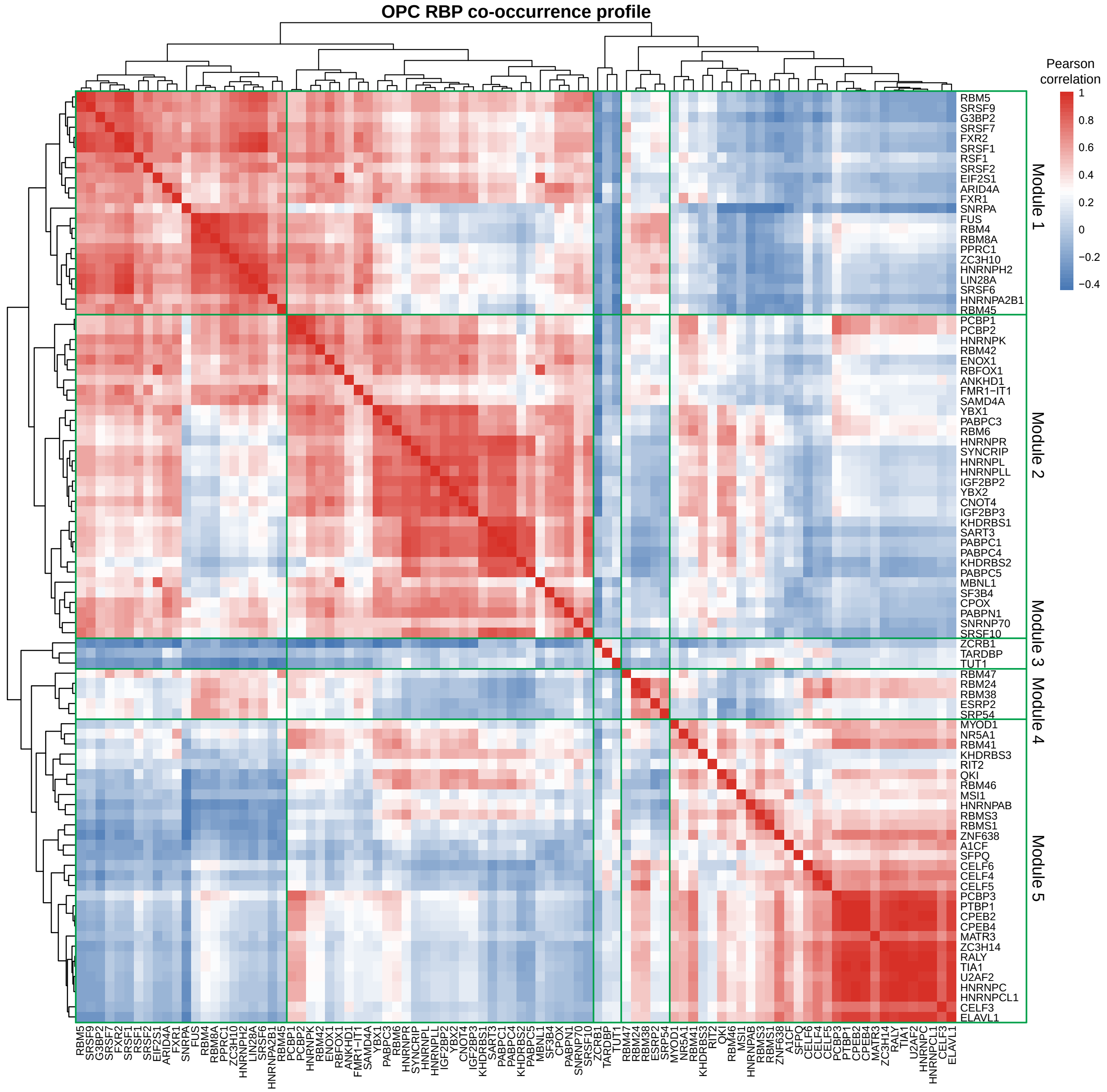


**Supplementary Fig. 24: RBP motif co-occurrence heatmap for OPCs in C9-ALS and sALS cell types.** Clustered RBP motif co-occurrence profile. The heatmap is computed using Pearson correlation of co-occurrence of RBP-aligned filters in OPC APA events. Five RBP modules are defined based on this clustering (labeled on y- and x-axes).

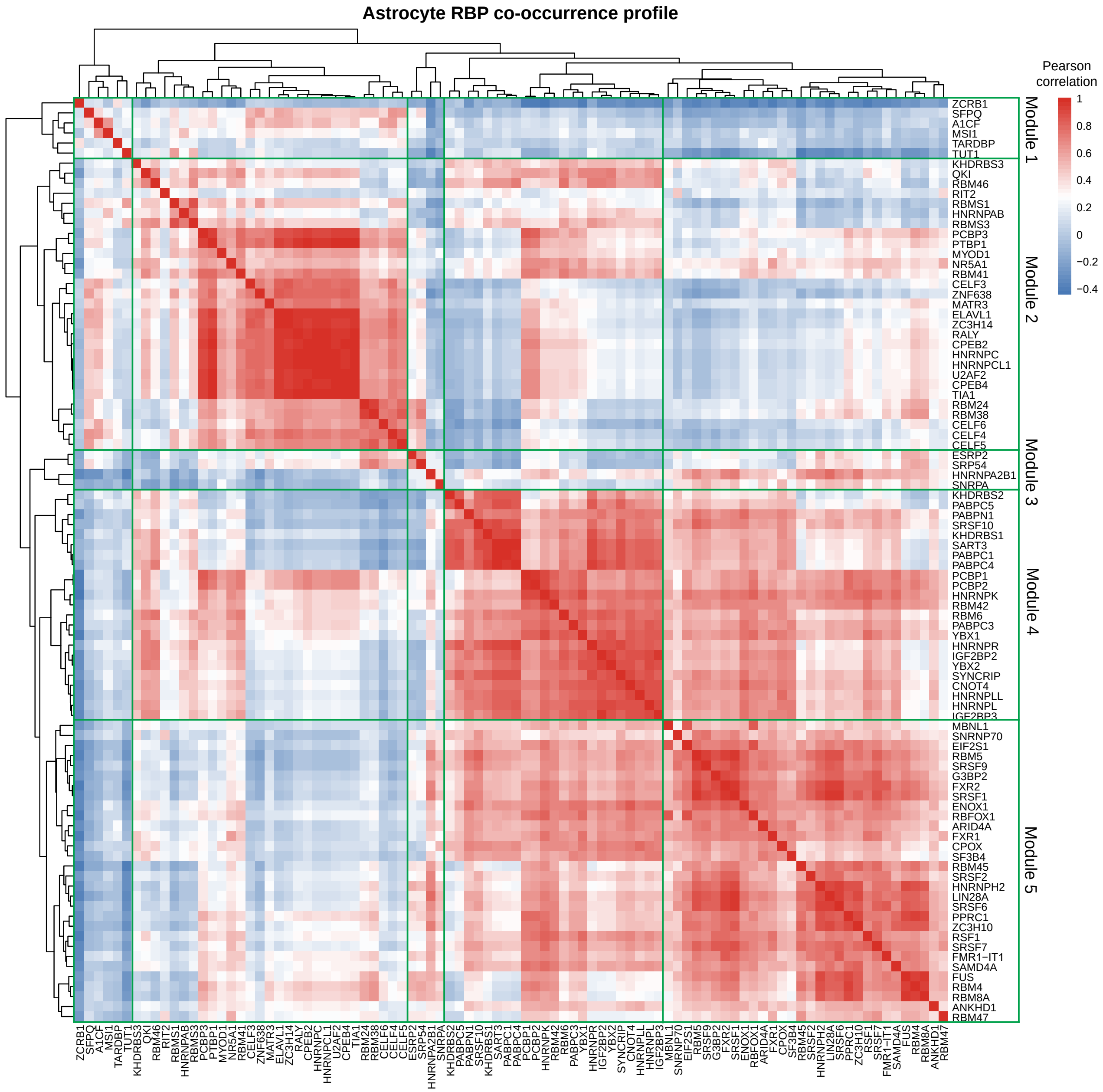


**Supplementary Fig. 25: RBP motif co-occurrence heatmap for astrocytes in C9-ALS and sALS cell types.** Clustered RBP motif co-occurrence profile in astrocytes. The heatmap is computed using Pearson correlation of co-occurrence of RBP-aligned filters in astrocyte APA events. Five RBP modules are defined based on this clustering (labeled on y- and x-axes).

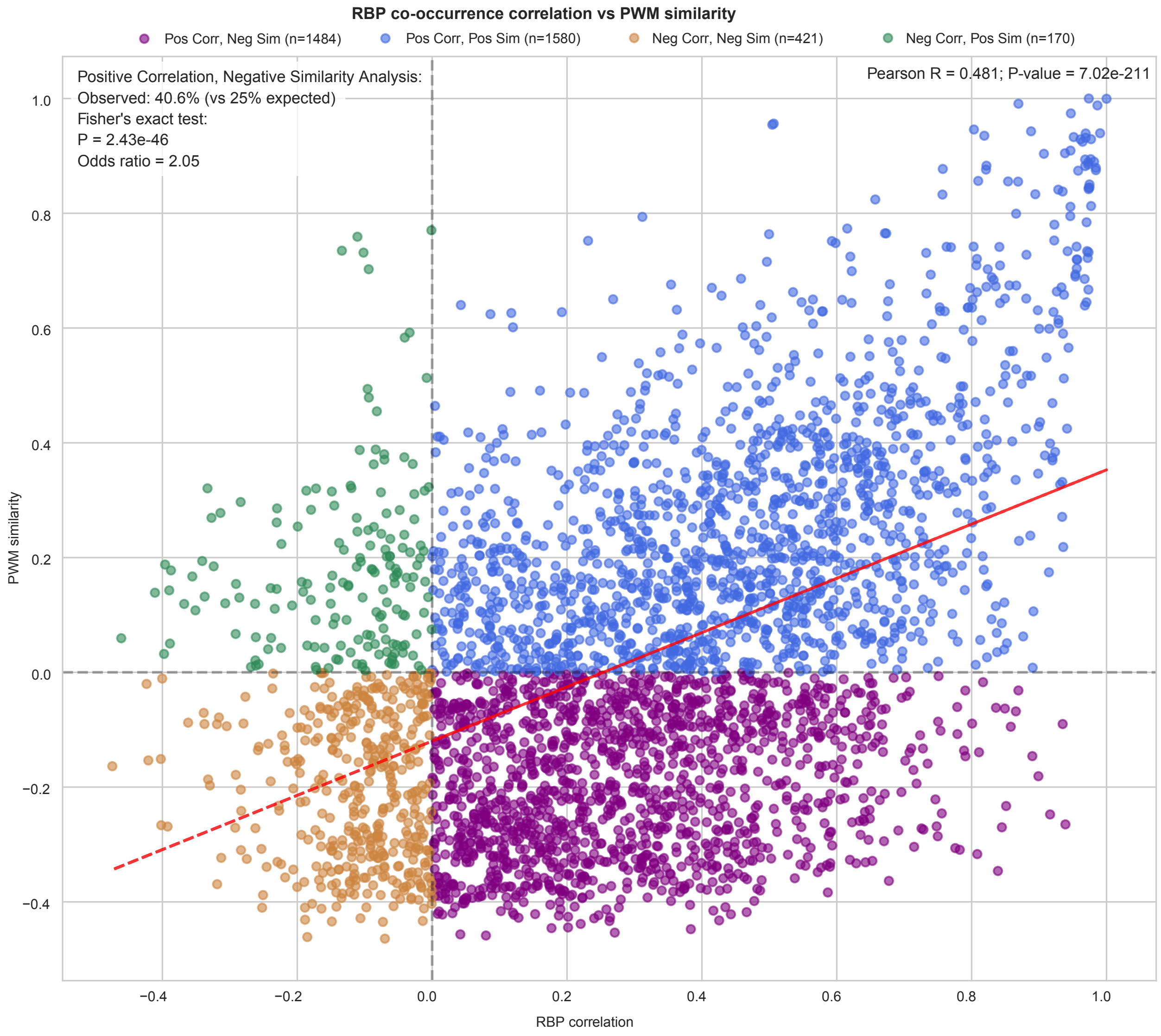


**Supplementary Fig. 26: Relationship between RBP co-occurrence and motif similarity reveals functionally related RBPs with distinct binding preferences.** Scatter plot comparing pairwise RBP co-occurrence correlation coefficients (x-axis) with their corresponding Position Weight Matrix (PWM) similarity scores (y-axis) in microglia across C9-ALS and sALS. Points are color-coded by quadrant, with purple highlighting pairs showing positive correlation but negative motif similarity (n = 1484 pairs). Statistical analysis using Fisher's exact test ($p < 2.4e\text{-}46$) confirms significant enrichment of RBP pairs exhibiting positive functional correlation despite zero or negative PWM similarity, suggesting cooperative regulation through distinct binding sites. Red dashed line represents the linear regression fit.
